# High Sensitivity Top-down Proteomics Captures Single Muscle Cell Heterogeneity in Large Proteoforms

**DOI:** 10.1101/2022.12.29.521273

**Authors:** Jake A. Melby, Kyle A. Brown, Zachery R. Gregorich, David S. Roberts, Emily A. Chapman, Lauren E. Ehlers, Zhan Gao, Eli J. Larson, Yutong Jin, Justin Lopez, Jared Hartung, Yanlong Zhu, Daojing Wang, Wei Guo, Gary M. Diffee, Ying Ge

**Author notes:** To whom correspondence should be addressed: Dr. Ying Ge, 1111 Highland Ave., WIMR II 8551, Madison, WI 53705.

## Abstract

Single-cell proteomics has emerged as a powerful method to characterize cellular phenotypic heterogeneity and the cell-specific functional networks underlying biological processes. However, significant challenges remain in single-cell proteomics for the analysis of proteoforms arising from genetic mutations, alternative splicing, and post-translational modifications. Herein, we have developed a highly sensitive functionally integrated top-down proteomics method for the comprehensive analysis of proteoforms from single cells. We applied this method to single muscle fibers (SMFs) to resolve their heterogeneous functional and proteomic properties at the single cell level. Notably, we have detected single-cell heterogeneity in large proteoforms (>200 kDa) from the SMFs. Using SMFs obtained from three functionally distinct muscles, we found fiber-to-fiber heterogeneity among the sarcomeric proteoforms which can be related to the functional heterogeneity. Importantly, we reproducibly detected multiple isoforms of myosin heavy chain (~223 kDa), a motor protein that drives muscle contraction, with high mass accuracy to enable the classification of individual fiber types. This study represents the first “single-cell” top-down proteomics analysis that captures single muscle cell heterogeneity in large proteoforms and establishes a direct relationship between sarcomeric proteoforms and muscle fiber types, highlighting the potential of top-down proteomics for uncovering the molecular underpinnings of cell-to-cell variation in complex systems.

**Significance Statement:** Single-cell technologies are revolutionizing biology and molecular medicine by allowing direct investigation of the biological variability among individual cells. Top-down proteomics is uniquely capable of dissecting biological heterogeneity at the intact protein level. Herein, we develop a highly sensitive single-cell top-down proteomics method to reveal diverse molecular variations in large proteins (>200 kDa) among individual single muscle cells. Our results both reveal and characterize the differences in protein post-translational modifications and isoform expression possible between individual muscle cells. We further integrate functional properties with proteomics and accurately measure myosin isoforms for individual muscle fiber type classification. Our study highlights the potential of top-down proteomics for understanding how single-cell protein heterogeneity contributes to cellular functions.

## Introduction

Single-cell analysis has revealed that even morphologically and genetically identical cells can differ dramatically in their functional properties, as well as how they respond to stressors and extracellular ques (1–4). Mass spectrometry (MS)-based methods for single-cell proteomic analysis hold great promise for unraveling the molecular underpinnings of cellular heterogeneity and thus are highly desired (4–12), but have lagged behind the development of single-cell genomics and transcriptomics (3, 13, 14). The major challenges for MS-based single-cell proteomics are the limited protein content of most mammalian cell types, the broad dynamic range of the proteome, and the high complexity in proteoforms arising from sequence variations and post-translational modifications (PTMs) (4, 6, 12, 15).

The two principal MS-based proteomics techniques are the peptide-centric “bottom-up” and protein-centric “top-down” approaches (16–18). Since peptides are more easily separated, ionized, and fragmented than intact proteins, bottom-up proteomics has served as the workhorse for MS-based proteomics (19–21). Nearly all MS-based single-cell proteomics studies to date employ bottom-up proteomics (8–11, 22, 23). Nevertheless, this approach suffers from intrinsic limitations, including the protein inference problem and loss of information and connectivity in mapping sequence variations and PTMs (24, 25). Top-down proteomics circumvents these limitations by analyzing intact proteins directly allowing for the identification and analysis of proteoforms—a term encompassing the myriad protein products arising from a single gene as a result of sequence variations, alternative splicing, and PTMs (25–28). Although inherently less sensitive than current bottom-up approaches due to the exponential decay in the signal-to-noise ratio with increasing molecular weight (29), top-down proteomics is ideally-suited for dissecting cellular heterogeneity at the level of proteoforms (25–28, 30–33).

In this study, we developed a high sensitivity top-down proteomics method to examine proteoform heterogeneity at the single cell level. We applied this method to single muscle fiber (SMFs), multinucleated single muscle cells with heterogeneous structural and functional properties (34–37), to establish a direct relationship between proteoforms and muscle fiber types (**Figure 1A-B**). Notably, we have detected large proteoforms (>200 kDa) and visualized the single cell heterogeneity in the SMFs. Using SMFs obtained from three functionally distinct muscle types, we found fiber-to-fiber heterogeneity in a large number of proteoforms in the sarcomere. Sarcomere is the basic contractile units in muscle, which consists of thin and thick filaments flanked serially by dense protein structures known as Z-disks (38–41). We have simultaneously characterized PTMs together with and isoforms (from different genes) from SMFs. Importantly, we detected multiple isoforms of myosin heavy chain (MyHC, 223 kDa) (36, 37, 42–45), a motor protein of thick filament playing a critical role in muscle contraction, from the SMFs with high mass accuracy and reproducibility enabling the classification of fiber type at the single-cell resolution. As illustrated, top-down proteomics of SMFs provides single cell resolution and reveals biological phenomena that are masked by bulk proteomics analysis.

**Figure 1.**
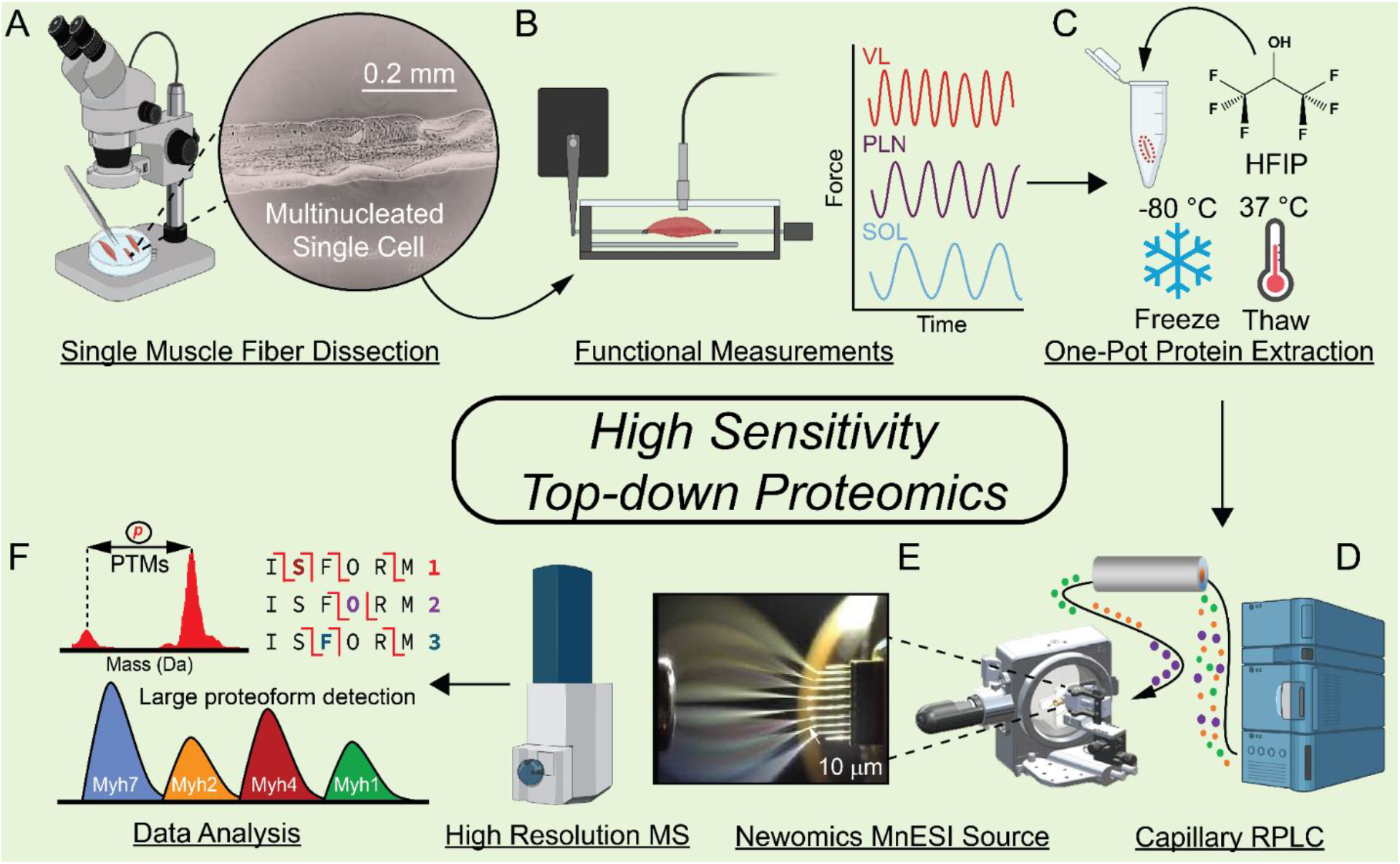
High sensitivity top-down proteomics of single muscle fibers (SMFs), multinucleated single muscle cells. **A)** SMFs were mechanically dissociated from their constituent muscles in relaxation buffer using fine point tweezers and a microscope to visualize the fibers. **B)** SMFs from each muscle were placed in a force transducer to measure their shortening velocity (n=10 fibers per muscle). **C)** A separate group of SMFs were placed directly into individual low protein binding microcentrifuge tubes for top-down proteomics measurements (n=6 fibers per muscle). Proteins were extracted from SMFs using hexafluoro-2-propanol (HFIP) and a freeze-thaw lysis. **D)** Proteins from SMF extracts were loaded onto a capillary reversed phased liquid chromatography (RPLC) column and separated based upon their hydrophobicity. **E)** Eluting proteins from SMFs were ionized with a Newomics microflow nanospray electrospray ion (MnESI) source and analyzed using a Bruker maXis II mass spectrometer. **F)** Analysis of the resultant top-down proteomics data included post-translational modification (PTM) measurements, isoform characterization using tandem mass spectrometry (MS/MS), as well as detection of large proteoforms.

## Results

### A high sensitivity top-down proteomics platform

Our high sensitivity top-down proteomics method is designed to minimize adsorptive protein losses by performing the protein extraction and follow-up sample processing in one pot using MS-compatible solvents. Subsequently, the extracted proteins are separated using high sensitivity low-flow capillary liquid chromatography (LC) coupled to a microflow multi-emitter nanoelectrospray (MnESI) source to increase ionization efficiency followed by tandem MS (MS/MS) for intact protein analysis (**Figure 1**).

Inspired by previous bottom-up single-cell proteomic studies (9, 10, 46–48), we sought to establish a one-pot sample preparation method to minimize protein losses (**Figure 1C, Figure S1**). First, we optimized the lysis buffers to effectively extract proteins using a minimal amount of tissue samples (1 mg) (**Supplemental Note 1, Figure S2**). Addition of hexafluoro-2-propanol (HFIP) is proved to be the most effective at extracting proteins from a small amount of tissue sample while maintaining MS-compatibility (**Figure S2**). Subsequently, by evaluation of different lysis techniques, we found that freeze-thaw lysis clearly performed better than sonication for protein extraction from SMFs (**Supplemental Note 2, Figure S3)**. Next, we assessed various concentrations of HFIP in the extraction solution and chose 25% HFIP which yielded the greatest protein extraction and maintained MS-compatibility (**Supplemental Note 3, Figure S4**). To determine whether freeze-thaw lysis in HFIP affected protein PTMs (49), we compared this method to our established protein extraction protocol (50, 51) (**Supplemental Note 4**). Our results have shown no discernible changes in PTMs or proteolysis resulting from freeze-thaw lysis (**Figure S5**). Thus, we have chosen a solution of 25% HFIP and a freeze-thaw cycle for cell lysis and protein extraction in this one-pot sample preparation method (**Figure 1C**).

We next developed a highly sensitive top-down LC-MS/MS method by minimizing sample dilution during RPLC and maximizing ionization efficiency. We chose a capillary RP column (Thermo MAbPac) with a narrow column inner diameter (150 μm) to minimize sample dilution during protein separation and provide robust separation afforded by μL flow rates (2 μL/min). The capillary column was coupled to a Newomics micro-flow-nanospray electrospray ionization (MnESI) source, which splits μL flow rates into 8 nanoelectrospray ionization (nanoESI) emitters substantially increasing ionization efficiency (52, 53) (**Figure S6**). The MnESI source allows for high sensitivity, throughput, and quantitative accuracy for intact protein characterization (54, 55) that resulted in a 0.5 ng detection limit of a standard protein using our method (**Figure S6**). We found that there was approximately a 4-fold gain in LC-MS sensitivity and almost twice as many fragment ions detected when using the Newomics MnESI source compared to a conventional ESI source (**Figure S7**). We combined this high sensitivity top-down proteomics method with one-pot sample preparation to achieve minimal sample losses and highly sensitive capillary LC-MS/MS for proteoforms (from the same gene) and isoforms (from different genes of the same protein family) characterization (**Figure 1E-G**).

### Integration of functional properties and top-down proteomic analysis of SMFs

Next, we applied our high sensitivity top-down proteomics approach to SMFs (multi-nucleated single muscle cells) because skeletal muscle is remarkably heterogeneous, containing a mixture of SMFs, blood vessels, nerves, and connective tissue (34–37, 41, 44) (**Figure 2A-B**); thus, methods to study contractile proteoforms at the single fiber level are highly desirable. SMFs are broadly classified as either fast-twitch or slow-twitch based on their contractile and metabolic properties (37). We chose the vastus lateralis (VL), plantaris (PLN), and soleus (SOL) muscles because VL and SOL contain predominantly fast- and slow-twitch fibers, respectively, whereas PLN contains a mixture of fast- and slow-twitch fibers (37, 56) (**Figure 1A, 2A**).

**Figure 2.**
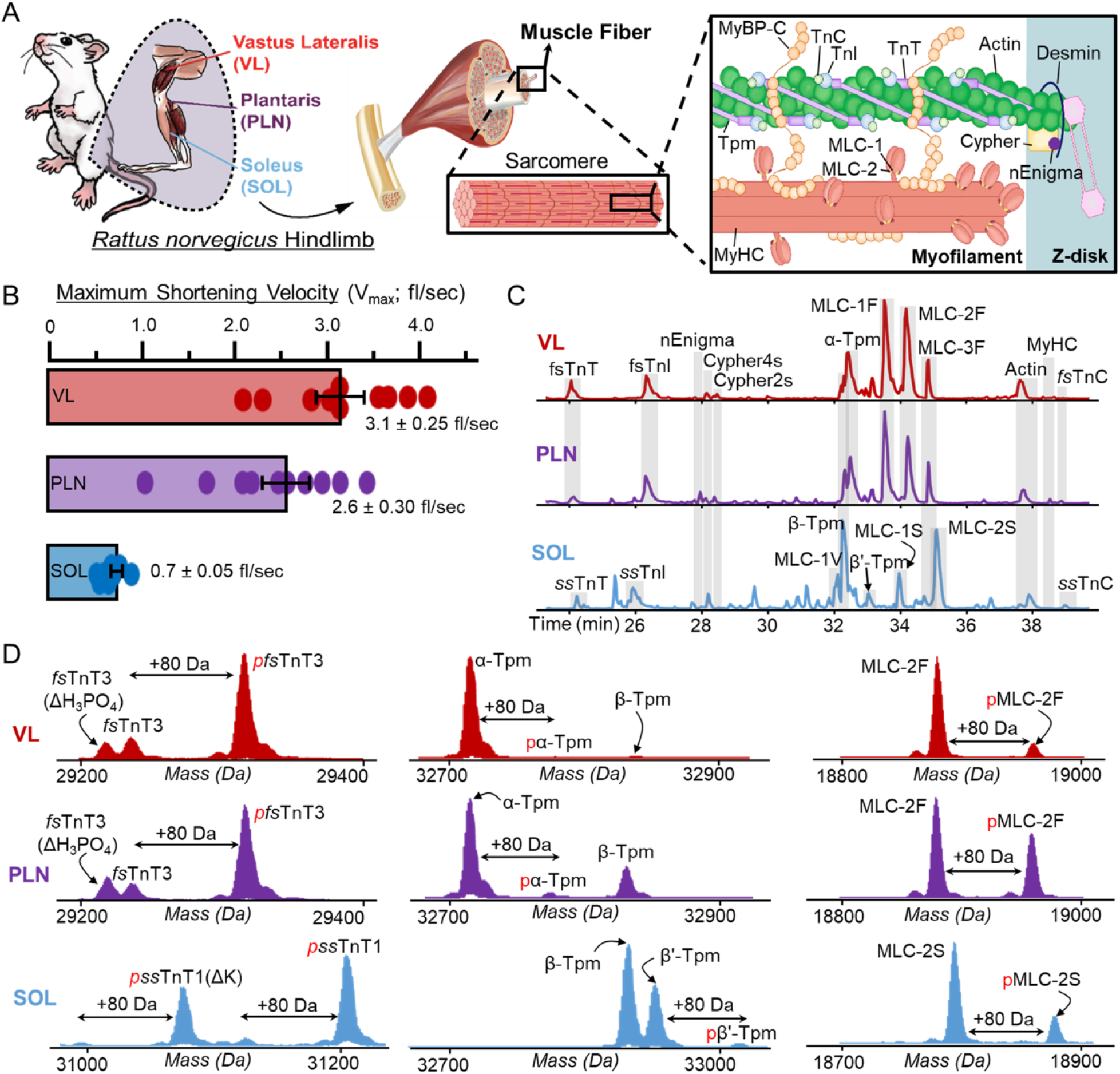
SMFs obtained from skeletal muscles have unique contractile properties and sarcomere proteoform landscapes. **A)** Schematic representation of skeletal muscle structure, which are made up of SMFs. Within SMFs are the sarcomeres, which contain the proteins necessary for contraction and relaxation including thin filament proteins (TnT, TnI, TnC, Tpm, Actin), thick filament proteins (MLC-1, MLC-2, MyHC), and Z-disk proteins (e.g. Cypher2s, Cypher 4s, nEnigma). **B)** Measurement of the maximum shortening velocity (V_max_; fiber lengths (fl)/second) of SMFs isolated from the VL, PLN, and SOL muscles (n=10 fibers per muscle). **C)** Separation and detection of major sarcomeric proteoforms detected in SMFs from VL, PLN, and SOL muscles via LC-MS/MS; base peak chromatograms (BPCs) shown (n=6 fibers per muscle). **D)** Representative deconvoluted mass spectra displaying proteoforms from VL (fast-twitch), PLN (mixed fast- and slow-twitch), and SOL (slow-twitch) single fibers. Mono-phosphorylation is indicated by red “p”, “ΔH_3_PO_4_” indicates a loss of phosphate from pfsTnT3, and “-K” indicates loss of a lysine residue from ssTnT1 or pssTnT1.

To determine the unique functional properties amongst fiber types from three muscle types (VL, PLN, SOL), SMFs from each muscle group (n=10) were dissected in a relaxation buffer that contained protease and phosphatase inhibitors, as well as antioxidants, under a confocal microscope (**Figure 1B, SI Appendix**). The average maximum shortening velocities was measured for SMFs obtained from VL (3.23 ± 0.25 fiber lengths/sec (fl/sec)), PLN (2.56 ± 0.30 fl/sec), and SOL (0.71 ± 0.05 fl/sec) muscles (**Figure 1C, 2B, SI Appendix**). These values are in agreement with those previously reported in the literature for fast- and slow-twitch muscle fibers isolated from rat muscle (57). There was a larger amount of heterogeneity in the shortening velocity measurements of SMFs from the fast-twitch VL and PLN muscles, which have three distinct fast muscle fiber types (Type IIa, IIb, and IIx), compared to the slow-twitch SOL muscle, which only has one slow muscle fiber type (**Figure 2B**).

Next, we employed high sensitivity top-down proteomics to decipher the proteoform heterogeneities that correlate with unique functional properties in SMFs from VL, PLN, and SOL (**Figure 1D–1F, Figure 2C**). The basic contractile apparatus of SMFs is the sarcomere, which consists of thin and thick filaments flanked serially by dense protein structures known as Z-disks (38–40, 58) (**Figure 2A**). Single muscle cell top-down proteomic analysis of SMFs was highly reproducible across technical replicates (**Figure S8**), allowing for the relative quantification of sarcomeric proteoforms between SMFs obtained from the VL, PLN, and SOL muscles (**Figures S9-S11**). Several key myofilament (including thin and thick filaments) proteoforms and isoforms were detected in SMFs from the VL, PLN, and SOL muscles (**Figures 2C, Table S1**). As expected, fast skeletal myofilament protein isoforms, including fast skeletal troponin complex (fsTnT, fsTnI, and fsTnC), alpha tropomyosin (α-Tpm), and fast skeletal myosin light chains (MLC-1F, MLC-2F, and MLC-3F) predominated in fibers from the VL and PLN muscles (**Figure 2C–2D, Figures S12-S18**). In contrast, slow skeletal troponin complex (ssTnT, ssTnI, and ssTnC), beta-tropomyosin (β-Tpm), and slow skeletal myosin light chains (MLC-1S, MLC-1V and MLC-2S) were detected in SMFs from SOL muscles (**Figure 2C–2D, Figures S12-S19**). Considerably, the proteomics results of the isoform distributions can be related to the unique functional properties of the SMFs from VL, PLN and SOL muscles. Other key myofilament proteins found across all samples included alpha skeletal actin (α-sActin) and MyHC isoforms (**Figures S20-S21**). Additionally, the Z-disk proteins nEnigma (40), Cypher2s, and Cypher4s were found across all fiber samples with varying relative expression (**Figures S22-S24**). These important sarcomere proteins were all identified based on their intact protein mass within a 10 ppm mass error, and identifications were further confirmed based upon MS/MS data, consistent with previous studies on skeletal muscle top-down proteomics (39, 58, 59). Notably, we were able to measure proteoforms >30 kDa such as TnT isoforms, Cypher isoforms, α-sActin (42 kDa) and MyHC isoforms (223 kDa) (**Figures S12, S20-S23**). These examples illustrated the high sensitivity of our top-down proteomics method in detecting intact sarcomeric proteoforms at the level of single muscle cells.

### Measurement of intact MyHC isoforms for fiber-type classification

Diversity of MyHC isoforms, molecular motor proteins that generate energy for force and contraction, influences the structural and functional properties of SMFs (36, 37, 42–44). MyHC isoforms are approximately 223 kDa in size and share over 80% sequence homology, making them difficult to detect and quantify using bottom-up proteomics, but each isoform corresponds to unique fiber contractile properties (36, 37, 42–44). In rat skeletal muscle, there are four major MyHC isoforms encoded by the genes *Myh1, Myh2, Myh4*, and *Myh7* (56). MyHC7 is the slow-twitch isoform that is the predominant isoform in SOL tissue (Type I), MyHC4 is a fast-twitch isoform that is the predominant isoform found in VL fibers (Type IIb), and PLN tissue mainly contains a heterogeneous mixture of the three fast-twitch MyHC isoforms (MyHC2 - Type IIa, MyHC4 - Type IIb, and MyHC1 - Type IIx) (37, 56).

We have observed the characteristic charge state envelope of a large proteoform, as evident by the myriad high charge state ions, at the later phase of 1D RPLC gradient (38 min), unexpectedly (**Figure 3A; Figure S20A**). The deconvoluted mass spectra revealed that there were four unique masses in total from the SMFs obtained from VL, PLN, and SOL, presumably corresponding to the four major MyHC isoforms found in rat (**Figure 3B; Figure S20B**). A single MyHC isoform with a mass of 222,679 ± 1 Da was detected in SMFs from SOL, which we assigned as MyHC7 (encoded by *Myh7*) due to the prevalence of Type I fibers in this muscle (**Figure 3C**). Similarly, a single MyHC isoform with a mass of 222,867 ± 2 Da was detected in SMFs from VL, which likely corresponds to MyHC4 (encoded by *Myh4*) based on the high abundance of Type IIb fibers in this muscle (**Figure 3D**).

**Figure 3.**
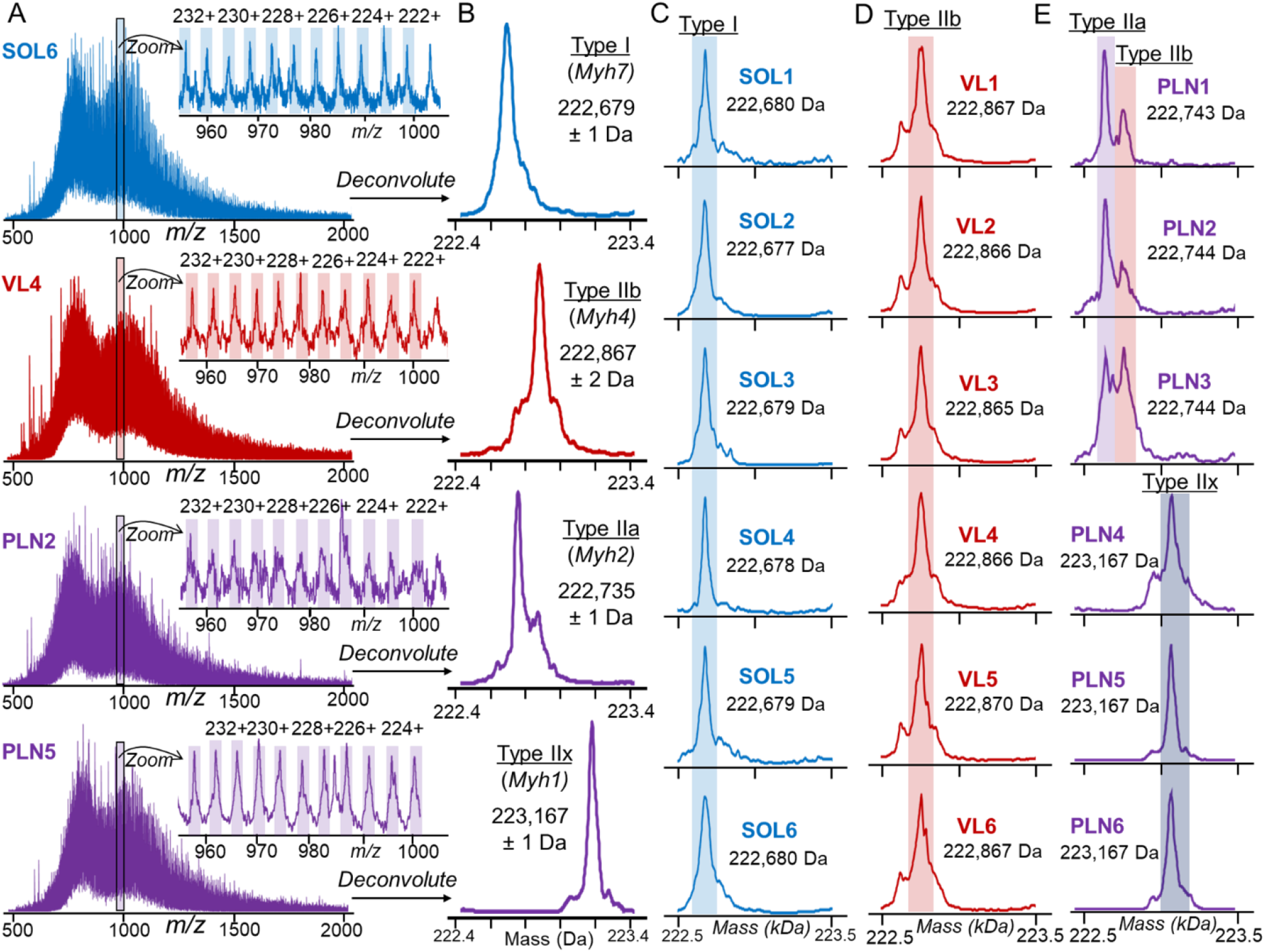
Myosin heavy chain (MyHC) isoforms detected from SMFs. **A)** Representative MS1 scans of MyHC isoforms from SMFs obtained from SOL, VL, and PLN muscles. Zoom-in of 960-1000 *m/z* region shows the highly charged ions characteristic of large proteins with high accuracy mass measurements (1-2 Da). **B)** Low resolution maximum entropy deconvolution of representative VL, PLN, and SOL SMFs reveals four distinct masses from the MS1 spectra presumably corresponding to Type I, Type IIa, Type IIb, and Type IIx MyHC isoforms. **C-E)** Deconvoluted mass spectra for all SMFs from SOL, VL, and PLN muscles (n=6 fibers per muscle). **C.** SOL, Type I; **D)** VL, Type IIb; and **E)** PLN, Type IIa, Type IIb, and Type IIx. SOL, VL, and PLN muscles (n=6 fibers per muscle). **C.** SOL, Type I; **D)** VL, Type IIa and Type IIb; and **E)** PLN, Type IIa, Type IIb, and Type IIx.

MyHC isoform expression was particularly heterogeneous in SMFs obtained from PLN muscle. Three of the SMFs obtained from PLN exhibited dual fiber characteristics, expressing both MyHC4 (Type IIb) and a second isoform with a mass of 222,735 ± 1 Da that we believe to be MyHC2 (encoded by *Myh2;* Type IIa) (**Figure 3E**). The other three SMFs obtained from PLN expressed MyHC isoforms with a mass of 223,167 ± 1 Da, which we putatively assigned as MyHC1 (Type IIx) (**Figure 3E**). Remarkably, the molecular mass measurements of MyHC isoforms were highly consistent across the fiber samples with standard deviations of 1-2 Da, enabling the classification of SMFs based upon MyHC isoform expression. The greater variability in maximum shortening velocities in fibers from PLN muscle reflects the fact that multiple MyHC isoforms are expressed in PLN fibers versus only predominantly one MyHC isoform in SOL and VL fibers (**Figure 2B, Figure 3D-E**) (56, 57). This is the first report on the detection of intact MyHC isoforms at the single muscle cell level which was made possible by our high sensitivity top-down proteomics method.

### Top-down MS analysis of isoforms together with PTMs in SMF

Both isoforms and PTMs are known to serve as key regulators for muscle contraction and relaxation (33, 59, 60). We have previously shown that top-down proteomics presents unique advantages in characterizing PTMs together with isoforms encoded by different genes from a multigene family, which often exhibit high sequence homology (39, 40, 61). Here top-down MS data provide a “bird’s eye” view of the sarcomeric proteoforms (arising from PTMs and sequence variations from a single gene) together with isoforms (resulting from different genes in a protein family) (**Figure 2D**). Many key sarcomere proteins present a diversity of isoforms in three different muscle types (VL, PLN, SOL) together with PTMs such as N-terminal di-methylation and acetylation, as well as phosphorylation and methylation (**Figure 2D, Figures S12-S24**).

Perhaps one of the best examples of using top-down proteomics to analyze PTMs together with isoforms is illustrated by the vast number of fsTnT, the Tpm-binding subunit of the troponin complex, detected in VL and PLN SMFs (**Figure 4**). The mammalian *Tnnt3* gene, which encodes fsTnT3, contains 19 exons and can theoretically generate up to 256 variants due to the alternative splicing (62). Additionally, many fsTnT isoforms share high sequence homology and are found in low abundance making their detection by bottom-up proteomics challenging. Lastly, many fsTnT isoforms are known to be post-translationally modified, such as N-terminal acetylation and phosphorylation (39, 40, 51, 58, 59), adding to the complexity of fsTnT analysis. SMFs from the VL and PLN muscles contained several highly abundant fsTnT3 proteoforms, including mono-phosphorylated fsTnT3 (pfsTnT3), fsTnT3, and pfsTnT3 with the loss of phosphate (ΔH3PO4) proteoforms (**Figure 4A**). The fsTnT3 total level of phosphorylation (Ptotal) values (**Figure 4C, Figure S12**) were highly consistent across all of the SMF samples obtained from VL and PLN muscles, as well as the relative expression of fsTnT3 (**Figure 4D**). Appreciably, when we zoomed in on the VL and PLN deconvoluted spectra twenty-fold, there were several low-abundance and fiber-specific fsTnT isoforms and proteoforms revealed (**Figure 4B**). We detected fsTnT4 proteoforms from all VL and PLN, but the Ptotal values were highly variable, particularly for the SMFs obtained from PLN muscle (**Figure 4C**). Besides heterogeneity in fsTnT4 proteoform expression, we found that some of the SMFs obtained from PLN contained unique fsTnT isoforms, including fsTnT9 and fsTnT10. These observations were further exemplified by normalizing all of the low abundance proteoforms to their highest intensity peak (**Figure S25**).

**Figure 4.**
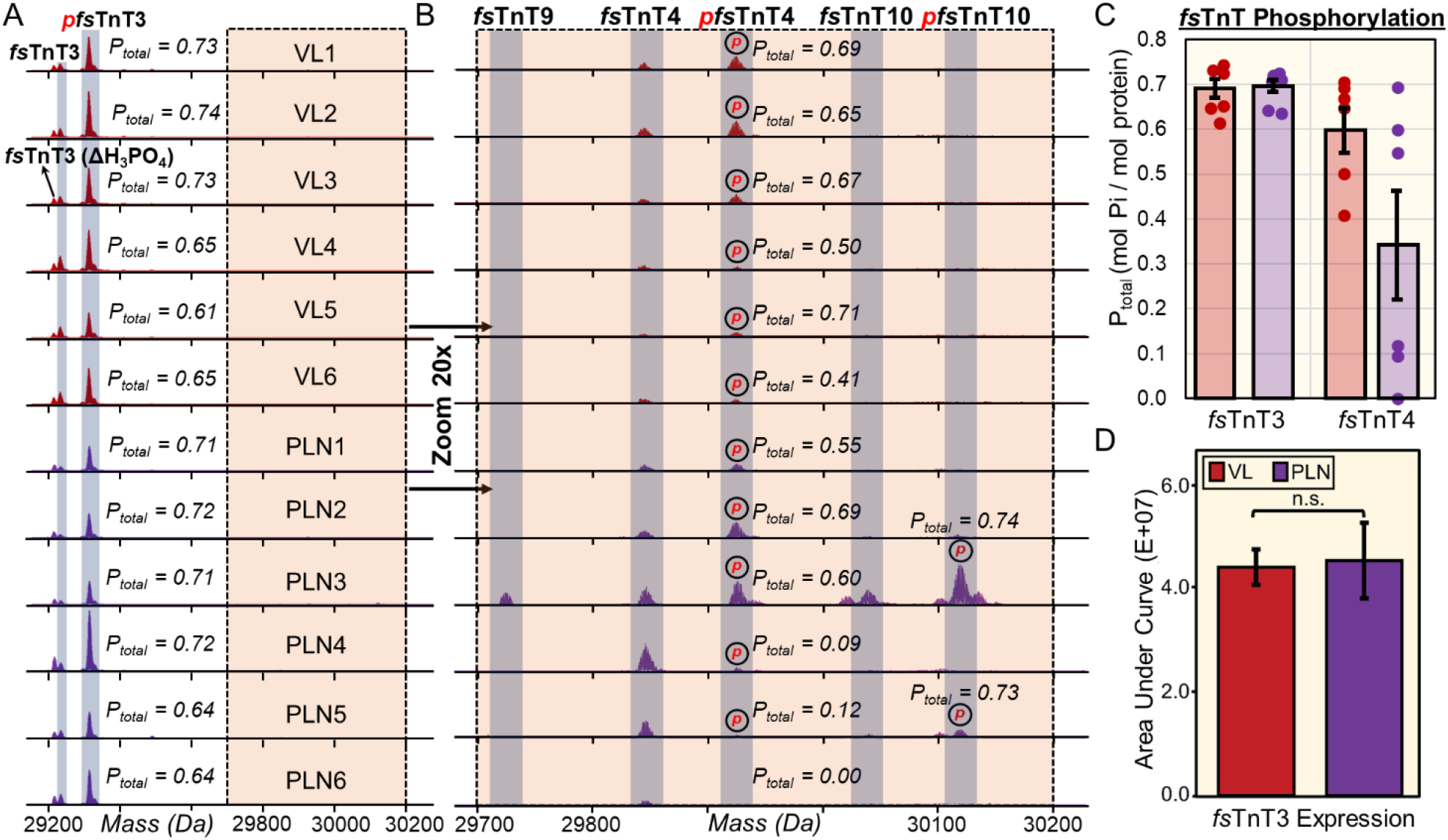
Altered phosphorylation of low-abundance fast skeletal troponin T (fsTnT) isoforms and proteoforms from fiber-to-fiber. **A)** Representative deconvoluted mass spectra of fast skeletal troponin 3 (*fs*TnT3) from SMFs isolated from fast-twitch VL (red) and PLN (purple) muscles. All the spectra are normalized to 1,000,000 intensity units. Mono-phosphorylation is denoted with red “p”; “ΔH_3_PO_4_” indicates phosphate loss from pfsTnT3. **B)** 20x magnitude zoom-in on 29,700 Da – 30,200 Da region of spectra reveals several low-abundance fsTnT isoforms (*fs*TnT4, *fs*TnT9, *fs*TnT10) in SMFs from fast-twitch VL (red) and PLN (purple) muscles. All the spectra are normalized to 50,000 intensity units. Mono-phosphorylation is denoted with red “p”. **C)** Total phosphorylation (P_tot_) calculated as mol Pi/mol protein for *fs*TnT3 and *fs*TnT4 from each fiber (n=6). **D)** Extracted ion chromatograms (EICs; top 5 most abundant ions) of *fs*TnT3 were made and the area under the curve was integrated to calculate the *fs*TnT3 expression. Groups were considered statistically different at p < 0.05; “n.s.” indicates statistically not significant by paired student t test.

Other examples of myofilament isoforms together with PTMs include thin filament protein α- and β-Tpm isoforms (**Figure 2D, Figure S15**), as well as fast- and slow-twitch MLC isoforms (**Figures S16-19**). Importantly, the top-down approach enabled the direct differentiation of two Tpm isoforms, β-Tpm and β’-Tpm (53), with only 26 Da mass difference (**Figure 2D, Figure S15**). As illustrated, high sensitivity top-down proteomics is uniquely suited to provide a “bird’s eye view” of the myriad isoforms together with PTMs from single muscle cells.

### Top-down proteomics captures single cell heterogeneity in SMFs at the proteoform level

The high sensitivity top-down proteomics of SMFs here clearly reveal single cell proteoform heterogeneity (**Figure 5, Figure S15, 18, 19, 24**). For example, the fast and slow isoforms of MLC-2 (MLC-2F and MLC-2S) displayed a remarkable amount of proteoform heterogeneity across the fiber samples (**Figure 5**). The high sensitivity top-down approach detected MLC-2F and pMLC-2F in the SMFs obtained from VL and PLN, whereas MLC-2S and pMLC-2S were detected in SMFs obtained from SOL (**Figure 5A-C, Figure S17**). MLC-2S and MLC-2F have distinct functions yet share similar sequences differing by only 88.9 Da; even so, the isoforms can be separated by our online LC-MS/MS method chromatographically (**Figure 5D**). The deconvoluted spectra clearly show that there are fiber-to-fiber differences in the total phosphorylation (Ptotal) of MLC-2F and MLC-2S (**Figure 5E**). On average, the MLC-2 Ptotal values for SMFs obtained from VL, PLN, and SOL were 0.15 ± 0.02, 0.34 ± 0.07, and 0.17 ± 0.08, respectively (**Figure 5E**). The SMFs obtained from PLN and SOL displayed a remarkable degree of phosphorylation heterogeneity with MLC-2 Ptotal values ranging from 0.21 to 0.39 for SMFs obtained from PLN and 0.05 to 0.29 for SMFs obtained from SOL. Previously, top-down proteomics of bulk skeletal muscle tissue samples reports an ensemble measurement of hundreds of fibers (39, 40, 58), which averages all the differences among diverse single muscle cells.

**Figure 5.**
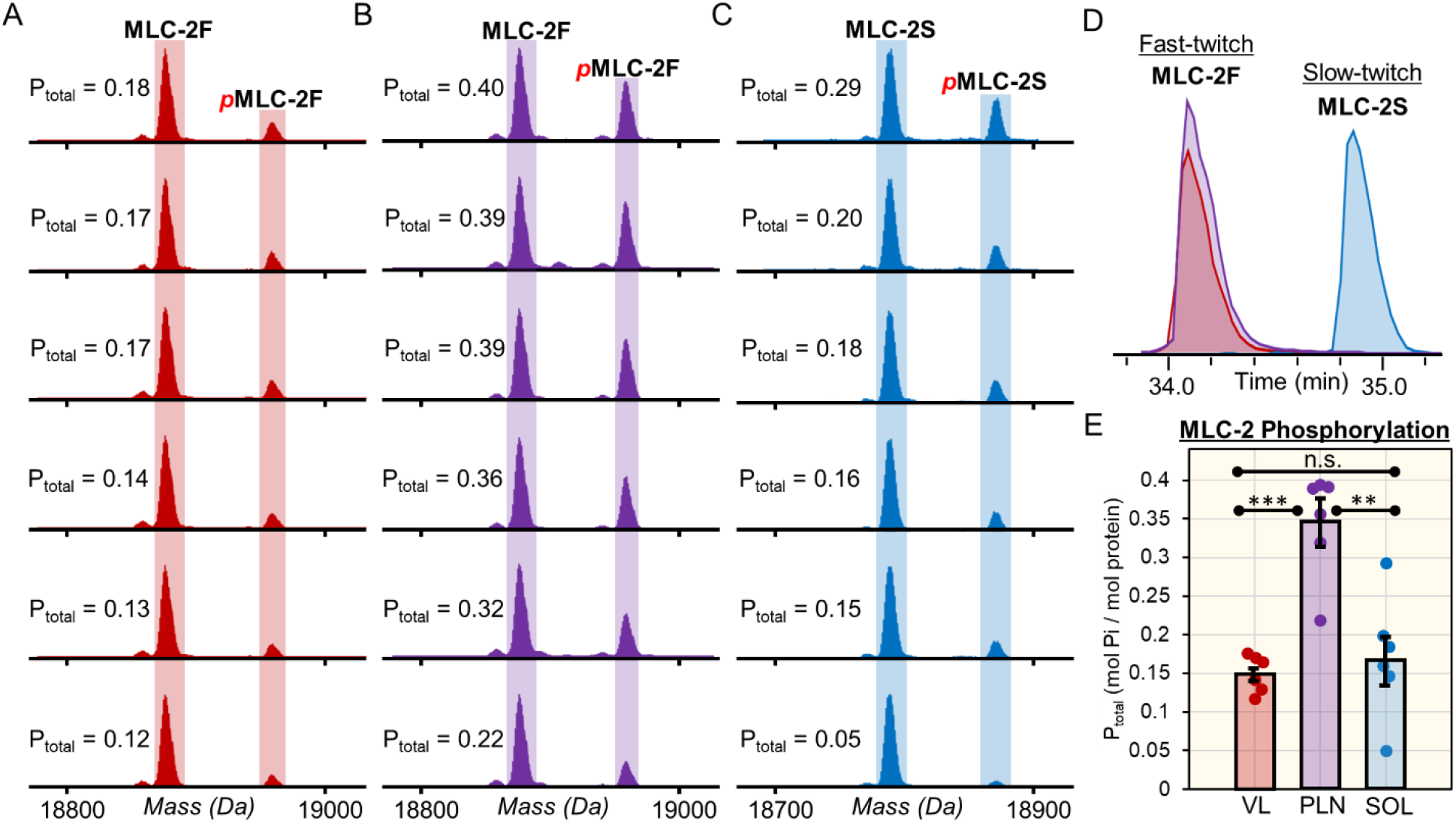
Proteoform heterogeneity in myosin light chain 2 (MLC-2) isoforms across different SMFs. **A)** EICs (top 5 most abundant ions) for fast and slow isoforms of myosin light chain 2 (MLC-2F and MLC-2S, respectively) obtained from SMFs from VL, PLN, and SOL muscles. **B-C)** Deconvoluted mass spectra of MLC-2F from SMFs from fast-twitch VL and PLN muscles. **D)** Deconvoluted mass spectra of MLC-2S from SMFs from slow-twitch SOL muscles. **E)** Total phosphorylation (P_tot_) calculated by mol Pi/mol protein for MLC-2F and MLC-2S for SMFs from VL, PLN, and SOL muscles (n=6 fibers per muscle). Groups were considered statistically different at p < 0.05; “n.s.” indicates statistically not significant by paired student t test.

Moreover, fiber-to-fiber differences have also present in other sarcomeric proteoforms. For instance, both SMFs obtained from VL and PLN expressed the slow-twitch associated isoform, β-Tpm, however SMFs obtained from PLN had a higher abundance (**Figure S15**). Furthermore, we found the MLC-3F was in all the SMFs obtained from SOL but at a much lower abundance than found in the SMFs obtained from PLN (**Figure S18**). Additionally, one SMF obtained from PLN (PLN1) expressed slow-twitch MLC-1V, whereas none of the other SMFs obtained from PLN contained this proteoform (**Figure S19**). Lastly, the ratio of MLC-1F to MLC-3F isoform expression, thick filament proteins involved in the structural stability of MyHC, was found to be highly heterogeneous across the SMFs obtained from VL and PLN muscles (**Figure S26**), implicating the structural role of isoform abundance in myosin stability. The myofilament proteoform heterogeneity observed in our samples is uniquely captured by performing top-down proteomics at the single fiber level.

### Effective sequence characterization by top-down online LC-MS/MS in SMF

The capability of the top-down online LC-MS/MS for unambiguous proteoform identification and characterization at the single muscle level was demonstrated using the myriad MLC isoforms (**Figure 6**) found in single muscle cells. MLCs are associated with MyHCs and essential for physiological speeds of shortening skeletal muscle (43). SMFs from SOL contained two MLC-1 isoforms (MLC-1S and MLC-1V) and SMFs from PLN and VL contained one MLC-1 isoform (MLC-1F) (**Figure S16, S19**). Alignment of the sequences of MLC-1F, MLC-1S, and MLC-1V revealed the high homology of these isoforms (**Figure 6A**). The LC-MS/MS method effectively separates all three of these isoforms from one another (**Figure 6B**) and measures MLC-1 isoform spectra with high mass accuracy and isotopic resolution (**Figure 6C**) prior to CAD MS/MS, which generates several unique *b* and *y* ions that are characteristic of the MLC-1 isoform sequences (**Figure 6D**). Top-down proteomics enabled the confident characterization of each MLC-1 isoforms as well as their N-terminal modifications, such as acetylation and di-methylation (**Figure 6E**). In addition to MLC-1, we characterized the fast- and slow-twitch isoforms of MLC-2 (**Figure S27**). SMFs from SOL contained MLC-2S whereas SMFs from VL and PLN contained MLC-2F (**Figure S17**). Lastly, we were able to characterize MLC-3F, which was found in every SMF sample (**Figure S18**), with our online LC-MS/MS method (**Figure S28**). Significantly, our high sensitivity top-down proteomics approach is able to unambiguously characterize the entire MLC family including a large variety of isoforms.

**Figure 6.**
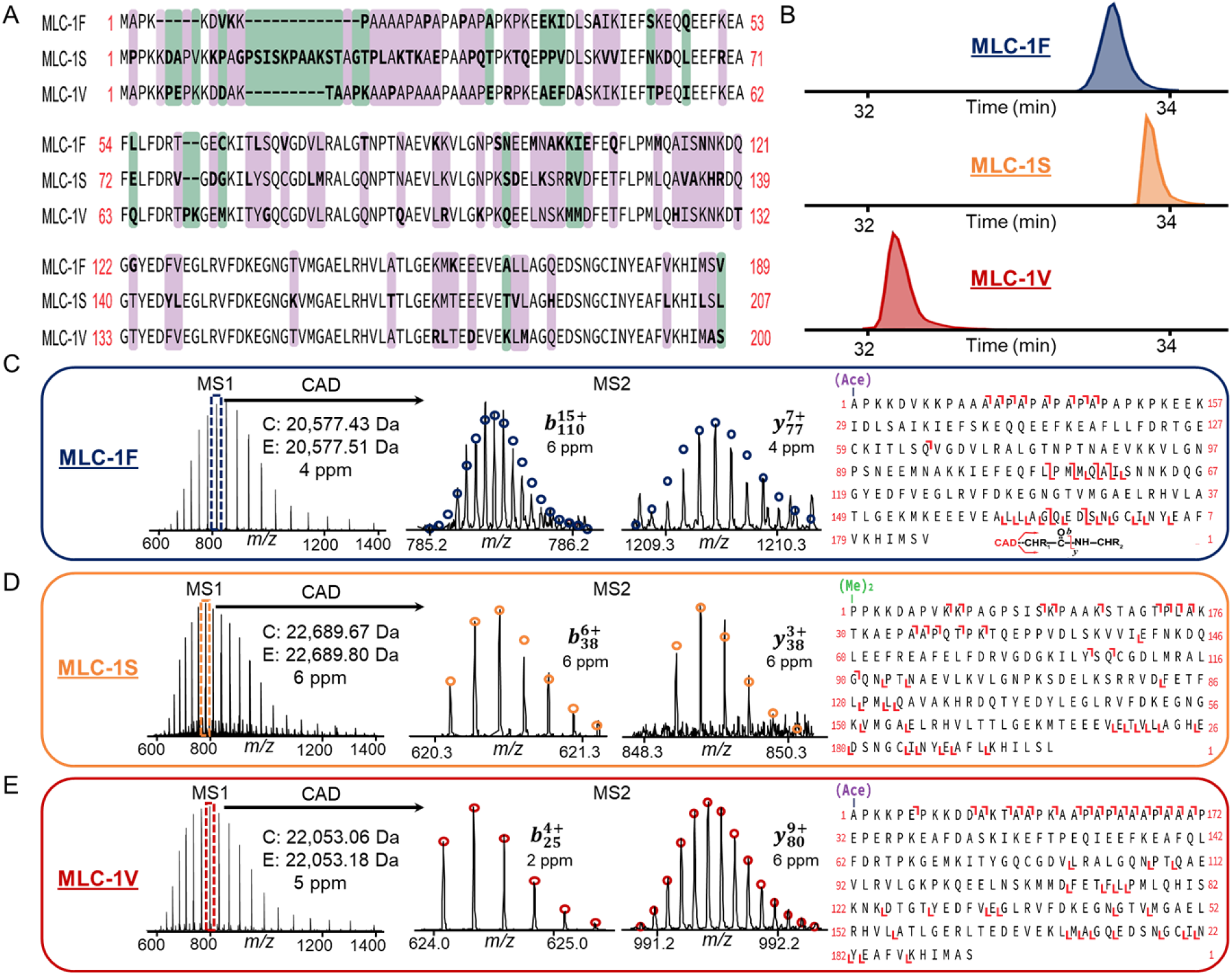
Top-down MS characterization of myosin light chain 1 (MLC-1) isoforms from SMFs. **A)** Sequence alignment of the fast, slow, and ventricular isoforms of myosin light chain 1 (MLC-1F, MLC-1S, and MLC-1V, respectively). Purple indicates no residues shared in common between the isoforms, green indicates at least one residue is shared between the isoforms, and no color indicates sequence homology between the isoforms. **B)** Representative extracted ion chromatograms (EICs) for MLC-1 isoforms. The EICs of MLC-1F, as well as MLC-1S and MLC-1V, are from SMFs from VL and SOL muscles, respectively. **C-E)** Online LC–MS/MS of MLC-1 isoforms. Precursor ions were selected for online collisionally activated dissociation resulting in signature *b* and *y* ions that are characteristic of MLC-1F **(C)**, MLC-1S **(D)**, and MLC-1V **(E)**. “(Ace)-”denotes N^α^-acetylation, “(Me)_2_-” denotes N^α^-dimethylation. Circles represent the theoretical isotopic abundance distribution corresponding to the assigned mass and based on the averagine model.

Next, we have showed detailed characterization of several key sarcomeric proteins including α-sActin, TnI, TnT, Tpm, and Cypher isoforms (**Figure S29-S33**). Importantly, we have demonstrated the capability of this high sensitivity top-down proteomics platform for characterization of large proteoform (>30 kDa) from single muscle cells at the chromatographic time scale. For example, online LC-MS/MS was successfully performed on a critical thin filament protein, α-sActin (Calc’d: 41,845.83 Da, Expt’l: 41,846.03 Da) resulting in multiple high-quality *b* and y ions for sequence characterization (**Figure S29**). Top-down MS unambiguously characterized highly similar isoforms such as the fast- and slow-twitch isoforms of TnI, TnT and Tpm (**Figure S30-S32**). Accurate mass measurements were obtained for the ssTnT (Calc’d: 31,187.07 Da, Expt’l: 31,187.18 Da) and fsTnT3 (Calc’d: 29,299.33 Da, Expt’l: 29,299.45 Da) together with online MS/MS for sequence characterization; similarly LC-MS/MS data were obtained for β-Tpm (Calc’d: 32,858.56 Da, Expt’l: 32,858.68 Da) and α-Tpm (Calc’d: 32,702.68 Da, Expt’l: 32702.87 Da). Moreover, our online LC-MS/MS method allowed for characterization of the Z-disk proteins Cypher2s (Calc’d: 31,379.00 Da, Expt’l: 31,319.10 Da) and Cypher4s (Calc’d: 30879.76 Da, Expt’l: 30879.96 Da), which share highly similar sequences (98.3% sequence homology) (**Figure S33**).

Additionally, we searched the MS/MS data obtained from a representative VL, PLN, and SOL fiber and performed Gene Ontology analysis (**SI Appendix**). The results showcase many processes related to muscle contraction and mitochondrial functions found across all fiber samples, as well as nuclear proteins, such as histones, as well as mitochondrial proteins (**Figure S34, Table S2-4**). Top-down proteomics provides a robust platform for the unambiguous characterization of SMF proteins.

## Discussion

Single-cell technologies are revolutionizing biology and molecular medicine by elucidating cellular heterogeneity (1–4). Single-cell proteomics has high promise to reveal cellular phenotypic heterogeneity and cell-specific functional networks underlying biological processes by reliable and unbiased measurement of proteins (5–12). However significant challenges remain in single-cell proteomics to realize its potential in characterization of PTMs and sequence variations (3, 4, 13, 14). Bottom-up MS-based proteomics has made great progress towards the analysis of proteomes from single cells (8–11, 22, 23, 46–48), yet this approach infers protein identification and quantification from peptides losing the critical information provided by proteoforms during the digestion process, which is suboptimal for analysis of PTMs and sequence variants (24, 63). In contrast, top-down proteomics, analyzing whole proteins without digestion, is ideally suited for comprehensive analysis of proteoforms providing a bird’s eye view of all PTMs together with alternative splicing and genetic mutations (25–28, 30–33). Nevertheless, top-down proteomics is inherently less sensitive compared to bottom-up proteomics, due to the exponential decay in signal-to-noise as a function of molecular weight, lower resolution intact protein separations, and the adsorptive nature of proteins (29, 33). A continual and major challenge in the top-down proteomics field is the requirement for relatively large amounts of starting sample (often several micrograms of total protein or millions of cells) and, thus, there has been a push to improve sensitivity (39, 64, 65). Overcoming this limitation is particularly important for applying top-down proteomics to biological applications with limited sample amounts such as clinical biopsy samples and single mammalian cells (33).

To address this challenge, we have developed a high sensitivity top-down proteomics method including a one-pot sample processing to reduce protein losses, followed by high sensitivity capillary LC-MS/MS using low flow capillary RPLC and the MnESI source with detection limit as low as 0.5 ng (**Figure 1, Figure S7**). This is partly motivated by previous bottom-up single cell proteomics studies which have shared a common theme of minimizing sample handling to avoid adsorptive losses, incorporating highly sensitive protein separation methods, and exploiting ultrahigh-resolution MS instrumentations (8–11, 22, 23, 46–48). We have shown the effective separation of large proteins and detailed characterization of proteoforms by online LC-MS/MS with high sensitivity and reproducibility (**Figure 2, Figure S8-11**).

We have applied the high-sensitivity top-down proteomics method to skeletal muscle SMFs (multi-nucleated single cells) because 1) skeletal muscle fibers are remarkably heterogeneous at the molecular level and, thus, provide a “proteoform-rich” proteome for top-down study (34–37); 2) the fusion of skeletal muscle precursor cells, known as myoblasts, during development produces relatively large cells with greater protein content than most other somatic cell types (36, 37); and 3) The structural and functional diversity of skeletal muscle is reflected in its basic contractile unit, the sarcomere which contains a variety of isoforms from multi-gene families together with PTMs making it an ideal system to leverage the unique capability provided by our top-down approach (39, 40, 43, 58, 66). As demonstrated, the single cell top-down proteomics analysis provides a “bird’s eye” view of protein isoforms together with PTMs in conjunction with functional measurements and captures single muscle cell heterogeneity at the proteoform level (**Figure 2B–2D**).

Skeletal muscle fibers have typically been classified as either slow/type I and fast/type II based on their physical, morphological, histological, and biochemical characteristics (44). Previous studies have shown that muscle fiber classification, and thereby function, is largely defined by which MyHC isoform expressed within the fiber (i.e. Type I, IIa, IIb, or IIx) (36, 37, 42–45). In rat, there are four MyHC isoforms expressed; fibers may express only one of these isoforms (“pure”) or may express more than one isoform (“hybrid”) (37, 56). Variances in force generation have been linked with altered expression of MyHC isoforms demonstrating that SMF contractile performance and proteoform landscape forms a continuum (36, 43). For the first time, we have reported the detection of MyHC isoforms (~223 kDa) in SMF with single-cell resolution enabled by the high sensitivity top-down proteomics method (**Figure 3, Figure S20**). Although several MyHC isoforms have been studied from SMFs using bottom-up proteomics (34, 35, 67) or middle-down MS (68), intact MyHC isoforms have never been detected at the whole protein level in SMFs. The intact mass measurements of MyHC (~223 kDa) were highly consistent across the fiber samples with standard deviations of 1-2 Da allowing for accurate fiber type classification based on MyHC isoform expression. To connect the MyHC proteomics data to the functional measurements, the fast-twitch fibers (VL and PLN) had a greater maximum shortening velocity than slow-twitch fibers (SOL) (**Figure 2B**), which is strongly correlated with MyHC isoform expressions (**Figure 3**). Importantly, there was greater variance in the VL and PLN fiber functional measurements because multiple MyHC isoforms may be expressed in these fibers.

Our top-down proteomics data illustrates the heterogeneous nature of SMFs and captures single muscle cell heterogeneity at the proteoform level. Heterogeneity in MyHC isoform expression was observed, particularly for the fast-twitch PLN muscle where multiple MyHC isoforms were detected (Type IIa, IIb, IIx) (**Figure 3**). Additionally, we found that the expression of MLC-1F and MLC-3F varied from fast-twitch fiber-to-fiber (**Figure S26**). This heterogeneity in thick filament proteins is noteworthy because both MyHC and MLC isoforms determine the maximum shortening velocity of SMFs (43, 66). In fact, the functional data demonstrated extensive heterogeneity in shortening velocity, particularly in the fast-twitch fibers, which can be related to the proteomics heterogeneity (**Figure 2B**). In addition, low abundance fsTnT isoforms together with PTMs as well as MLC-2 proteoforms demonstrated fiber-to-fiber differences from the bird’s eye view provided by top-down proteomics (**Figure 4, 5**). Besides fsTnT and MLC-2 proteoform heterogeneity, we demonstrate that several other proteins, including both myofilament and Z-disk proteins (**Figure S15-S30**), display varying degrees of proteoform heterogeneity as uniquely enabled by our top-down approach. Our results clearly demonstrate that there are fiber-to-fiber differences in proteoforms, whereas in bulk tissue analysis the results represent an ensemble measurement of hundreds of fibers in addition to other components found in skeletal muscle.

Notably, in this study, we have been able to detect large proteoforms (>200 kDa) from a single muscle cell. Although a previous study by Zhu et al. has used the nanoPOTs platform for the extraction and sampling of intact proteins from low numbers of HeLa cells, only small proteoforms (<15 kDa) were detected (69). Recently, a capillary electrophoresis coupled to MS/MS (CE-MS/MS) approach has been used for analysis of proteins from single-cells, nonetheless, the detected proteoforms were almost exclusively below 15 kDa with no proteins greater than 30 kDa (70). Top-down MS analysis of large proteins is challenging due to an exponential decay in S/N with increasing molecular weight (MW) as well as co-elution with low MW and high abundance proteins in a mixture (29, 33, 65). Typically, size-based fractionation prior to MS analysis is essential for detection of large proteins. We have previously developed a top-down 2D-LC method including serial size exclusion chromatography (sSEC) to enrich high MW proteins with reverse phase chromatography (RPC) and high-resolution top-down MS for detection of large proteins (65, 71). For the first time, we have detected large proteins (>30 kDa) using a simple 1D capillary RPLC top-down MS from single muscle cells and reveal heterogeneity at the level of proteoforms.

## Conclusion

Here, we developed a high sensitivity top-down proteomics method that combined one-pot sample preparation and highly sensitive capillary LC-MS/MS. We have shown the effective separation of proteins and detailed characterization of large proteoforms on the chromatographic time scale with high sensitivity and reproducibility. This method was applied to SMFs (multi-nucleated single cells) with remarkable functional and proteomics heterogeneity. We have integrated functional measurements and top-down proteomic analysis of SMFs and captured single muscle cell heterogeneity at the proteoform level. Using SMFs obtained from three functionally distinct muscles, we found fiber-to-fiber heterogeneity among the sarcomeric proteoforms and striking heterogeneity in large proteoforms (>200 kDa). Moreover, this study underlined the unique capability of top-down proteomics in simultaneous characterization of PTMs together with a variety of isoforms from multi-gene families in the sarcomere. Impressively, we detected multiple distinct MyHC isoforms (>220 kDa) from the SMFs with high mass accuracy and robustness which enables the classification of fiber type at the single-cell level. This represents the first report on the single cell resolution of large proteoforms, highlighting the potential of single cell top-down proteomics in advancing our understanding of phenotypic heterogeneity and functional diversity among individual cells toward precision medicine. We envision this high-sensitivity top-down proteomics method developed here can be extended to other applications requiring high sensitivity.

## Materials and Methods

Detailed materials and methods are outlined in SI Methods.

### Reagents and chemicals

All reagents were purchased from Sigma-Aldrich, Inc. unless otherwise noted. HPLC-grade water and acetonitrile were purchased from Fisher Scientific. Hexafluro-2-propanol (HFIP) was purchased from Alfa Aesar. Protease inhibitor cocktail were purchased from Thermo Fisher Scientific. Amicon, 0.5 mL cellulose centrifugal filters with a 10 kDa molecular weight cutoff (MWCF) were purchased from MilliporeSigma.

### Animals

Male Fischer 344 Norway F1 hybrid (F344BN) rats (aged 6-months) were obtained from the University of Wisconsin-Madison School of Veterinary Medicine. The rats were given access to food and water *ad libitum* and individually housed in clear plastic cages on a 12 h/12 h light/dark cycle. All procedures involving animals were carried out following the recommendations in the Guide for the Care and Use of Laboratory Animals published by the National Institutes of Health and were approved by the Institutional Animal Care and Use Committee of the University of Wisconsin-Madison.

### Relaxation solution

The relaxation solution used for the isolation of SMFs was similar to that described previously (72) (2 mM EGTA, 100 mM KCl, 10 mM imidazole, 4 mM ATP, 5 mM MgCl2) with minor modifications to accommodate top-down proteomics studies, including the addition of anti-oxidants (5 mM tris(2-carboxyethyl)phosphine), as well as protease (2 mM phenylmethylsulfonyl fluoride) and phosphatase inhibitors (2 mM sodium vanadate and 10 mM β-glycerophosphate).

### Isolation of SMFs

Bundles of ~50 fibers were dissected from the VL, PLN, and SOL muscles and placed in a petri dish containing relaxation solution under a confocal microscope. For each experiment, an individual fiber (~1.5–2.5 mm in length) was pulled from the end of the bundle using fine point tweezers and placed in a labelled low protein bind microcentrifuge tube. The location of the fiber in the tube was outlined with marker prior to freezing at −80 °C until the day of the experiment. In total, there were 10 fibers per tissue type used for mechanical measurements and 6 fibers per tissue type used for top-down proteomics analysis. All fibers were isolated from the muscles of a single rat.

### Shortening velocity measurements of SMFs obtained from VL, PLN, and SOL

The experimental technique for performing contractile measurements on skeletal muscle fibers has been described previously (58, 72). Briefly, the fiber segments were attached between a capacitance-gauge transducer (Model 403, sensitivity of 20 mV/mg and resonant frequency 600 Hz; Aurora Scientific) and a DC torque motor (Model 308; Aurora Scientific). Length changes during contractile measurements were introduced at one end of the preparation driven by voltage commands from a PC via a 16 bit D/A converter.

The velocity of unloaded shortening (Vo) was measured using the slack-test method described previously by Edman (73). With the fiber length at L, the fiber was transferred to pCa 4.5 solution and steady force was allowed to develop. The fiber was then quickly released to a shorter length, resulting in an immediate tension decline, followed by a gradual recovery of tension. The time interval before tension was redeveloped was plotted against the length of release for several different releases, and the slope of this line was calculated as the velocity of unloaded shortening. All velocity values were converted from millimeters per second to fiber lengths per second by dividing by L.

### Extraction of sarcomeric proteins from SMFs

SMFs from the VL, PLN, and SOL muscles were thawed on ice the day of LC-MS/MS experiments. To minimize artificial protein modifications and oxidation, protein extraction was performed at 4 °C. Samples were washed with 40 μL of 150 mM ammonium acetate to remove any remaining relaxation buffer by pipetting the solution over the fiber such that the fiber remained on the wall of the tube. Samples were briefly centrifuged (1,100 × *g*, 1 min, 4 °C) and the solution was removed. The fiber was suspended in 20 μL of protein extraction solution (25% HFIP, 10 mM L-methionine, 1× HALT protease inhibitor cocktail, and 1× phosphatase inhibitor cocktail A, pH 7.5). The samples were incubated for 15 min on ice followed by addition of 20 μL of mobile phase A (MPA; 0.1% formic acid (FA) in water) and a freeze-thaw cycle (incubation at −80 °C for 5 min followed by incubation for 1 min at 37 °C). The freeze-thaw cycle was repeated two additional times for a total of three cycles with mixing by gentle agitation between cycles. Samples were centrifuged (21,000 × *g*, 15 min, 4 °C). The fiber extracts were desalted prior to LC-MS/MS analysis using a 10 kDa MWCF and buffer exchanged into MPA concentrating to a final volume of 35 μL. 10 μL of the desalted extracts were used for a Bradford protein assay to estimate the protein concentration. The remaining 25 μL was transferred to a HPLC vial.

### Online LC-MS/MS analysis of SMF extracts

LC-MS/MS analysis was carried out using a NanoAcquity ultra-high pressure LC system (Waters) coupled to a high-resolution maXis II quadrupole time-of-flight mass spectrometer (Bruker Daltonics). A microflow-nanospray electrospray ionization source (MnESI) equipped with multinozzle M3 emitters (8 nozzles, 10 μm ID) provided by Newomics Inc. (Berkeley, California) was used for nanoESI. Approximately 100 ng of total sarcomeric protein extract (per sample) was separated using a MAbPac™ Capillary Reversed Phase HPLC Column (Thermo Scientific, 1500 Å pore size, 4-μm particle size, 150-μm inner diameter). SMF proteoforms were eluted using a linear 60 min gradient going from 10 to 95% mobile phase B (MPB; 0.1% FA in acetonitrile) at a constant flow rate of 2 μL/min. Data-dependent LC-MS/MS was performed on single fiber protein extracts. The three most intense ions in each mass spectrum were selected and fragmented by CAD with a scan rate of 2 Hz from 200-3000 *m/z*. The isolation window for online AutoMS/MS CAD was 10 m/z. The collision DC bias was set from 20 to 40 eV for CAD with nitrogen as the collision gas. All singly charged ions were excluded for fragmentation and dynamic exclusion was enabled after the collection of 4 of the same precursor ions to prevent fragmentation of the same species.

### Data analysis

DataAnalysis (Version 4.3; Bruker Daltonics) software was used to process and analyze the LC-MS data. The resolving power for maximum entropy deconvolution was set to 70,000 for proteins that were isotopically resolved and 10,000 for proteins that were not isotopically resolved. Mono-isotopic masses are reported for all isotopically resolved MS and MS/MS data using the Sophisticated Numerical Annotation Procedure (SNAP) peak-picking algorithm. Most abundant masses are reported for all non-isotopically resolved MS data using the Sum Peak algorithm. The total protein phosphorylation (Ptotal) for each phosphorylated protein was calculated using the following equation:

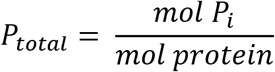

The top five most abundant charge state ions of the sarcomeric proteins of interest were used to produce extracted ion chromatograms and the area under the curve was integrated to determine the relative abundance for each protein. Tandem mass spectra were exported from DataAnalysis software and analyzed using MASH Explorer (Version 2.0) (74).

## Acknowledgments

The authors would like to acknowledge Kalina Reese for her contributions in designing the figures. Additionally, we would like to acknowledge Guillaume Tremintin (Bruker Daltonics) for his assistance in coupling the Newomics source with Bruker mass spectrometer. The authors would also like to thank Xuefei Sun and Shanhua Lin (Thermo Fisher Scientific) for providing the LC columns used in this study. Y.G. would like to acknowledge National Institutes of Health (NIH) R01 GM125085, R01 HL096971, GM117058 and S10 OD018475. W.G. would like to acknowledge the support from the NIH R01 HL148733, AHA 19TPA34830072, and USDA-NIFA AAI4641. J.A.M. acknowledges support from the Training Program in Translational Cardiovascular Science, T32 HL007936-20 and T32 HL007936-21, for funding during the duration of this project. D.S.R. acknowledges the support from the American Heart Association Predoctoral Fellowship Grant No. 832615/David S. Roberts/2021. K.A.B. acknowledges the Vascular Surgery Research Training Program Grant T32 HL110853. E.A.C. acknowledge support from the NIH Chemistry-Biology Interface Training Program T32 GM008505. Some aspects of the figures were created using BioRender.com.

## Notes

### Competing Interest Statement

Dr. Daojing Wang is the founder & CEO of Newomics.

### Summary of Updates

Removed content related to journal-specific formatting.

