## Supplemental Information for "High Sensitivity Top-down Proteomics Captures Single Muscle Cell Heterogeneity in Large Proteoforms"

### SUPPLEMENTAL METHODS

***Detailed shortening velocity measurements of VL, PLN, and SOL fibers***.

Bundles of ~50 fibers were dissected from the vastus lateralis (VL), plantaris (PLN), and soleus (SOL) muscles of Fischer 344 rats (aged 6 months), tied to glass capillary tubes, and stored at −22 °C for up to 4 weeks in relaxation solution containing 50% (v/v) glycerol. An individual fiber, 1.5-2.5 mm in length, was pulled from the end of the bundle for each experiment and attached to the apparatus. Fibers were attached by placing the ends of the preparation into stainless steel troughs, secured by overlaying a 0.5 mm length of 4–0 monofilament nylon suture on each end, and then tying the suture into the troughs with two loops of 10–0 monofilament suture. This preparation yields very low end compliance and highly uniform striation patterns during $Ca^{2+}$ activations. The length of the preparation was adjusted so that sarcomere length was set to 2.5 μm in relaxation solution and sarcomere length was monitored in pCa 4.5 to ensure sarcomere length did not change significantly during activation. Length changes during contractile measurements were introduced at one end of the preparation driven by voltage commands from a PC via a 16-bit D/A converter. Force and length signals were digitized at 1 kHz using a 16-bit A/D converter and stored on a PC using a custom software in LabView for Windows (National Instruments Corp.). The experimental chamber contained three troughs into which the single fiber was moved to effect rapid solution changes. The apparatus was cooled to 15°C using Peltier devices (Cambion) and a circulating water bath. The entire mechanical apparatus was mounted on a pneumatic vibration isolation table having a cut-off frequency of ~1 Hz.

***Cleaning, tuning, and calibration of the mass spectrometer***

To achieve the highest sensitivity as possible, we thoroughly cleaned the front-end of our mass spectrometer. Contaminants, even minor ones, could have the potential to impair high sensitivity top-down proteomics experiments. We sonicated the spray shield and cap first in water, then equal parts of isopropanol: acetonitrile, and lastly methanol. The capillary was also flushed with the same sequence of solvents. The Bruker maXis II quadrupole time-of-flight mass spectrometer was given time to reach operating pressure and the instrument was tuned and calibrated using ESI tune mix. Instrument parameters were tuned to maximize the intensity and resolution of the tune mix peaks. We found the tuning the extractor fill and extractor tune settings helped increase the intensity to 4e6 and the resolution to 96,000 at 1221 *m/*z. The calibration had a mass error of 0.04 ppm. Based upon the tune and calibration, we were confident that the mass spectrometer was prepared to perform high sensitivity top-down proteomics. Lastly, all mobile phases were made the day of the LC-MS/MS experiments and the LC system was primed and washed with fresh mobile phases.

***Parameters and alignment for the Newomics MnESI source***

In order for the Newomics source to operate at full capacity, source parameters and alignment needed to be optimized. We found that using an end plate offset and capillary voltage set at 500 V and 3750 V, respectively. Nebulizer pressure was set to 0.5 Bar and dry gas flow rate was 4.0 L/min. The dry gas temperature was set to 4.0 L/min and the dry gas temperature was set to 200 ºC. The distance between the emitter and the capillary was measured and optimized to be 3 mm, which is where we found the spray to be stable for all 8 emitters producing an ion current between 1000-2000 nA. **Figure S6** demonstrates the setup of the MnESI platform. The M3 emitter was connected to the LC outlet through a 50 µm ID nanoViper tubing. The emitter position was adjusted using the translation stage for both the x-axis and y-axis with the translation stage. To visualize the M3 emitter and electrospray, a built-in light source and USB camera were installed on the MnESI source.

***Online LC-MS/MS Detailed Parameters***

The capillary column was maintained at a pressure from 1200-2000 psi at room temperature. A total of three replicates (n=3 SMFs) were collected for each fiber type to establish instrument sensitivity and reproducibility (**Figure S8**). The fiber samples were randomized to correct for batch effects. During method development either a Bruker Apollo II ESI source or Newomics MnESI source was used for comparison. The Newomics MnESI source was used for all fiber LC-MS/MS analyses and source parameters can be found in previous protocol. The ESI source nebulizer pressure was set to 0.5 Bar, the dry gas pressure was 4.0 L/min, and the dry gas temperature was set to 220 $^{o}$C. The end plate offset and capillary voltage set at 500 V and 4500 V, respectively. 30 V of in-source collisional energy was applied to facilitate ionization. Intact mass spectra were collected at a scan rate of 1.0 Hz over the mass range of 200-3000 *m/z*.

***Data Analysis***

Bruker Data Analysis version 4.3 was used to analyze the LC-MS data from the fiber samples. All chromatograms shown were smoothed using the Gauss algorithm with a smoothing width of 1.05 sec. The mass spectra were averaged over a window where all proteoforms of the same protein eluted. All protein elution windows were consistent between samples due to the high reproducibility of our method.

Targeted top-down proteomics was performed on single fiber contractile proteins of interest. Tandem mass spectra (MS/MS) were analyzed using the MASH Explorer software (Version 2.0), which was developed in-house (1). Peak extraction of the spectra were deconvoluted using eTHRASH with a signal-to-noise ratio of 3 and a cutoff fit score of 60%. All the program-processed data were manually validated to obtain accurate sequence and PTM information. All identifications were made within a 20 ppm mass tolerance when matching the experimental fragment ions to the calculated fragment ions based on the amino acid sequence.

Discovery top-down proteomics was also performed on a representative VL, PLN, and SOL fiber using a database searching algorithm. A representative VL, PLN, and SOL RAW file were converted into mzXML files using the msconvert algorithm (2). TopFD (version 1.5.4) was used for spectral deconvolution and TopPIC (3) was used for database searching (Version 1.5.4) embedded within Mash Explorer software (1). Relevant TopPIC parameters are listed here: The search included no fixed modifications and up to 2 unexpected PTMs, with a maximum mass shift of 500 Da. The precursor mass tolerance was 0.02 *m/z* and fragment mass error tolerances were 15 ppm with a 1.2 Da mass window for proteoform spectral matches (PRSMs), a maximum charge of 50, and a maximum proteoform mass of 100 kDa. The E-value cutoff was set to 0.01. Canonical entries of the *Rattus noveregicus* UniProt database (UP000002494, 8,165 reviewed entries, version October 20, 2022) were used for the database search. To analyze TopPIC output proteoform spectral matches, the R package TopPICR was used to filter out redundant identifications to report only one identification per spectral feature and filer low-confidence identifications to establish a false discovery rate of 1%. Filtered

TopPIC search results were then used for gene ontology analysis by STRING version 11.5 (4) with highest confidence selected with the whole rat genome (22,763 proteins) as the enrichment background.

### SUPPLEMENTAL NOTES

#### Supplemental Note 1: *Evaluation of various protein extraction methods*

We evaluated various protein extraction methods using a minimal amount of tissue samples that can be weighed accurately (1 mg) (**Figure S1-S2**). A small piece of tissue (1 mg) from PLN muscle was first homogenized in 40 µL HEPES buffer (25 mM HEPES pH 7.5, 60 mM NaF, 1 mM $Na_3VO_4$, 1 mM PMSF, 1 mM Tris-(2-carboxyethyl) phosphine (TCEP), and 1 mM L-methionine) using a Teflon pestle (1.5 mL tube rounded tip; Scienceware, Pequannock, NJ, USA) (5–8). The homogenate was centrifuged (21,000 rcf, 30 min, 4 ºC) and the supernatant was discarded. The resultant pellet was extremely small in size to mimic the small amount of protein content in a single fiber. We homogenized the pellets in several different extraction solutions: 1) a photocleavable surfactant (0.1% Azo6 in 25 mM ammonium acetate (9)), 2) a non-ionic surfactant (1% DDM (10)) 3) trifluoroacetic acid (1% TFA (5)), and hexafluorisopropanol (100% HFIP (11)). Azo (12) and DDM (10) are MS-friendly surfactants at low concentrations and TFA and HFIP are known to extract contractile proteins while maintaining MS-compatibility (5, 7, 11, 13, 14). The pellets were homogenized in 20 µL of each of the lysis solutions. Protease and phosphatase inhibitors, as well as antioxidants, were included in all extraction solutions to minimize artifactual protein modification prior to LC-MS/MS analysis. The homogenate, which contained primarily sarcomeric proteins, was centrifuged (21,000 rcf, 30 min, 4 ºC) and the supernatant was collected for SDS-PAGE analysis.

An additional Azo and DDM extraction following the same protocol was performed to test the amount of protein loss during subsequent MS-compatibility steps. For the photocleavable sample, the surfactant was first degraded with UV light for 3 min in a degradation solution (50% isopropanol, 30 mM L-methionine, 2 mM TCEP, 1% formic acid (FA)) to keep proteins soluble. The degradation products of the reaction formed a salt which can be quickly removed prior to MS-analysis (Amicon 10k molecular weight cutoff filter). The DDM sample was also desalted to test the amount of sample loss. The samples were desalted four cycles with 0.1% FA in water and concentrated to a final volume of 20 µL. 10 µL of each extraction condition's supernatant (n=6 total) were loaded onto a homecast 12.5% acrylamide gel and separated by electrophoresis. The gel was stained following the SYPRO™ Ruby protocol and imaged on a gel imager. As expected, the use of surfactants aided protein extraction, but the sample loss resulting from the downstream clean-up procedures prior to MS analysis outweighed the benefits for high sensitivity single cell analysis (**Figure S2**).

**Supplemental Note 2:** ***Determining the most effective lysis method for single fiber samples***

Next, we sought to evaluate the lysis methods for high sensitivity single fiber (multinucleated single muscle cell) analysis. Although the use of minimal amount of tissue (1 mg) has been valuable in the initial method developments studies, the protein content is still much greater than that of a single fiber. In addition, the cellular components found in skeletal muscle tissue differ from that of a skinned single fiber. Given the small size of the single fiber, mechanical homogenization is not practical. Thus, we explored other possibilities, including sonication in a water bath and freeze-thaw lysis, to aid sample breakdown and protein extraction while simultaneously minimizing adsorptive protein loss in this one-pot extraction method. The previous bottom-up single cell proteomics studies have employed cell lysis in acidified water using consecutive cycles of freezing at -80 °C then thawing immediately at higher temperatures (>37 ºC) (15–18). Additionally, sonication devices have been explored for cell lysis as well.

We tested three separate conditions to determine the most effective lysis method for single fiber samples: a freeze-thaw lysis with the single fiber in 100% HFIP, a freeze-thaw lysis (15–18) with the single fiber in 0.1% FA in water (MPA), and water-bath sonication with the fiber in 100% HFIP (**Figure S3**). First, fibers of similar size were obtained from PLN tissue and dissected into low-bind micro centrifuge tubes. Samples were washed with 40 µL of 150 mM ammonium acetate to remove any remaining relaxation buffer by pipetting the solution over the fiber such that the fiber remained on the wall of the tube. Samples were briefly centrifuged (1,100 × g, 1 min, 4 ºC) and the solution was removed. 20 µL of each extraction solvent was added into the tubes and the fibers were suspended in the liquid. The samples were incubated for 15 min on ice. Two samples underwent freeze-thaw lysis (incubation at -80 ºC for 5 min followed by incubation for 1 min at 37 ºC). The freeze-thaw cycle was repeated two additional times for a total of three cycles with mixing by gentle agitation between cycles. The fiber extracts were desalted prior to LC-MS/MS analysis using a 10 kDa MWCF and buffer exchanged using 0.1% FA in nanopure water concentrating to a final volume of 20 µL. 5 µL of each desalted extract was analyzed via LC-MS/MS method following a similar protocol described in the main text. Our results clearly show that freeze-thaw lysis greatly aids in extraction of proteins from single fiber samples (**Figure S3)**.

**Supplemental Note 3:** ***Determining optimal HFIP concentration for single fiber extraction***

We next sought to determine the optimal concentration of HFIP in the extraction solution. An important benefit from the use of a lower percentage of HFIP is its compatibility with reversed phase LC (RPLC). Samples were washed with 40 µL of 150 mM ammonium acetate to remove any remaining relaxation buffer by pipetting the solution over the fiber such that the fiber remained on the wall of the tube. Samples were briefly centrifuged (1,100 × g, 1 min, 4 ºC) and the solution was removed. The fibers were suspended in 20 µL of either 100%, 50%, 25%, or 12.5% HFIP. The samples were incubated for 15 min on ice. All samples underwent freeze-thaw lysis (incubation at -80 ºC for 5 min followed by incubation for 1 min at 37 ºC). The freeze-thaw cycle was repeated two additional times for a total of three cycles with mixing by gentle agitation between cycles. The fiber extracts were desalted prior to LC-MS/MS analysis using a 10 kDa MWCF and buffer exchanged using 0.1% FA in nanopure water concentrating to a final volume of 20 µL. 5 µL of each desalted extract was analyzed via an LC-MS/MS method following a similar protocol described in the main text. We found that 25% HFIP was the optimal concentration of HFIP while maintaining MS-compatibility (**Figure S4)**.

**Supplemental Note 4:** ***Testing whether the freeze-thaw lysis affects fiber proteoforms***

A potential concern of using freeze-thaw lysis for protein extraction was that this procedure could lead to protein degradation or the loss of labile PTMs, such as phosphorylation (19). To do so, we chose to compare the results of an HFIP extraction followed by either a freeze-thaw lysis or no freeze-thaw lysis with that of previously published literature (5, 7, 8, 19–21). This method uses TFA, which we found to be ineffective at extracting proteins from a single fiber, but works effectively on tissue. Briefly, ~10 mg of swine right ventricle tissue (n=4) was homogenized in 100 µL HEPES buffer (25 mM HEPES pH 7.5, 60 mM NaF, 1 mM $Na_3VO_4$, 1 mM PMSF, 1 mM Tris-(2-carboxyethyl) phosphine (TCEP), 1 mM L-methionine, 1x protease inhibitor (Halt™ Protease inhibitor cocktail, Thermo) and 1x phosphatase inhibitor cocktail A (catalog# sc-45044, Santa Cruz Biotechnology, Inc)). The samples were centrifuged (21,000 × g, 30 min, 4 $^{o}$C) and the supernatant was removed. Two of the pellets were homogenized in a 25% HFIP solution and the remaining two pellets were homogenized in a 1% TFA solution. The samples were centrifuged (21,000 × g, 30 min, 4 $^{o}$C) and the supernatant was transferred to a new tube. One of the HFIP samples and one of the TFA samples were selected for freeze-thaw lysis is described above. 5 µL of each desalted extract was analyzed via LC-MS/MS method following a similar protocol described in the main text. Our results have shown no discernible changes in PTMs or proteolysis resulting from freeze-thaw lysis (**Figure S5**).

### SUPPLEMENTAL TABLES

Table S1. Sarcomeric proteoforms identified| Gene, Uniprot ID, retention time (RT), mono-isotopic mass (Mr Calc'd), proteoform name, and processing and PTMs for the proteoforms identified and quantified in this study. Proteoforms were either identified by MS/MS in this study or by accurate mass measurements based on previous publications or information derived from Uniprot. Tropomyosin (Tpm) isoform nomenclature was adopted from Geeves et al.(14), but common names are also included in parentheses. Abbreviations: fast skeletal Troponin T 3 (fsTnT3), slow skeletal Troponin T 1 (ssTnT1), slow skeletal Troponin I 1 (ssTnI), fast skeletal Troponin I 3 (fsTnI3), myosin light chain 1, skeletal muscle isoform (MLC-1F), myosin light chain 6B (MLC-1S), myosin regulatory light chain 2, skeletal muscle isoform (MLC-2F), Myosin light chain 3, skeletal muscle isoform (MLC-3F), Myosin regulatory light chain 2, cardiac muscle isoform (MLC-2S), fast skeletal Troponin C (fsTnC), SLOW skeletal Troponin C (ssTnC), acetylation (acetyl), phosphorylation (phospho), methionine removal (-Met), methylation (methyl).

| Gene | UniProt ID | RT (min ) | Monoisotopic Calc (Da) | Monoisotopic Exp't (Da) | Proteoform | Processing and PTMs |
|---|---|---|---|---|---|---|
| Tnnt3 | P09739 | 24.2 | 29,219.33 | 29,219.37 | fsTnT3 | Acetyl, Phospho |
| Tnnt1 | Q7TNB2 | 26.1 | 31,107.07 | 31,107.17 | ssTnT1 | -Met, Acetyl, Phospho |
| Tnni1 | Q9WUZ5 | 26.2 | 21,553.49 | 21,553.58 | ssTnI | -Met |
| Tnni2 | P27768 | 26.7 | 21,225.83 | 21,225.94 | fsTnI | -Met, Acetyl |
| Ldb3 | Q5XIG1 | 28.3 | 30,879.76 | 30,879.82 | Cypher4s | -Met, Acetyl |
| Ldb3 | Q9JKS4 | 28.6 | 31,319.00 | 31,319.08 | Cypher2s | -Met, Acetyl |
| Myl3 | P16409 | 32.1 | 22,053.06 | 22,053.19 | MLC-1V | -Met, Acetyl |
| Tpm2 | P58775 | 32.3 | 32,858.56 | 62,858.68 | β-Tpm | Acetyl |

| | | | | | | |
|---|---|---|---|---|---|---|
| Tpm1 | P04692 | 32.5 | 32,702.68 | 32,702.76 | α-Tpm | Acetyl |
| Myl1 | P02600 | 33.5 | 20,577.43 | 20,577.56 | MLC-1F | -Met, Acetyl |
| LOC120093525 | D3ZHA7 | 33.8 | 22,689.70 | 22,689.80 | MLC-1S | -Met, Dimethyl |
| Myl11 | P04466 | 34.2 | 18,868.32 | 18,868.41 | MLC-2F | -Met, Acetyl |
| Myl1 | P02600-2 | 34.8 | 16,514.11 | 16,514.15 | MLC-3F | -Met, Acetyl |
| Myl2 | P08733 | 34.9 | 18,779.36 | 18,779.46 | MLC-2S | -Met, Acetyl |
| Acta1 | P68136 | 37.5 | 41,845.83 | 41,846.00 | Actin | -Met, Acetyl, methyl H92 |
| Tnnc2 | Q304F3 | 38.2 | 17,995.30 | 17,995.39 | fsTnC | -Met, Acetyl |
| Tnnc1 | Q4PP99 | 38.9 | 18,450.50 | 18,450.58 | ssTnC | -Met, Acetyl |

### SUPPLEMENTAL FIGURES

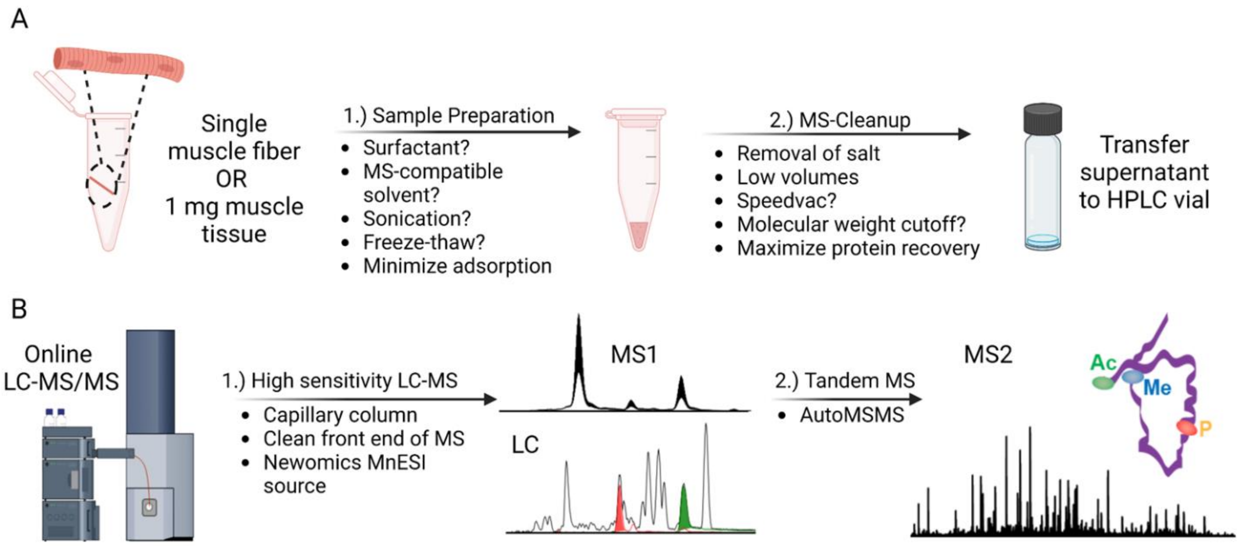


**Figure S1. Method development strategy for high sensitivity top-down proteomics. A)** The first step in the development of a high sensitivity top-down proteomics method was to minimize sample losses that occur during protein extraction and clean-up to ensure MS-compatibility. Initial method development was done with 1 mg of skeletal muscle tissue but when the method became more sensitive the remaining method development steps were performed on single fibers. We found that extracting proteins from SMFs with 25% HFIP and a freeze-thaw lysis balanced extraction efficiency with MS-compatibility. **B)** Once the sample preparation steps were optimized, we developed a high sensitivity LC-MS/MS method that uses a capillary reversed phase LC column to minimize sample dilution during the separation, the Newomics MnESI source for enhanced ionization efficiency, and an AutoMSMS method for proteoform characterization.

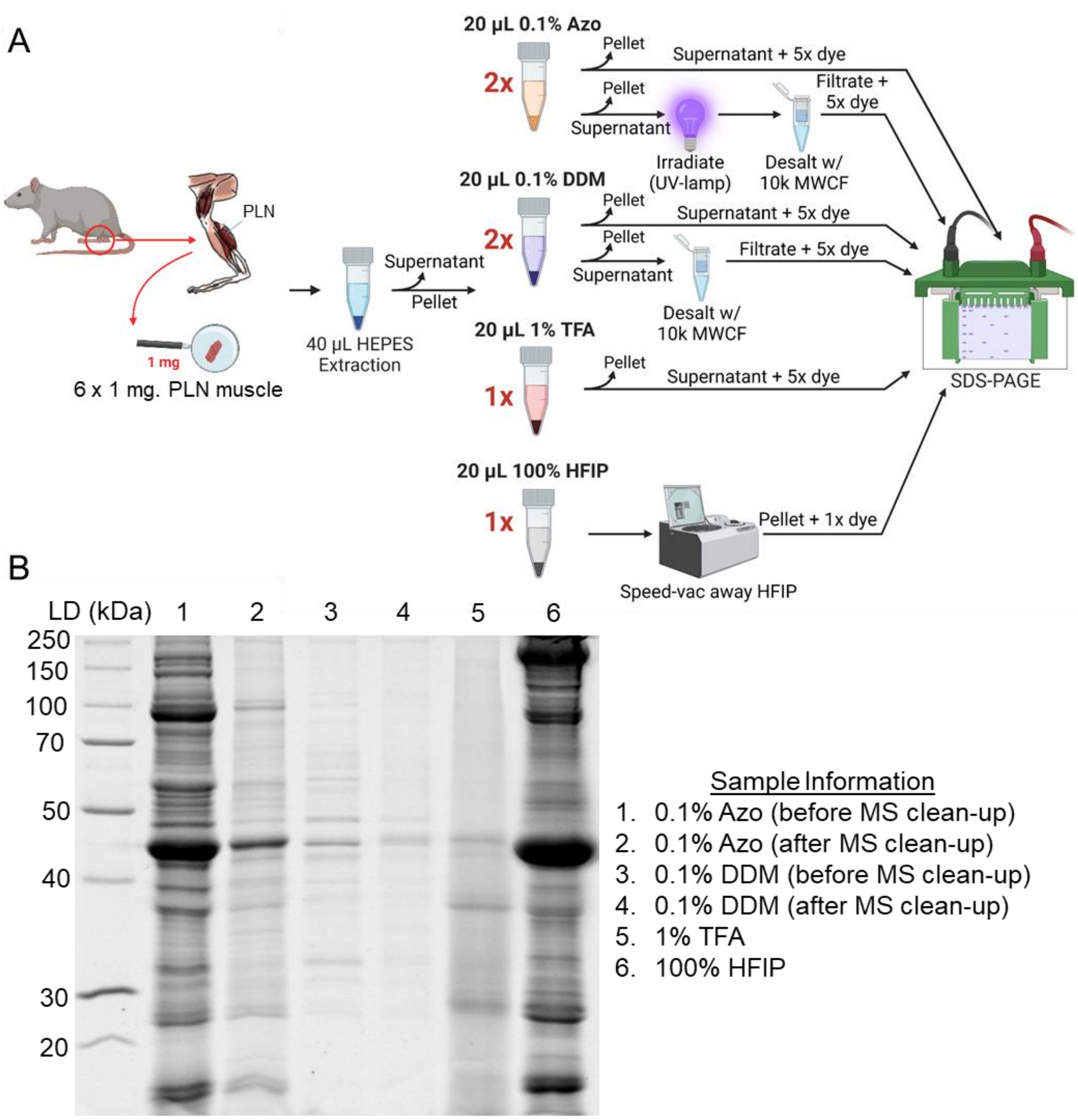


**Figure S2. SDS-PAGE analysis of various extraction methods from 1 mg skeletal muscle tissue shows that HFIP has the highest extraction efficiency while maintaining facile MS clean-up. A)** Workflow for experiment to test various extraction methods from 1 mg PLN muscle. **B)** We tested MS-friendly, photo-cleavable surfactant, Azo, commonly used non-ionic surfactant, DDM, 1% TFA, and 100% HFIP. 0.1% Azo in (Lane 1) and 25% HFIP (Lane 6) have shown the highest extraction efficiency. However, due to small amount of protein content and the relatively larger amount of Azo needed (CMC 0.4%), the degradation of Azo caused more significant protein loss than in the larger amount of sample.

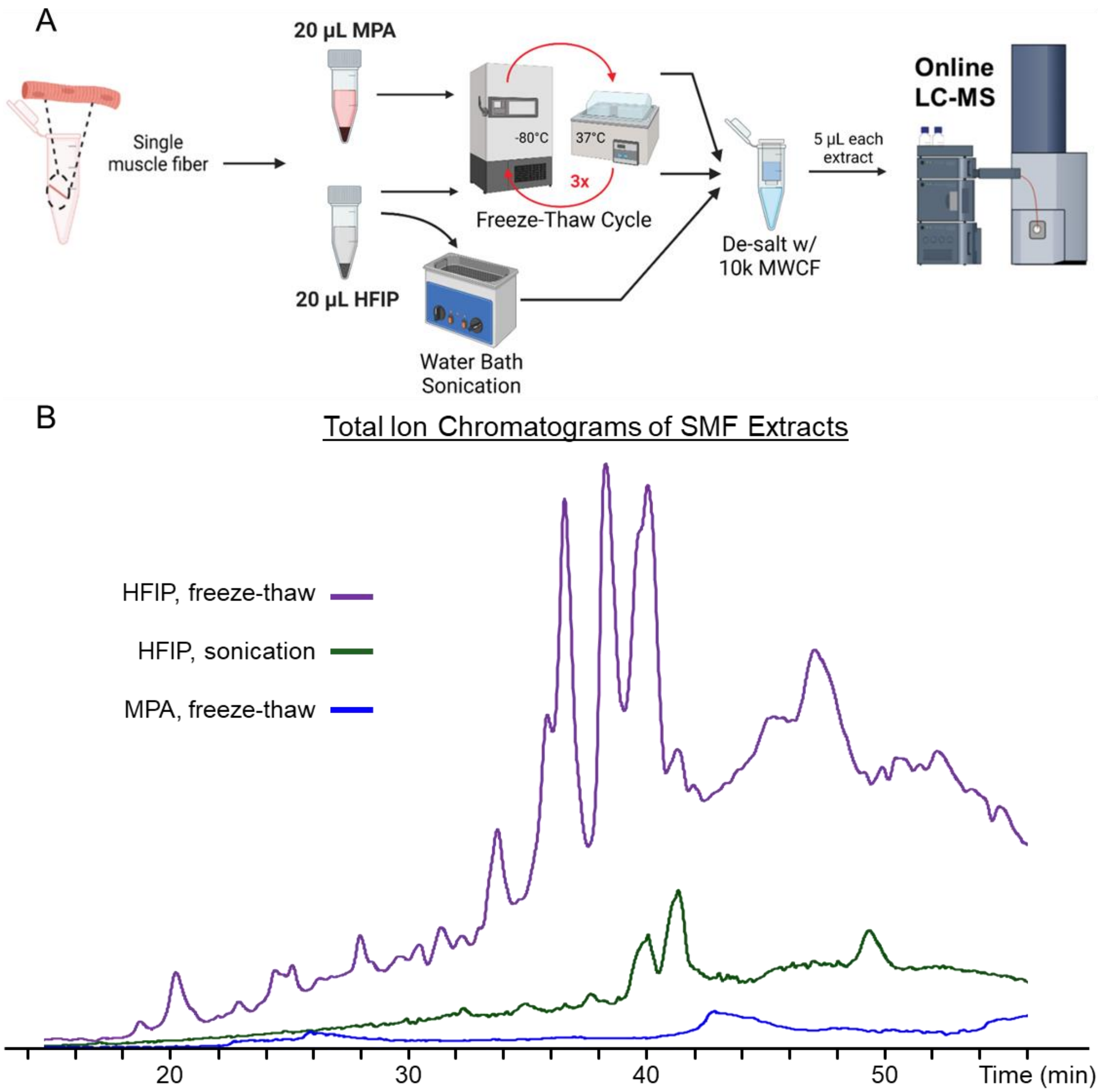


**Figure S3. LC-MS analysis of various lysis methods for SMFs shows that freeze-thaw enhances extraction efficiency. A)** Workflow for experiment to test various lysis methods to increase extraction efficiency from single fibers. **B**) Total ion chromatograms of equal volume injections of lysates obtained from extractions that used either a freeze-thaw lysis in HFIP or MPA as well as a water bath sonicator lysis in HFIP. For single fiber studies, freeze-thaw lysis greatly aids in the extraction of contractile proteoforms.

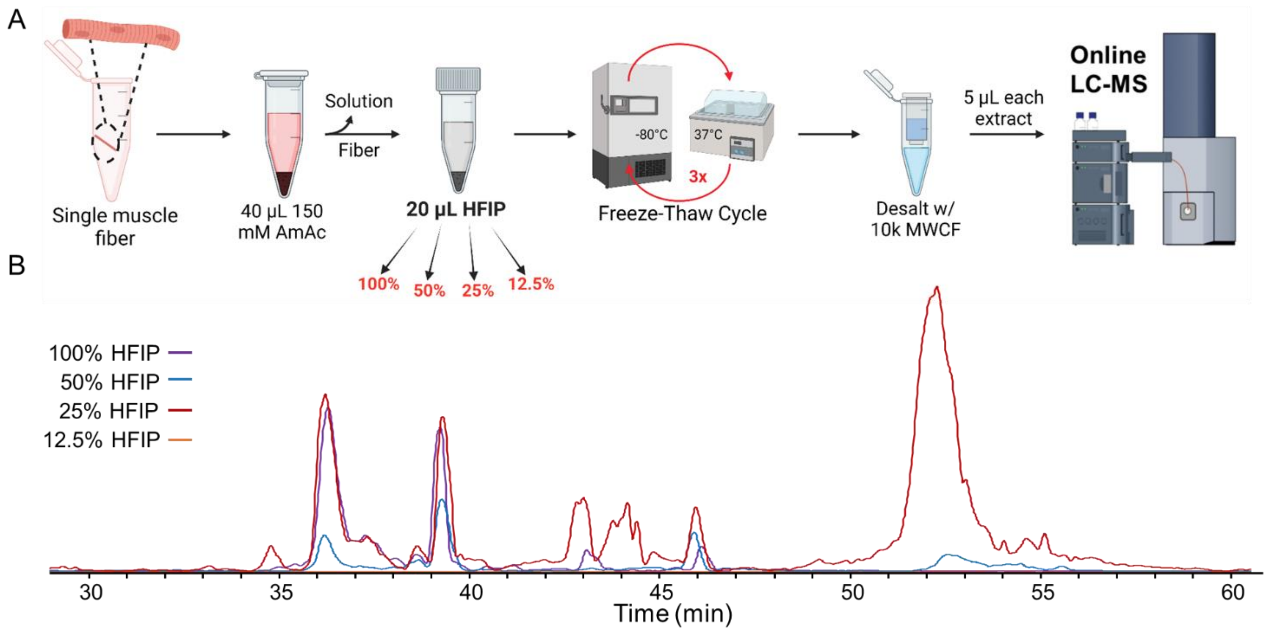


**Figure S4. 25% HFIP performs the best at extracting proteins from SMFs as shown by LC-MS. A)** Workflow for experiment to test optimal the HFIP concentration for single fiber extraction. **B)** Total ion chromatograms of equal volume injections of lysates obtained from extractions that used 100%, 50%, 25%, or 12.5% HFIP, respectively. The 25% HFIP extraction worked the best for the single fiber samples with the benefit of having an organic composition closer to the starting conditions of RPLC.

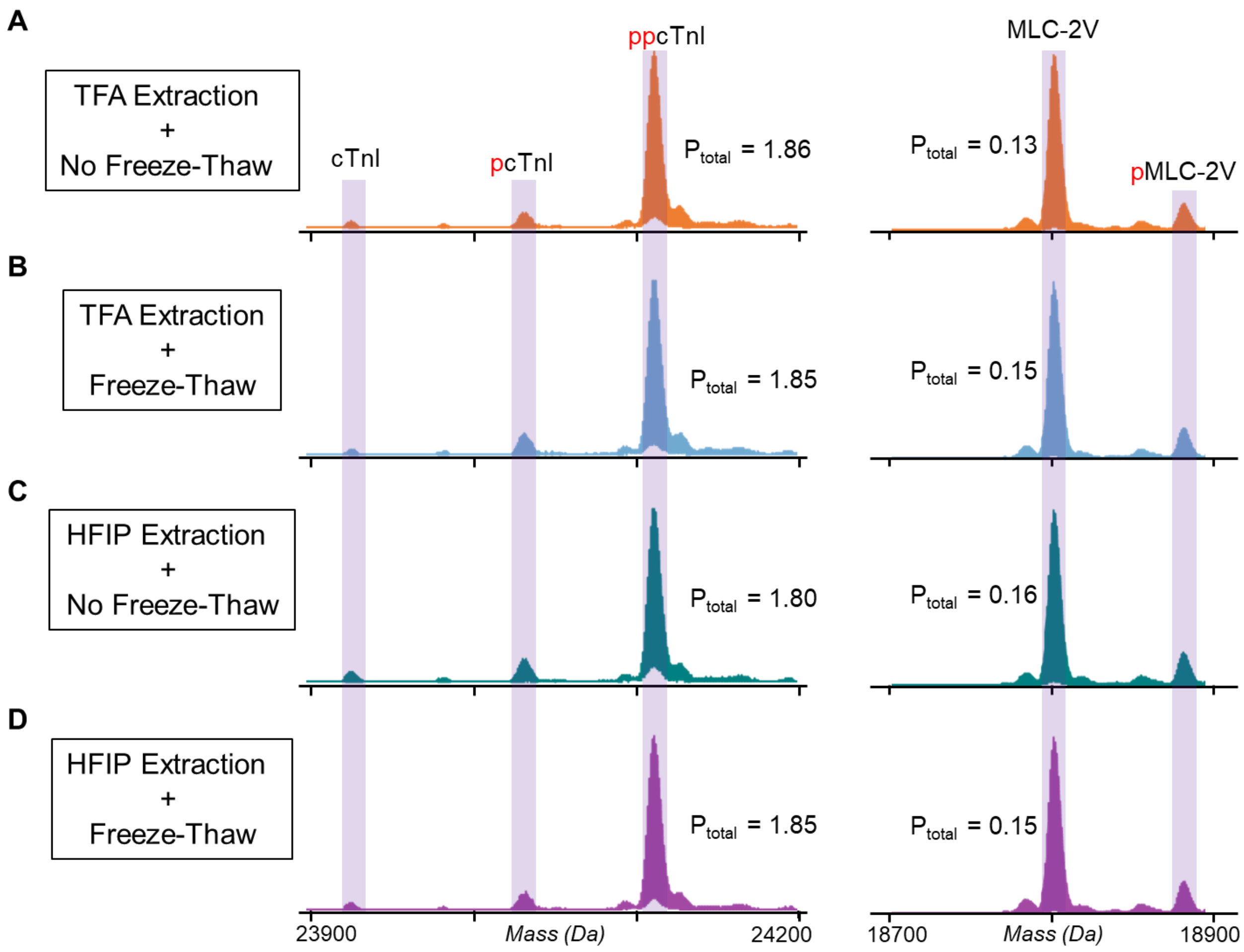


**Figure S5. Freeze-thaw lysis does not alter proteoforms.** Deconvoluted mass spectra of cardiac troponin I (cTnI) and the ventricular isoform of MLC-2 (MLC-2V) from **A)** lysate obtained from 1% TFA extraction without freeze-thaw lysis **B)** lysate obtained from 1% TFA extraction with a freeze-thaw lysis, **C)** lysate obtained from 25% HFIP extraction without freeze-thaw lysis and **D)** lysate obtained from 25% HFIP extraction with freeze thaw lysis. The Ptotal values are highly similar for all the different extraction methods. Mono-phosphorylation is indicated by red "p", bis-phosphorylation is indicated by red "pp". Total phosphorylation level, Ptotal (mol Pi/mol protein), was calculated to determine if there were any major changes in proteoform abundance.

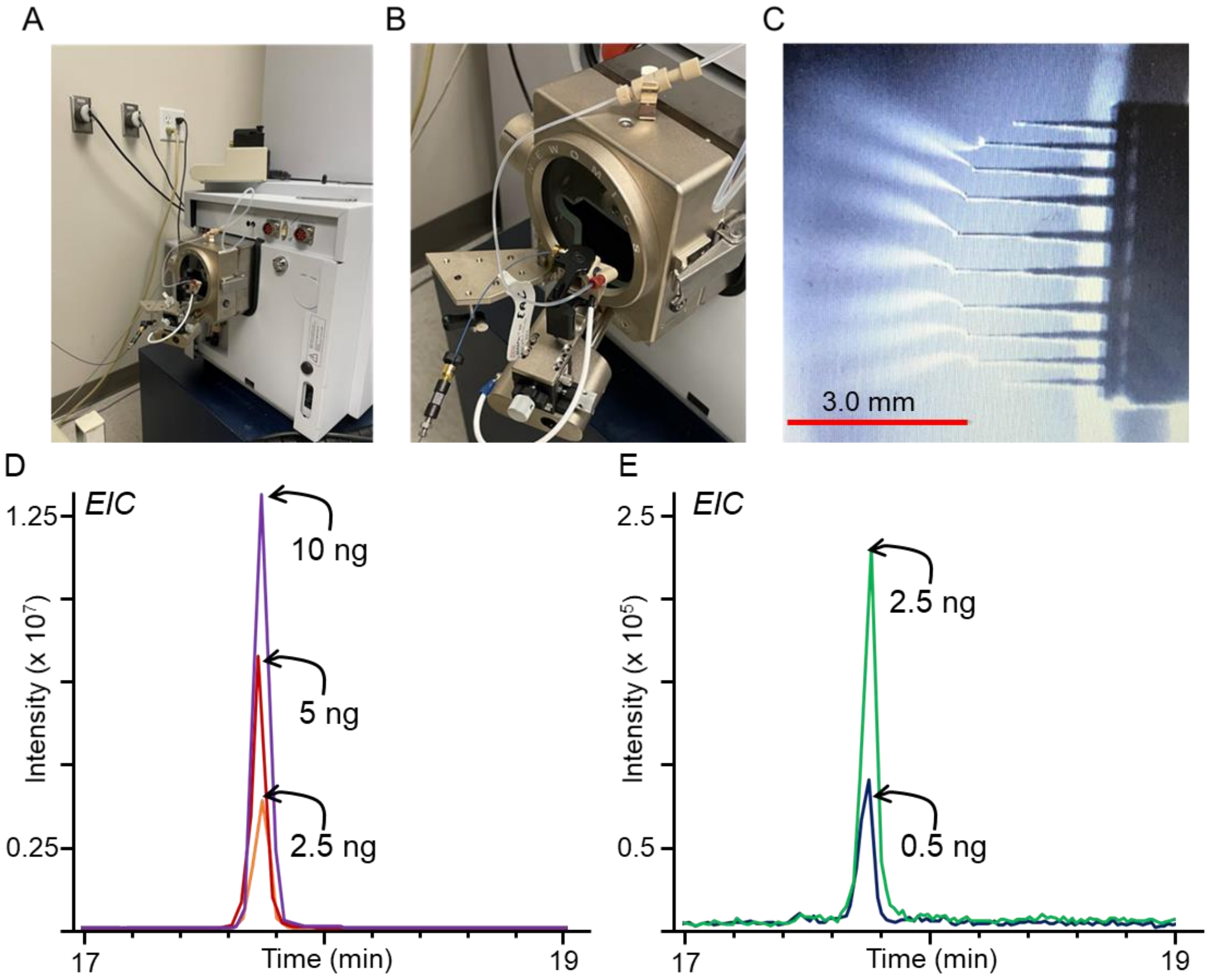


**Figure S6. Setup for the LC-MnESI-q-TOF configuration and evaluation of detection limit. A)** Picture of the Newomics MnESI source connected to a Bruker MaXis II q-TOF mass spectrometer. The MnESI source inlet is attached to a Thermo MabPac capillary column flowing at 2 µL/min. **B)** Zoom-in picture of the MnESI source. The M3 emitter fits inside the black housing containing the column intlet. **C)** Picture taken from a built-in camera and light source to visualize nanoelectrospray. All 8 nanoelectrospray emitters are effectively spraying towards the mass spectrometer inlet ensuring proper function of the source. **D)** Three overlaid EICs (top 5 most abundant ions) of a 10, 5, and 2.5 ng injection of carbonic anhydrase (CA) the LC-MnESI-q-TOF configuration. **E)** Two overlaid EICs (top 5 most abundant ions) of a 1 and 0.5 ng injection of CA the LC-MnESI-q-TOF configuration. The detection limit of our method is approximately 0.5 ng for CA, a standard 29 kDa protein.

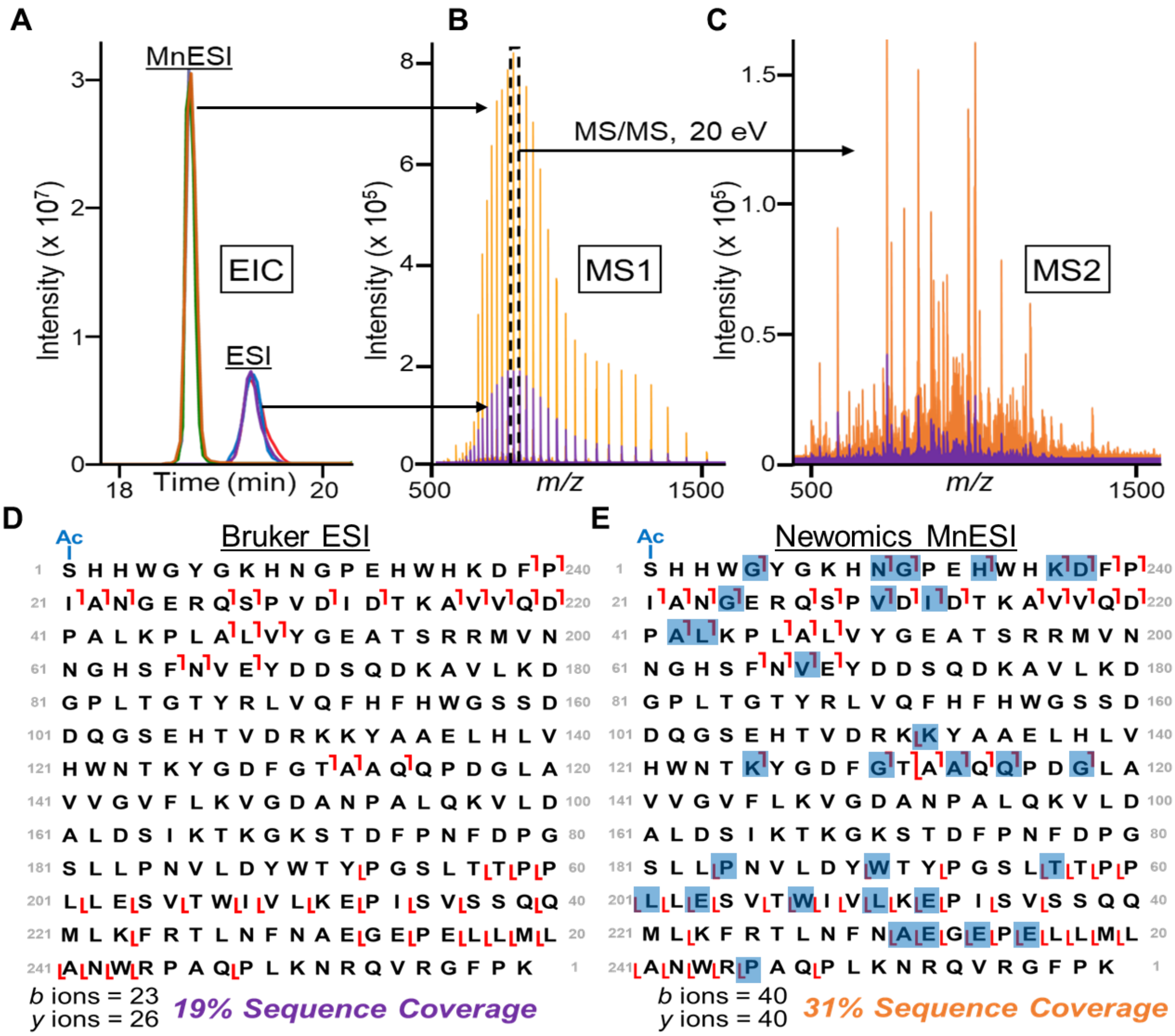


**Figure S7. Newomics MnESI provides a 4x increase in MS1 sensitivity compared to conventional ESI source, which results in higher top-down sequence coverage. A)** Three overlaid EICs of a 10 ng injection of carbonic anhydrase (CA) using both the Newomics MnESI and Bruker ESI sources. The CA peak elutes earlier in the MnESI source due to smaller dead volume in the source, which results in sharper chromatographic peaks as well. **B)** CA MS1 spectra using MnESI source (orange) versus the ESI source (purple). The most abundant precursor was selected for further fragmentation. **C)** Tandem MS data as a result of fragmenting the most abundant precursor with 20 eV of CID energy. **D)** Fragmentation table generated from fragment ions matched within a 10 ppm mass error using the ESI source. In total, this experiment confidently identified 23 *b* ions and 26 *y* ions resulting in 19% sequence coverage. **E)** Fragmentation table generated from fragment ions matched within a 10 ppm mass error using the MnESI. In total, the MnESI source confidently identified 40 *b* ions and 40 *y* ions resulting in 31% sequence coverage.

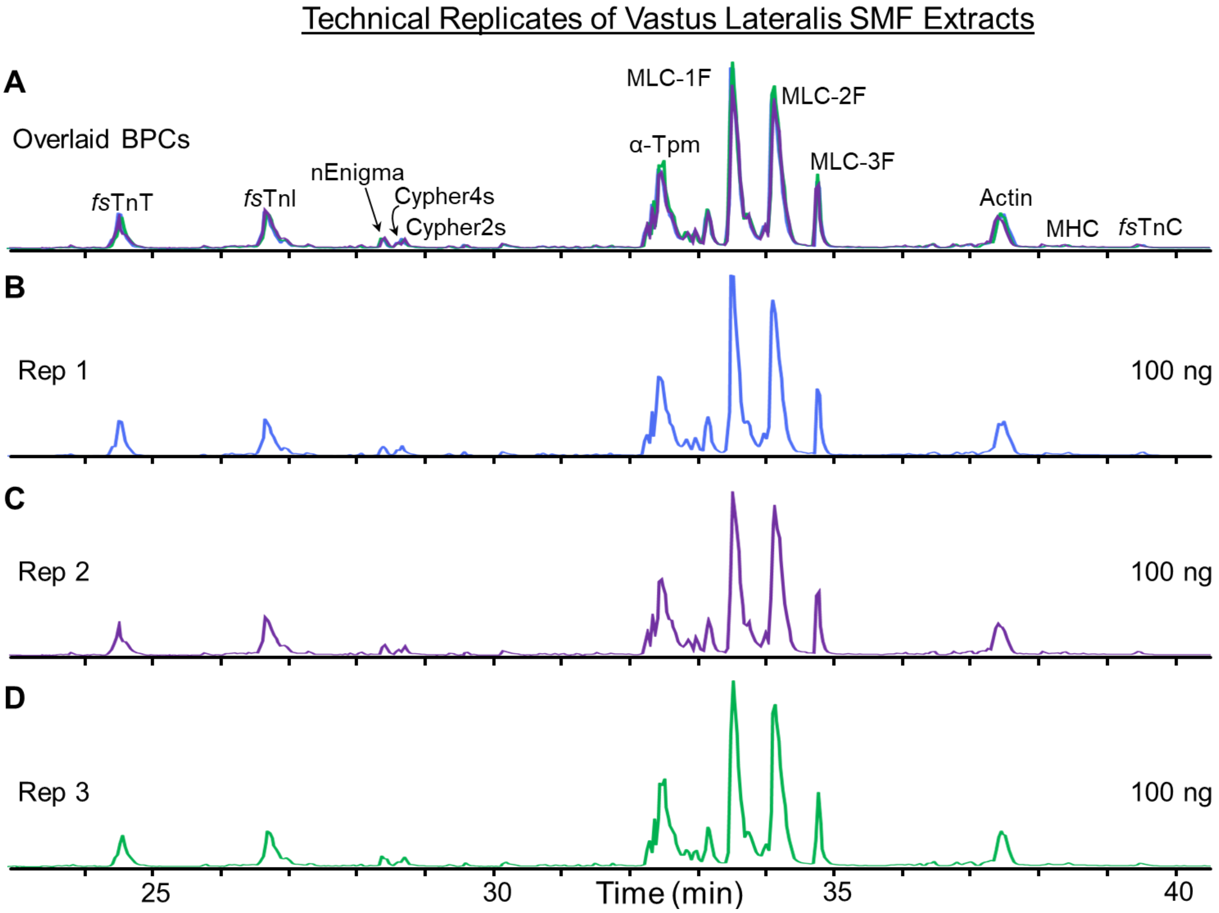


**Figure S8. Highly reproducible high sensitivity top-down LC-MS.** Representative BPCs for consecutive injection replicates (n=3) of 100 ng of total lysate from a VL single fiber sample.

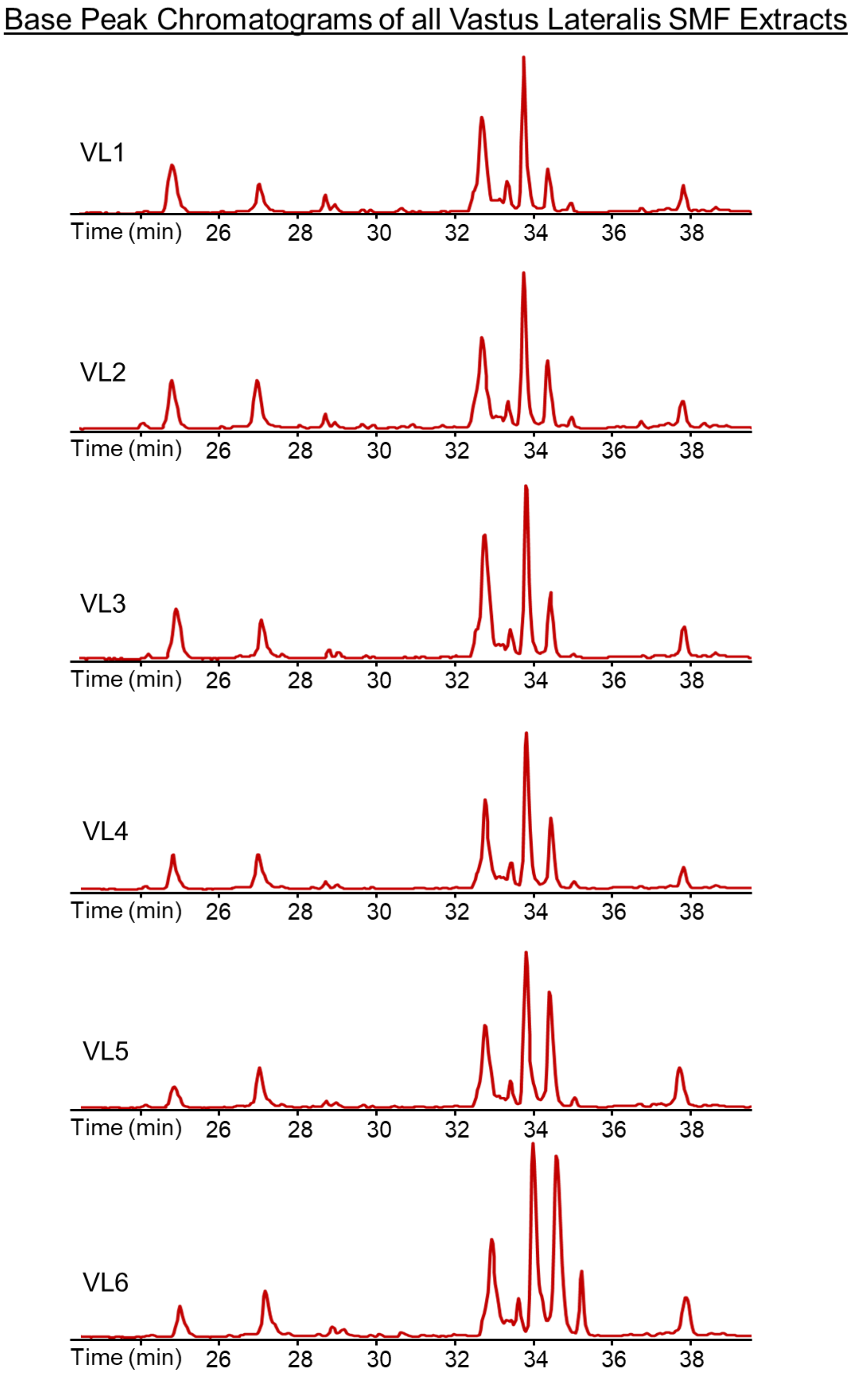


**Figure S9. Stacked BPCs show highly reproducible LC-MS results.** Approximately 100 ng of single fiber lysate from vastus lateralis (VL) was injected for each LC-MS run. Six SMFs were used in this analysis.

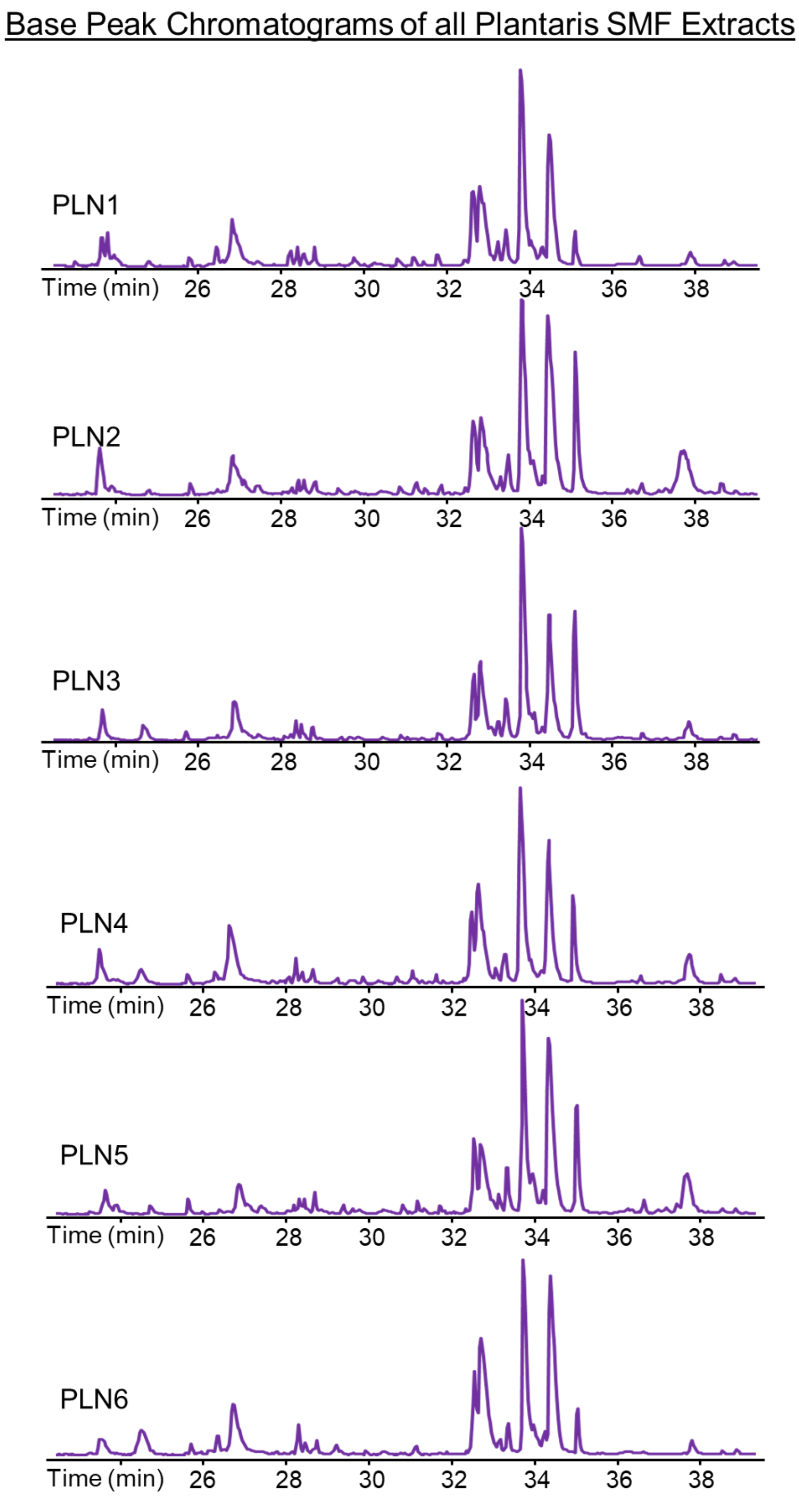


**Figure S10. Stacked BPCs show reproducible LC-MS results.** Approximately 100 ng of single fiber lysate from plantaris (PLN) was injected for each LC-MS run. Six SMFs were used in this analysis.

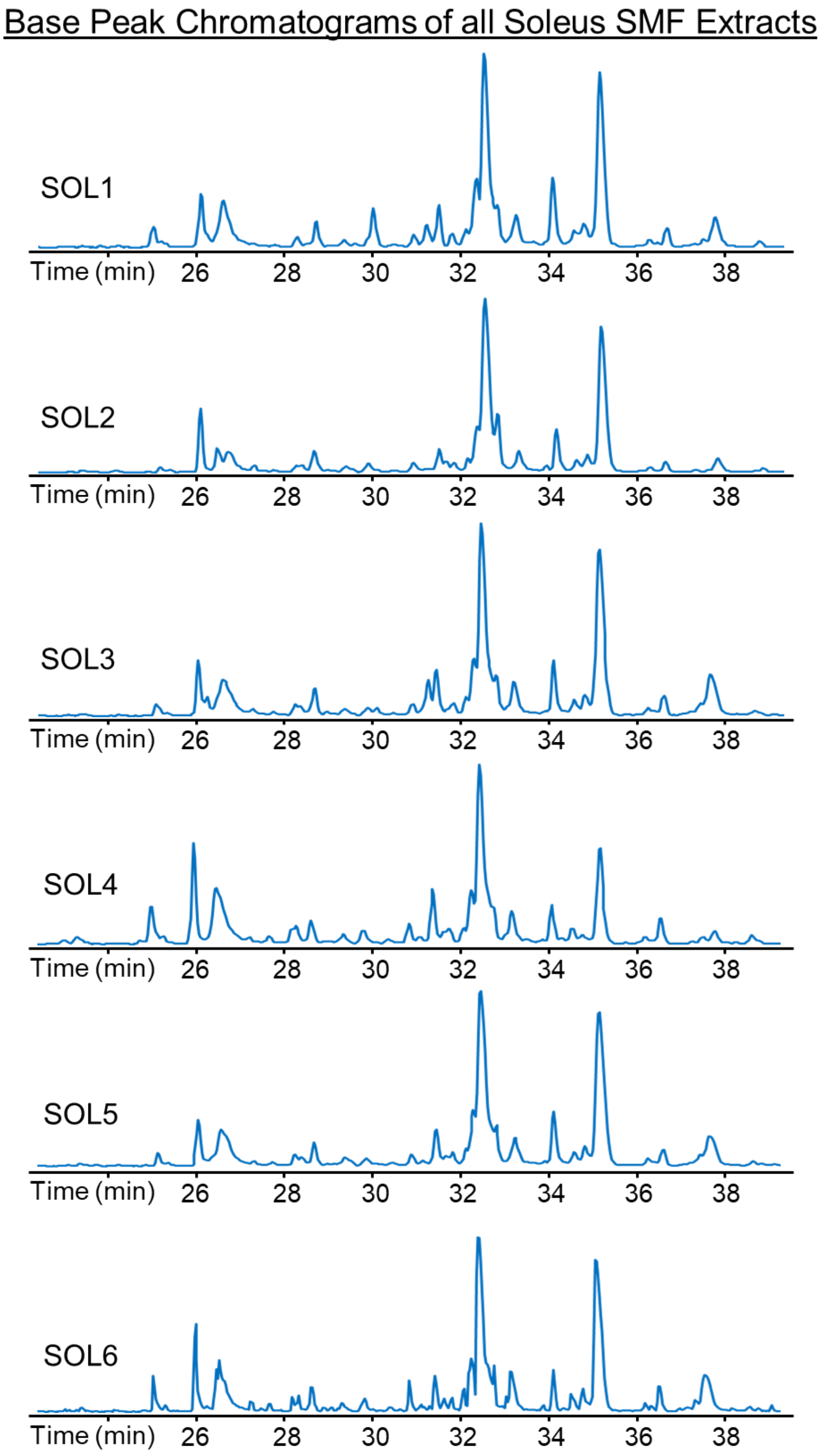


**Figure S11. Stacked PLN BPCs show highly reproducible LC-MS results.** Approximately 100 ng of single fiber lysate from soleus (SOL) was injected for each LC-MS run. Six SMFs were used in this analysis.

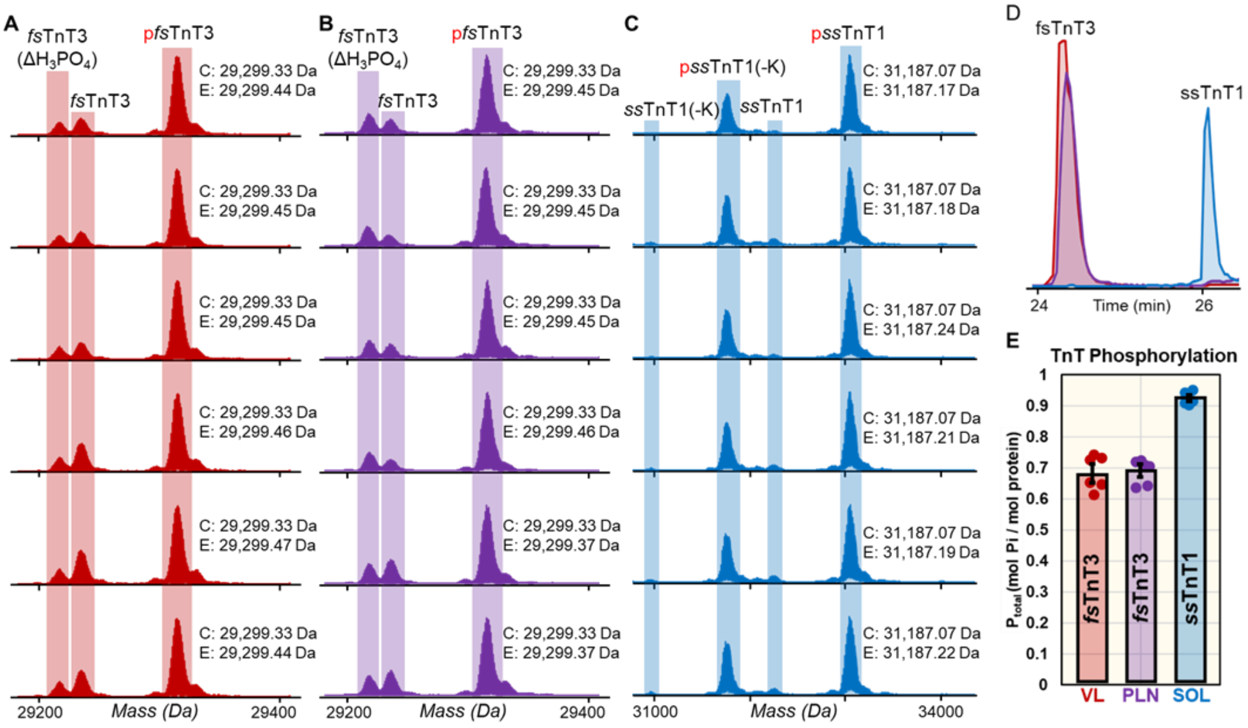


**Figure S12. Top-down proteomics provides a "bird's eye view" of TnT isoforms and proteoforms from single muscle cells.** Deconvoluted mass spectra for TnT isoforms closely match predicted masses for SMFs obtained from **A)** VL (fsTnT3), **B)** PLN (fsTnT3), and **C)** SOL (ssTnT1) muscles. Monoisotopic masses reported for the calculated (C) and experimental (E) masses of TnT isoforms. The UniProt accession number, P09739, was used for the fsTnT3 calculated mass. The UniProt accession number, Q7TNB2, was used for the ssTnT1 calculated mass. The calculated and experimental masses are within a 10 ppm mass error. Mono-phosphorylation is indicated by red "p", "$\Delta H_3PO_4$" indicates a loss of phosphate from pfsTnT, and "-K" indicates loss of a lysine residue from ssTnT1 or pssTnT1. **D)** fsTnT3 and ssTnT1 EICs (top 5 most abundant ions) from VL, PLN, and SOL single fibers. The LC separation shows high reproducibility of fsTnT3 separation for VL and PLN fibers and are separated from the later eluting ssTnT1. **E)** Total phosphorylation level, Ptot (mol Pi/mol protein), for fsTnT3 and ssTnT1 from VL, PLN, and SOL fibers (n=6 per tissue).

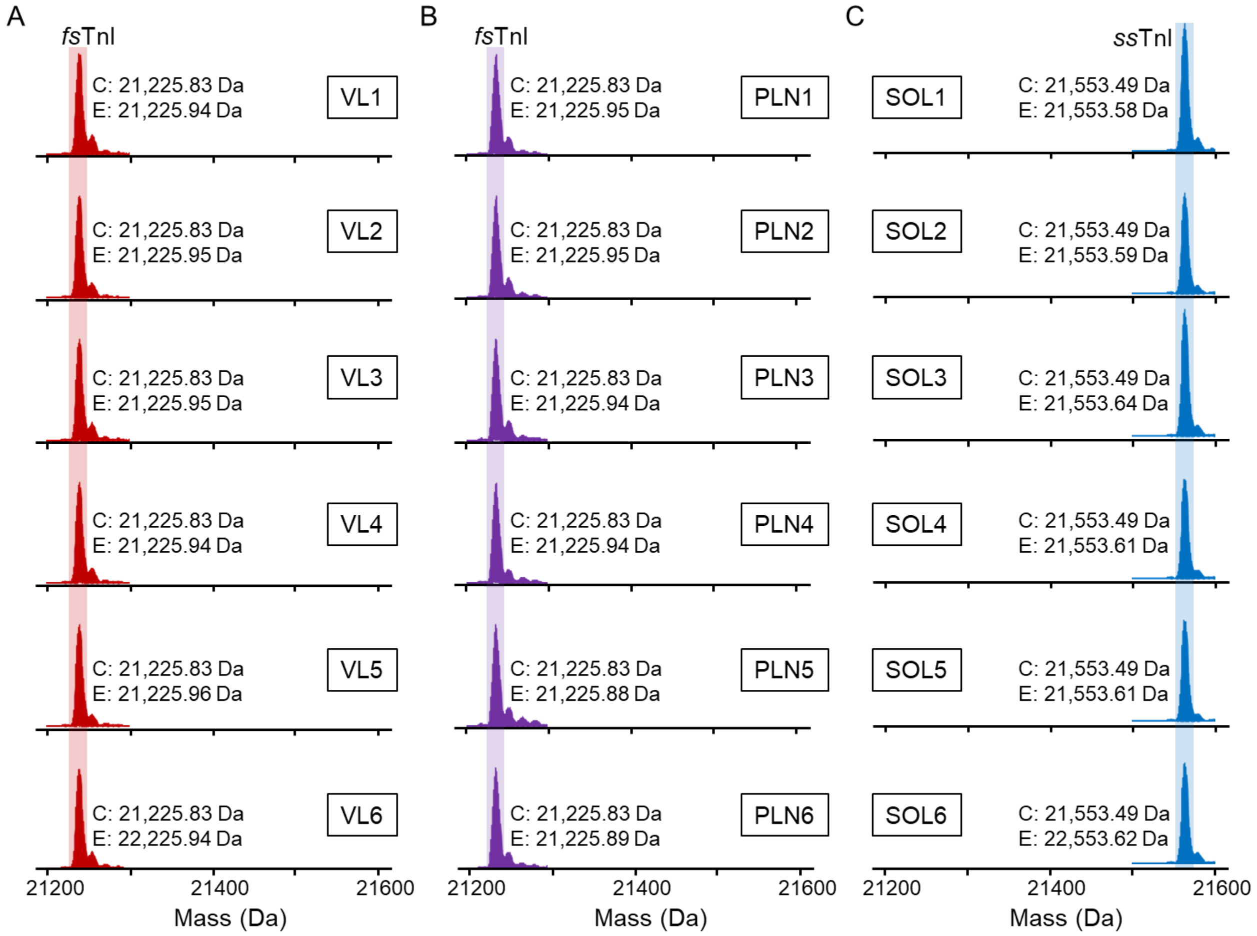


**Figure S13. Top-down proteomics of TnI isoforms from single muscle cells.** Deconvoluted mass spectra for TnI isoforms closely match predicted masses for SMFs obtained from **A)** VL (fsTnI), **B)** PLN (fsTnI), and **C)** SOL (ssTnI) muscles. Monoisotopic masses reported for the calculated (C) and experimental (E) masses of TnI isoforms. The UniProt accession number, P27768, was used for the fsTnI calculated mass. The UniProt accession number, Q9WUZ5, was used for the ssTnI calculated mass. The calculated and experimental masses are within a 10 ppm mass error. N=6 SMFs per group.

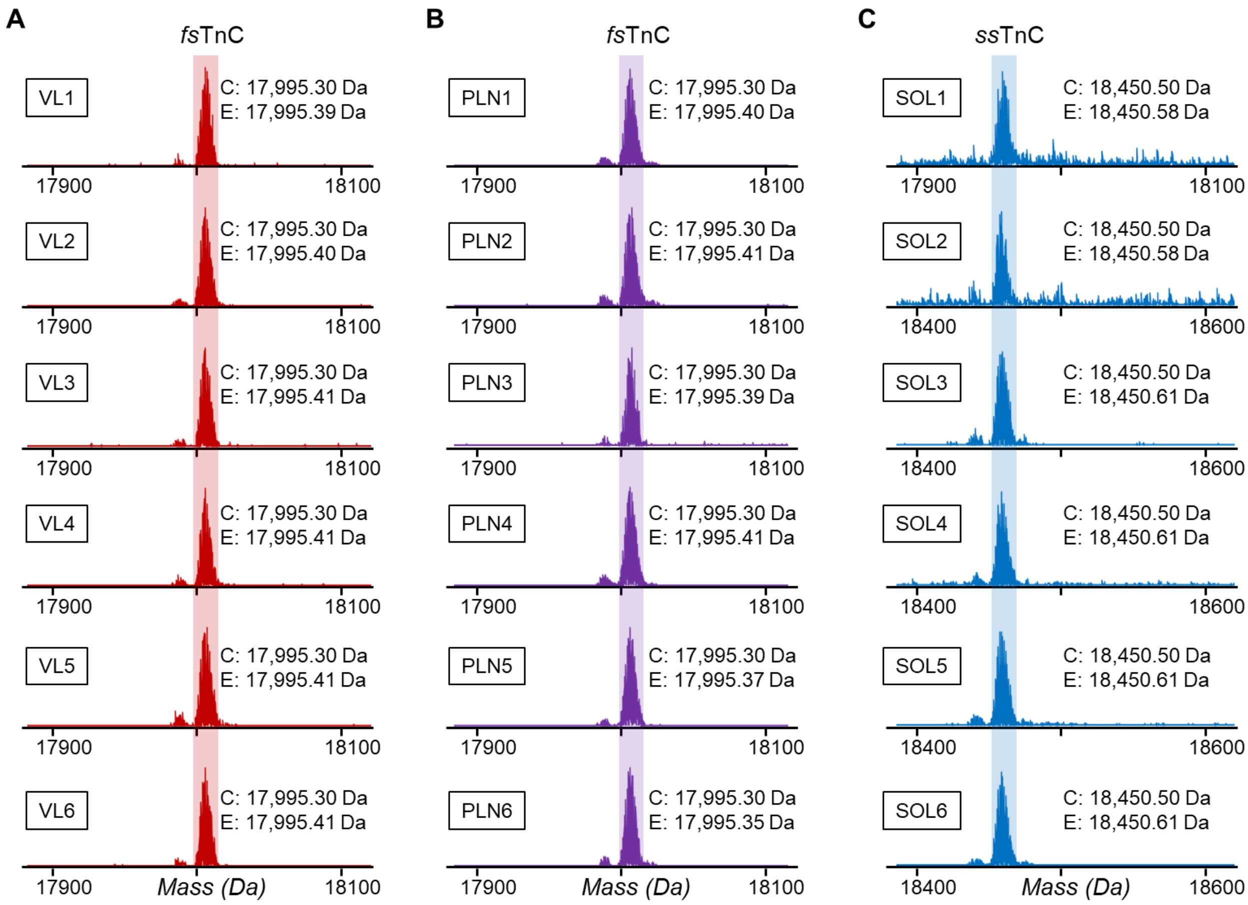


**Figure S14. Top-down proteomics of TnC isoforms from single muscle cells.** Deconvoluted mass spectra for TnC isoforms closely match predicted masses for SMFs obtained from **A)** VL (fsTnC), **B)** PLN (fsTnC), and **C)** SOL (ssTnC) muscles. Monoisotopic masses reported for the calculated (C) and experimental (E) masses of TnC isoforms. The UniProt accession number, Q304F3, was used for the fsTnC calculated mass. The UniProt accession number, Q4PP99, was used for the ssTnC calculated mass. The calculated and experimental masses are within a 10 ppm mass error. N=6 SMFs per group.

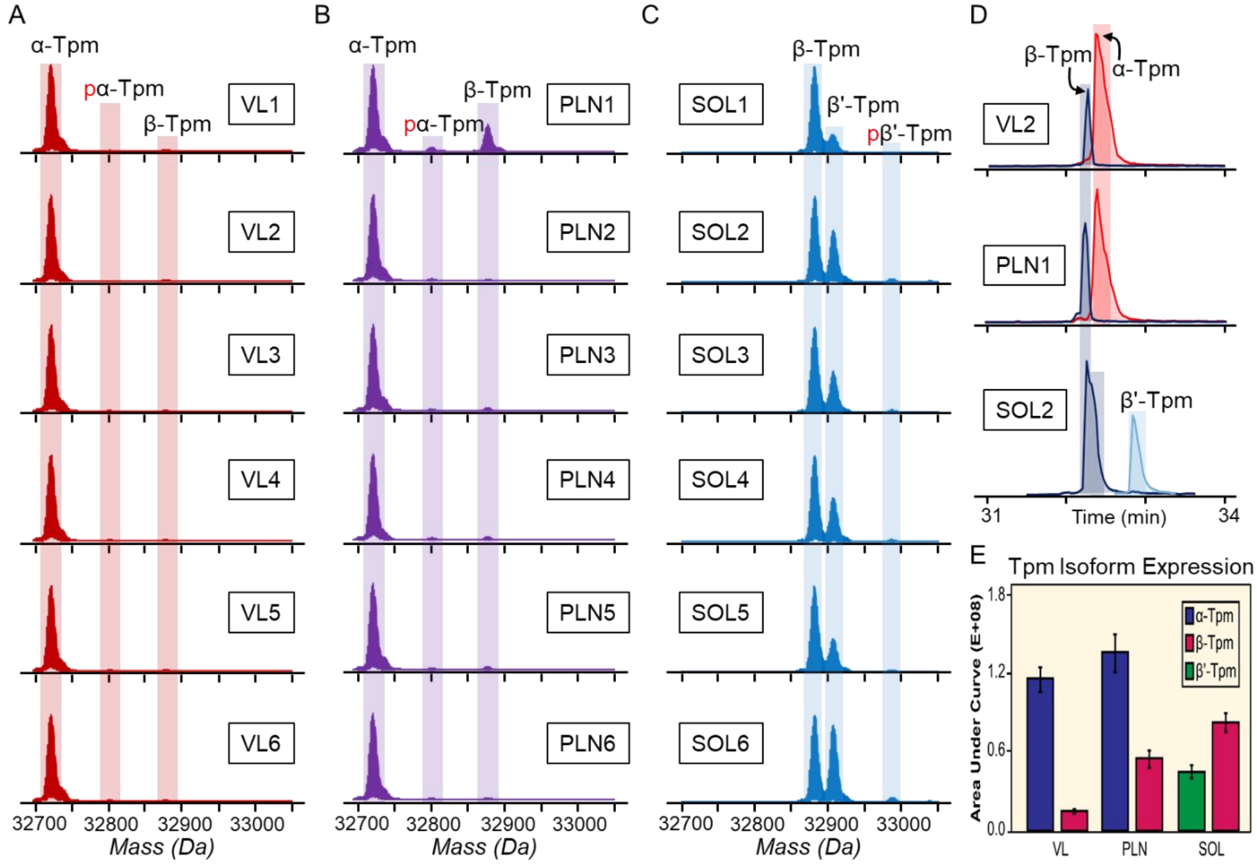


**Figure S15. Top-down proteomics provides a "bird's eye view" of Tpm isoforms and proteoforms from single muscle cells.** Deconvoluted mass spectra for TnC isoforms closely match predicted masses for SMFs obtained from **A)** VL, **B)** PLN, and **C)** SOL muscles.The VL and PLN fibers contained α-Tpm (UniProt Accession #P04692), mono-phosphorylated α-Tpm and β-Tpm (UniProt Accession #P58775). The SOL fibers contained β-Tpm, β'-Tpm and mono-phosphorylated β'-Tpm (5, 7). Mono-phosphorylation is indicated by red "p". **D)** Extracted ion chromatograms for the predominate Tpm isoforms in a representative VL, PLN, and SOL fiber **E)** Area under the curve quantitation of the various Tpm isoforms found across the fiber samples. Isoform expression is heterogenous across the single fiber samples. N=6 SMFs per group.

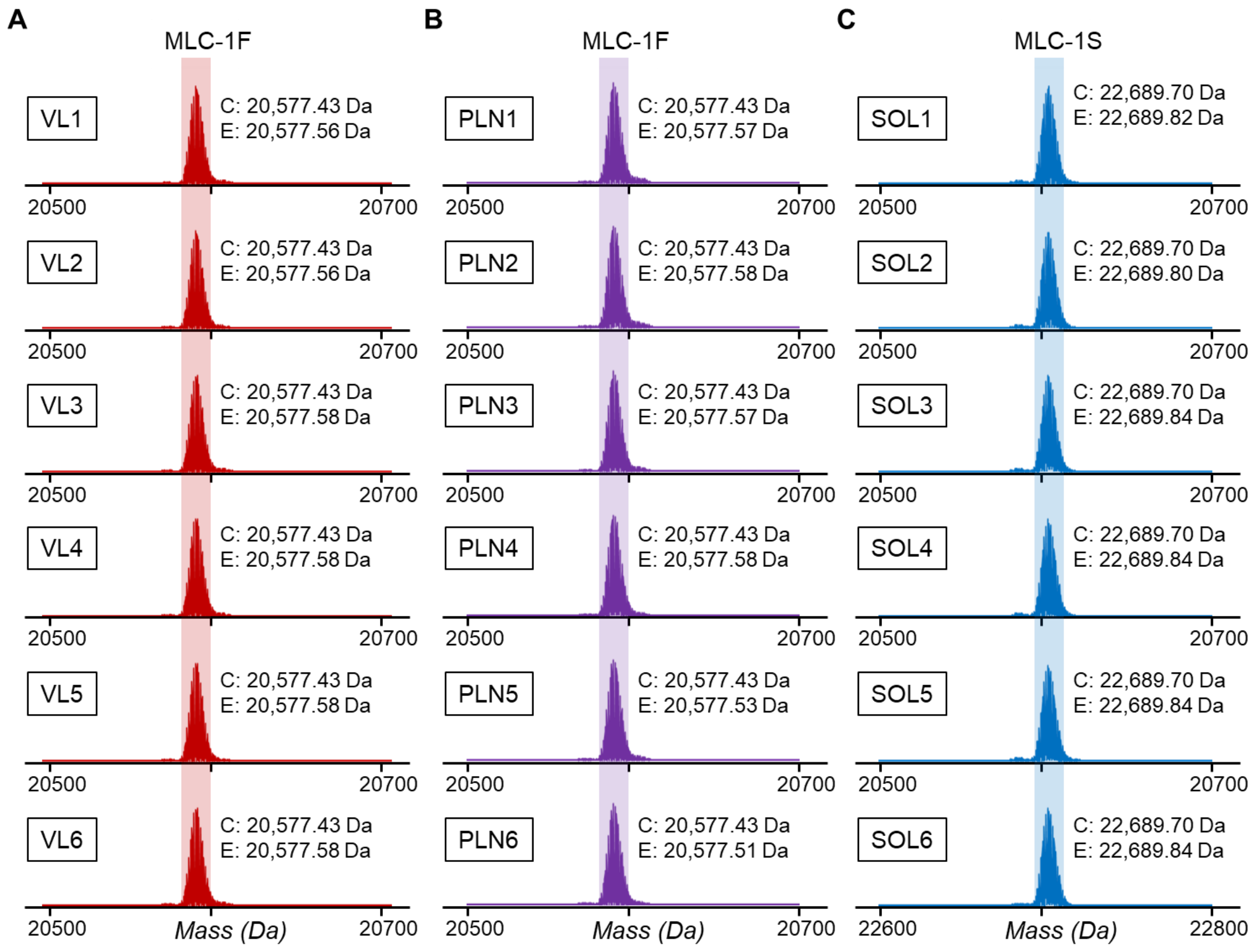


**Figure S16. Top-down proteomics of MLC1 isoforms from single muscle cells.** Deconvoluted mass spectra for MLC1 isoforms closely match predicted masses for SMFs obtained from **A)** VL (MLC-1F), **B)** PLN (MLC-1F), and **C)** SOL (MLC-1S) muscles. Monoisotopic masses reported for the calculated (C) and experimental (E) masses of MLC-1 isoforms. The UniProt accession number, P02600, was used for the MLC-1F calculated mass. The UniProt accession number, D3ZHA7, was used for the MLC-1S calculated mass. The calculated and experimental masses are within a 10 ppm mass error. N=6 SMFs per group.

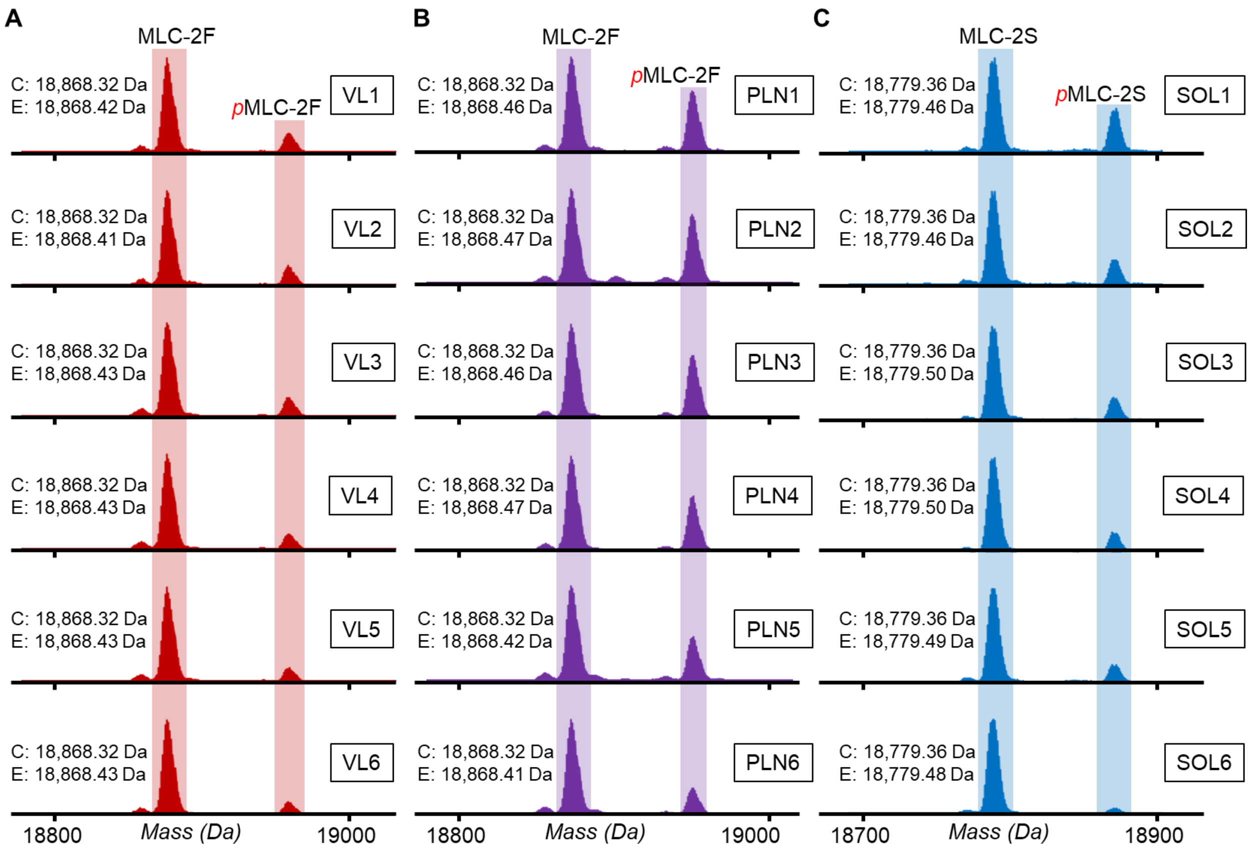


**Figure S17. Top-down proteomics provides a "bird's eye view" of MLC2 isoforms and proteoforms from single muscle cells.** Deconvoluted mass spectra for MLC2 isoforms closely match predicted masses for SMFs obtained from **A)** VL (MLC-2F), **B)** PLN (MLC-2F), and **C)** SOL (MLC-2S) muscles. Monoisotopic masses reported for the calculated (C) and experimental (E) masses of MLC-2 isoforms. The UniProt accession number, P04466, was used for the MLC-2F calculated mass. The UniProt accession number, P08733, was used for the MLC-2S calculated mass. The calculated and experimental masses are within a 10 ppm mass error. Mono-phosphorylation is indicated by red "p". N=6 SMFs per group.

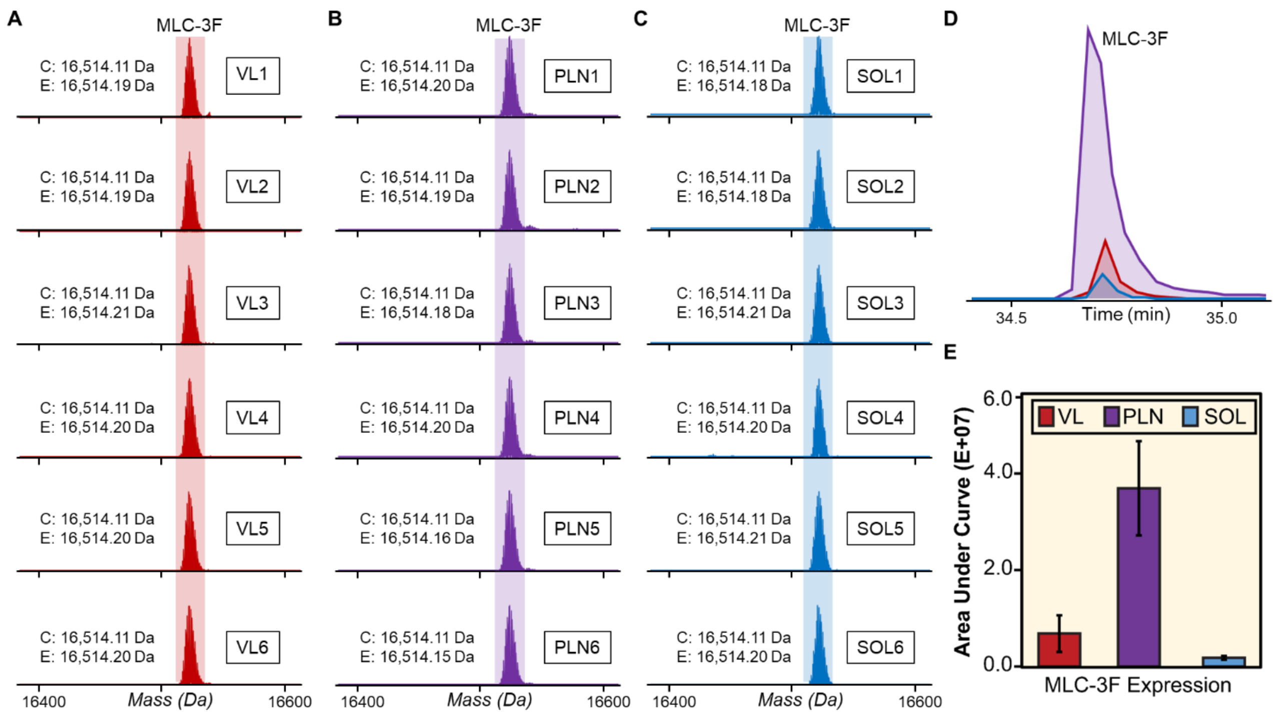


**Figure S18. Top-down proteomics of MLC3 isoforms from single muscle cells.** MLC-3F is found in each of the single fiber samples but most predominantly in PLN fibers. **A-C)** Deconvoluted mass spectra of MLC-3F from SMFs obtained from **A)** VL **B)** PLN and **C)** SOL muscles. Monoisotopic masses reported for the calculated (C) and experimental (E) masses of MLC-3F. The UniProt accession number, P02600-2, was used for the MLC-3F calculated mass. The calculated and experimental masses are within a 10 ppm mass error. **D)** EICs (top 5 most abundant ions) for MLC-3F obtained from SMFs from VL, PLN, and SOL muscles. **E)** AUC of the MLC-3F EICs generated from for the fiber samples. Error bars reported are the standard deviation. N=6 SMFs per group.

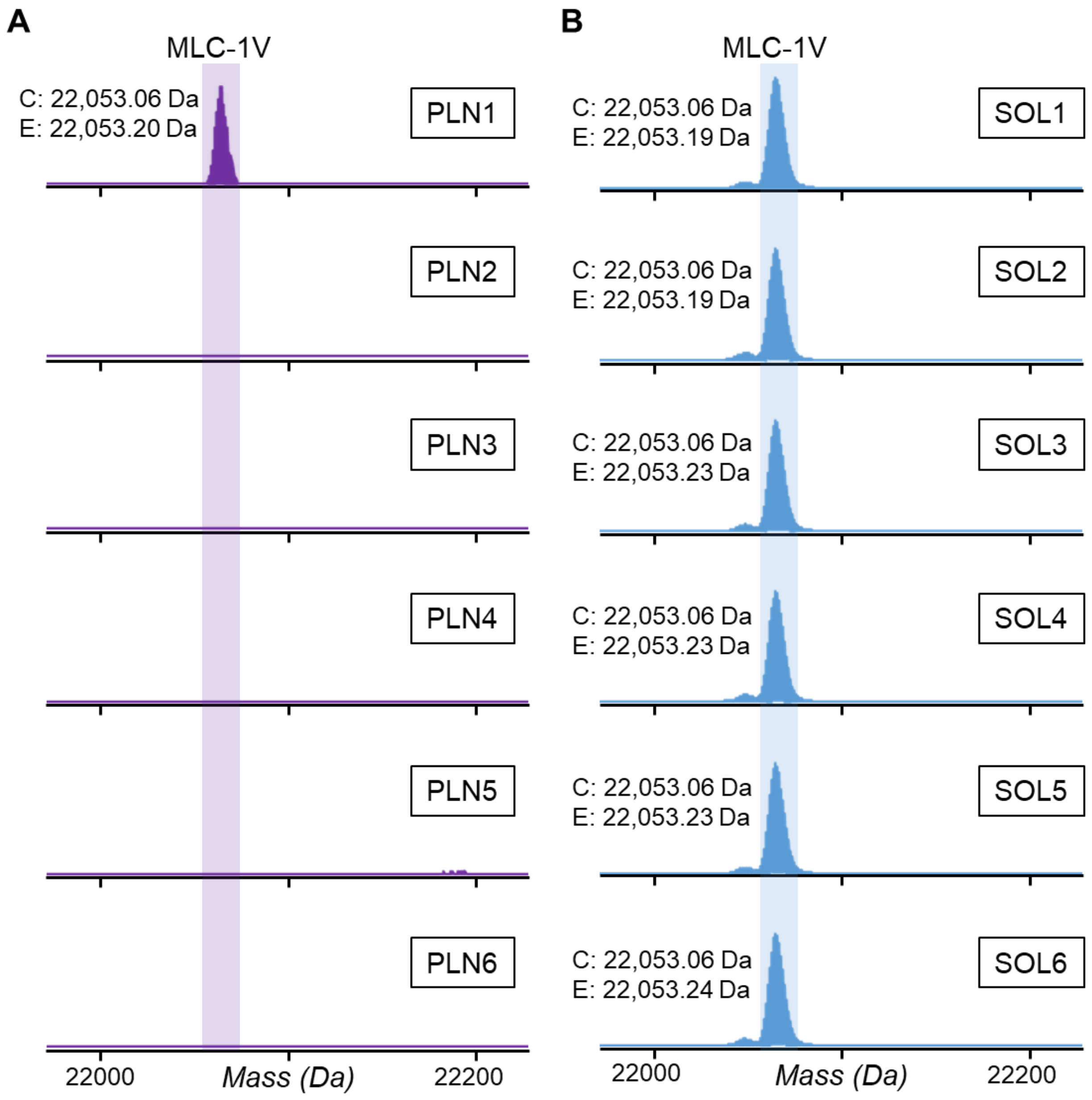


**Figure S19. Top-down proteomics of MLC-1V isoforms from single muscle cells.** Deconvoluted mass spectra for the MLC-1V reveals expression in **A)** unexpextedly one PLN fiber (fast-twitch) and **B)** all SOL fibers (slow-twitch). Monoisotopic masses reported for the calculated (C) and experimental (E) masses of MLC-1V. The UniProt accession number, P16409, was used for the MLC-1V calculated mass. The calculated and experimental masses are within a 10 ppm mass error. N=6 SMFs per group.

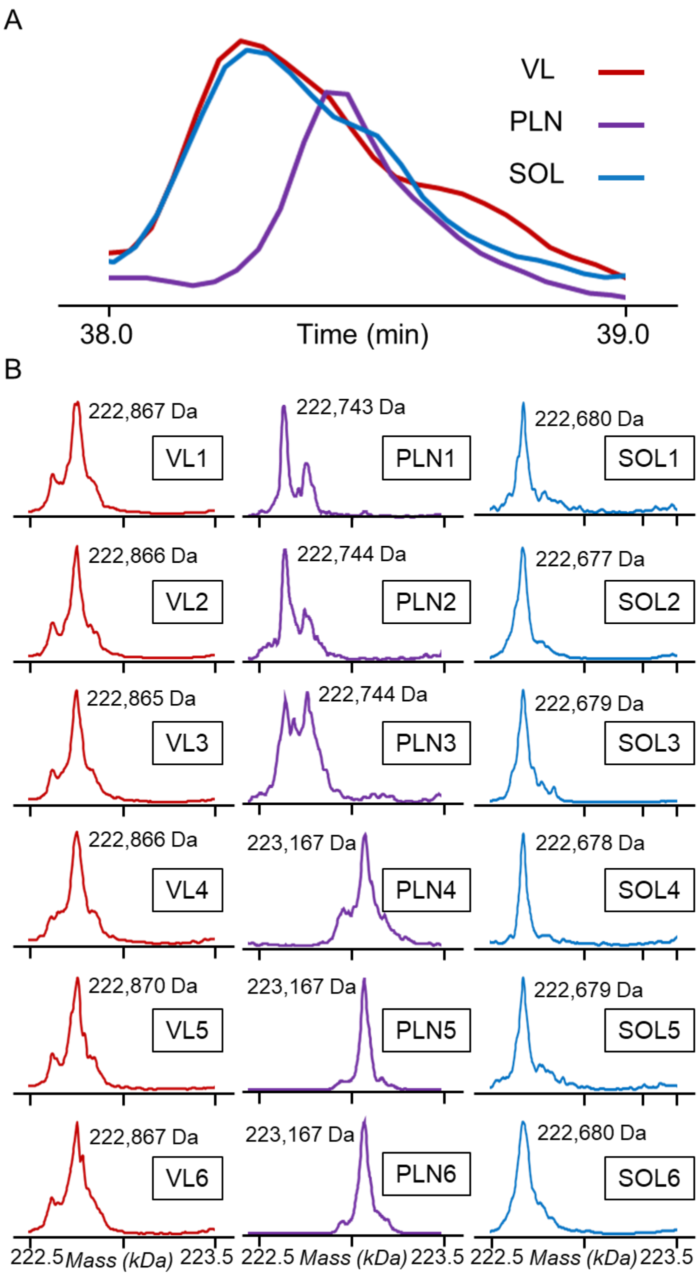


**Figure S20. Detection of MyHC isoforms from single muscle cells A)** EICs using the top 7 most abundant ions of MyHC isoforms from representative SMFs obtained from VL, PLN, and SOL muscles **B)** Deconvoluted mass spectra for myosin heavy chain (MyHC) isoforms from SMFs obtained from VL**,** PLN, and SOL muscles. Most abundant mass reported for the experimental (E) mass of α-MHC isoforms using the Sum Peak algorithm. The maximum entropy deconvolution parameters were set to low resolution (10,000) because the charge states of the MS1 spectra were not fully resolved. N=6 SMFs per group.

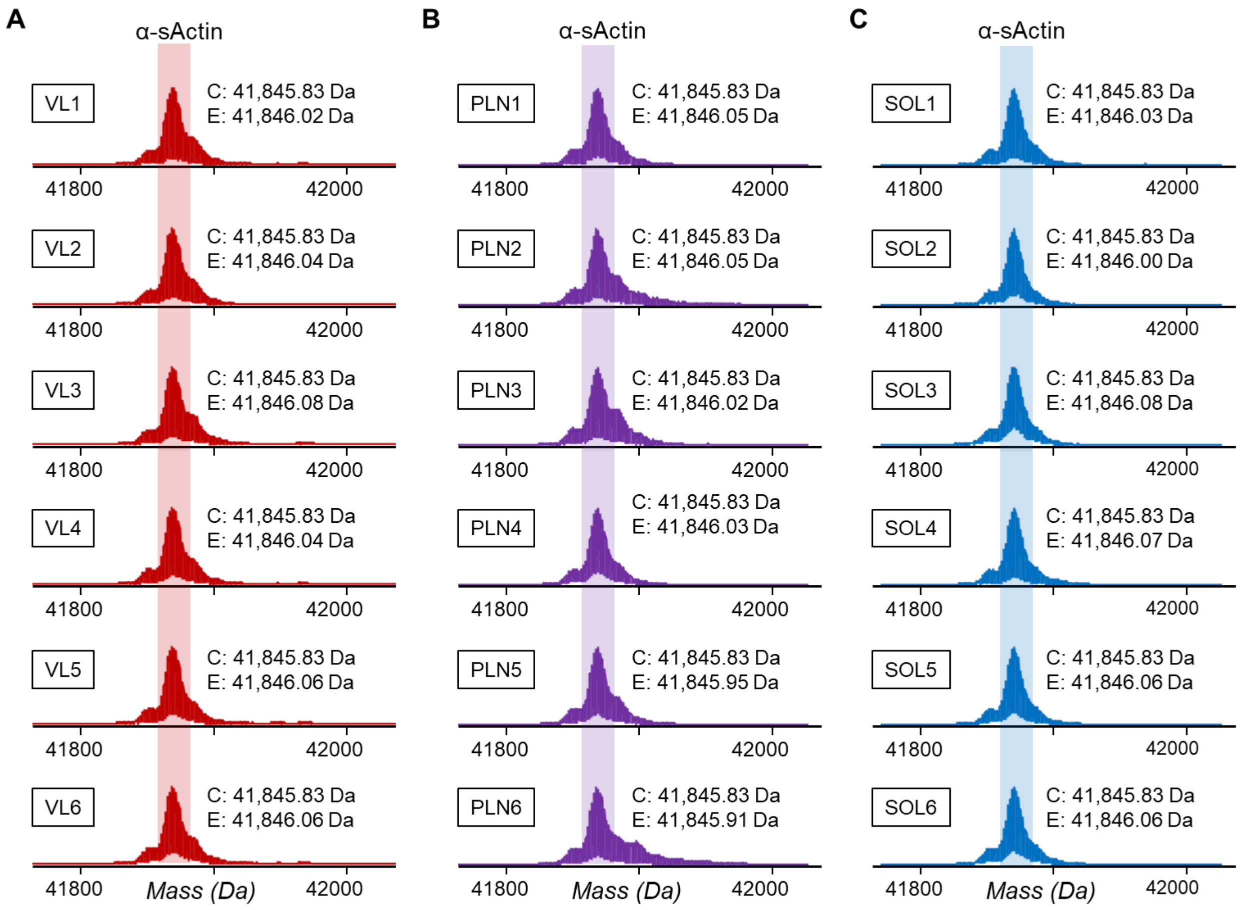


**Figure S21. Top-down proteomics of alpha skeletal actin proteoforms from single muscle cells.** Deconvoluted mass spectra for α-sActin closely match predicted masses for SMFs obtained from **A)** VL, **B)** PLN, and **C)** SOL muscles. Monoisotopic masses reported for the calculated (C) and experimental (E) masses of α-sActin. The UniProt accession number, P68136, was used for the α-sActin calculated mass. The calculated and experimental masses are within a 10 ppm mass error. N=6 SMFs per group.

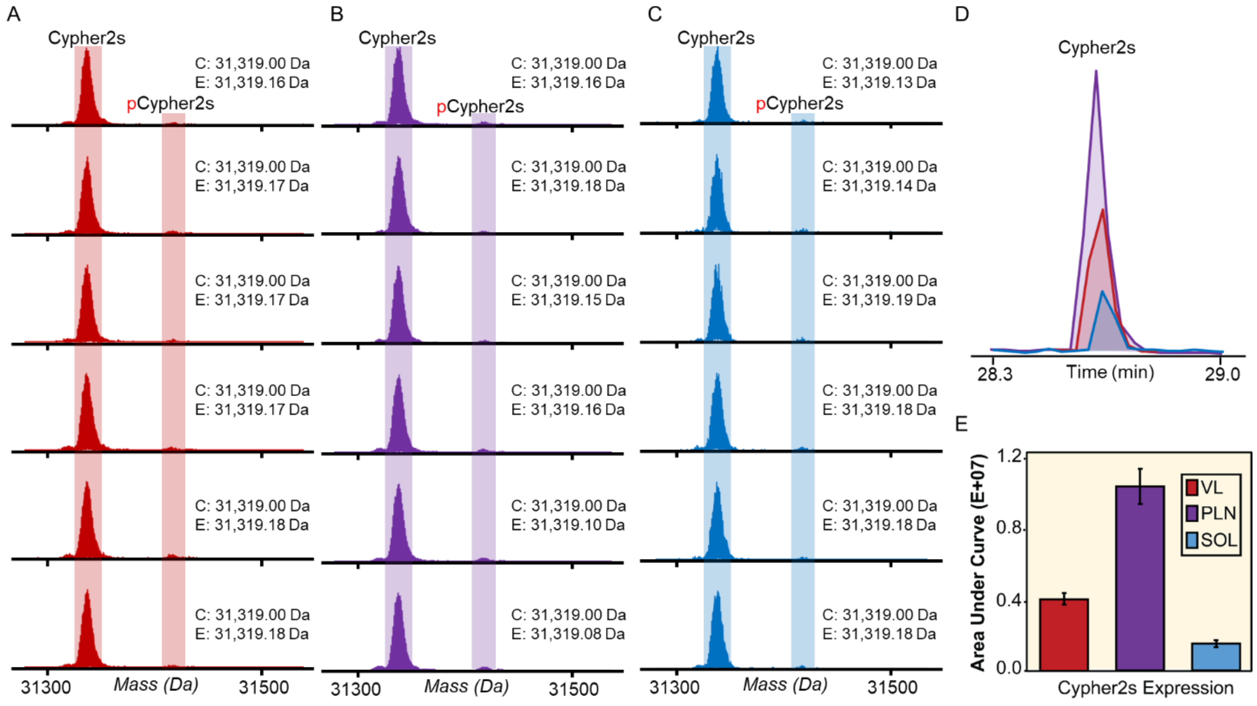


**Figure S22. Top-down proteomics provides a "bird's eye view" of Cypher2s proteoforms from single muscle cells.** Cypher2s proteoforms are found in each of the single fiber samples but most predominantly in PLN fibers. **A-C)** Deconvoluted mass spectra of Cypher2s from SMFs obtained from **A)** VL **B)** PLN and **C)** SOL muscles. Monoisotopic masses reported for the calculated (C) and experimental (E) masses of Cypher2s. The UniProt accession number, Q9JKS4, was used for the Cypher2s calculated mass. The calculated and experimental masses are within a 10 ppm mass error. Mono-phosphorylation is indicated by red "p". **D)** Total phosphorylation level, Ptotal (mol Pi/mol protein), for Cypher2s from VL, PLN, and SOL fibers (n=6 per tissue). **E)** AUC of the Cypher2s EICs generated from for the fiber samples (n=6). Error bars reported are the standard deviation.

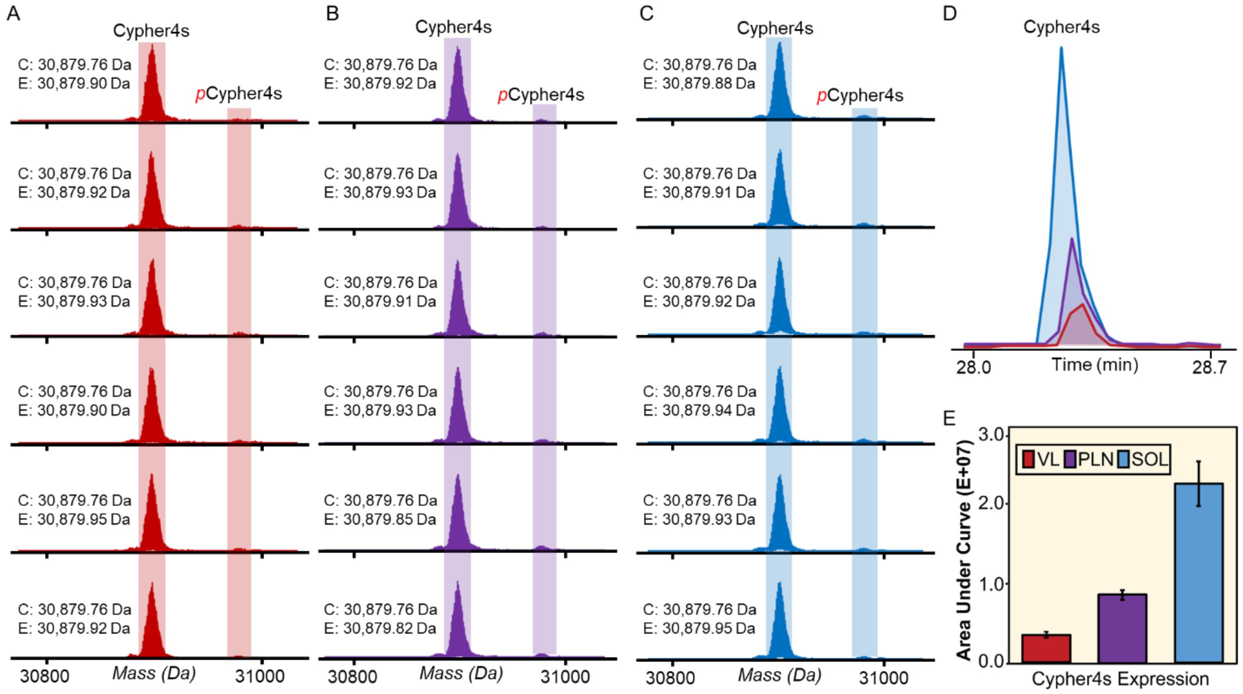


**Figure S23. Top-down proteomics provides a "bird's eye view" of Cypher4s proteoforms from single muscle cells.** Cypher4s proteoforms are found in each of the single fiber samples but most predominantly in SOL fibers. **A-C)** Deconvoluted mass spectra of Cypher4s from SMFs obtained from **A)** VL **B)** PLN and **C)** SOL muscles. Monoisotopic masses reported for the calculated (C) and experimental (E) masses of Cypher4s. The UniProt accession number, Q5XIG1, was used for the Cypher4s calculated mass. The calculated and experimental masses are within a 10 ppm mass error. Mono-phosphorylation is indicated by red "p". **D)** Total phosphorylation level, Ptotal (mol Pi/mol protein), for Cypher4s from VL, PLN, and SOL fibers (n=6 per tissue) reveals similar levels of phosphorylation across all fiber samples. **E)** AUC of all fiber samples. EICs generated from for the fiber samples (n=6). Error bars reported are the standard deviation.

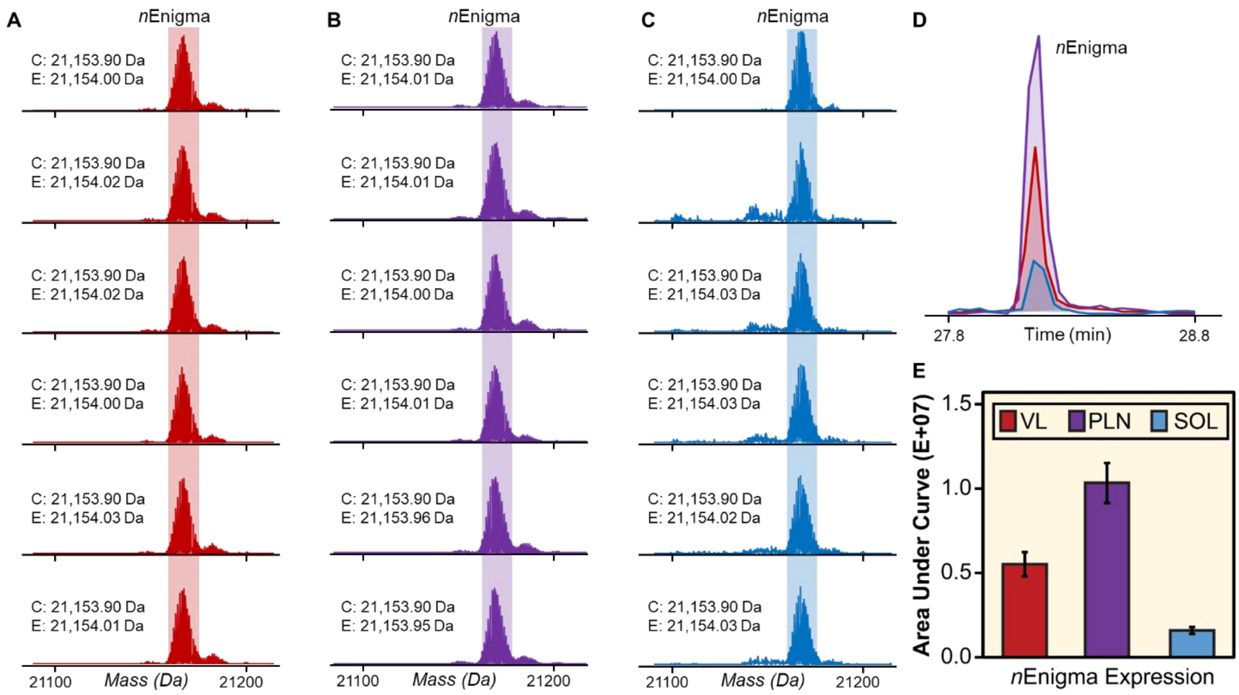


**Figure S24. Top-down proteomics provides a "bird's eye view" of nEnigma proteoforms from single muscle cells.** Z-disk protein, nEnigma, is found in each of the SMFs but most predominantly in PLN fibers. **A-C)** Deconvoluted mass spectra of nEnigma from SMFs obtained from **A)** VL **B)** PLN and **C)** SOL muscles. Monoisotopic masses reported for the calculated (C) and experimental (E) masses of nEnigma (7). The calculated and experimental masses are within a 10 ppm mass error. Mono-phosphorylation is indicated by red "p". **D)** EICs (top 5 most abundant ions) for nEnigma obtained from SMFs from VL, PLN, and SOL muscles. **E)** AUC for all each of the nEnigma EICs generated from for the fiber samples (n=6). Error bars reported are the standard deviation.

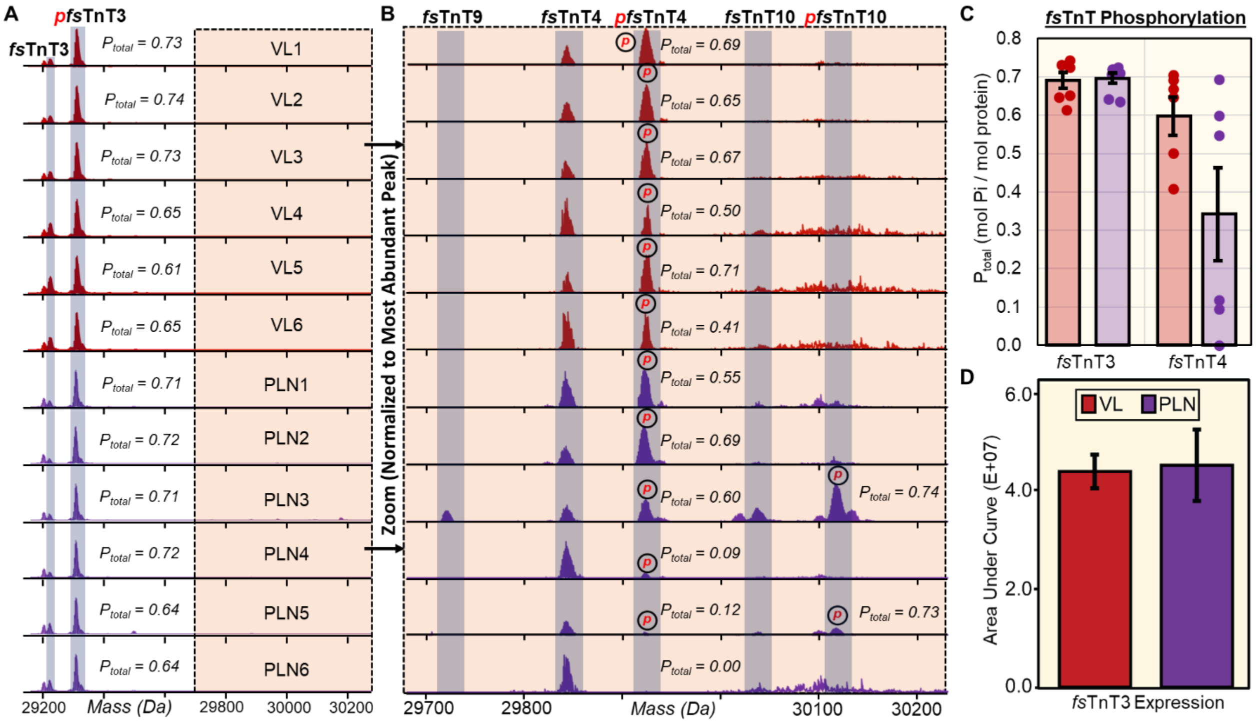


**Figure S25. Altered phosphorylation of low abundance fast skeletal troponin T (fsTnT) proteoforms from fiber-to-fiber. A)** Representative deconvoluted mass spectra of fast skeletal troponin 3 (fsTnT3) from SMFs isolated from fast-twitch VL (red) and PLN (purple) muscles. All of the spectra are normalized to the most intense peak. Mono-phosphorylation is denoted with red "p". **B)** Zoom-in on 29,700 Da – 30,200 Da region of spectra then further normalization to most intense peak further displays several low abundance fast skeletal troponin isoforms (fsTnT4, fsTnT9, fsTnT10) in SMFs from fast-twitch VL (red) and PLN (purple) muscles. All of the spectra are normalized to 50,000 intensity units. Mono-phosphorylation is denoted with red "p". **C)** Total phosphorylation (Ptot) calculated as mol Pi/mol protein for fsTnT3 and fsTnT4 from each fiber (n=6). **D)** Extracted ion chromatograms (EICs; top 5 most abundant ions) of fsTnT3 were made and the area under the curve was integrated to calculate the fsTnT3 expression.

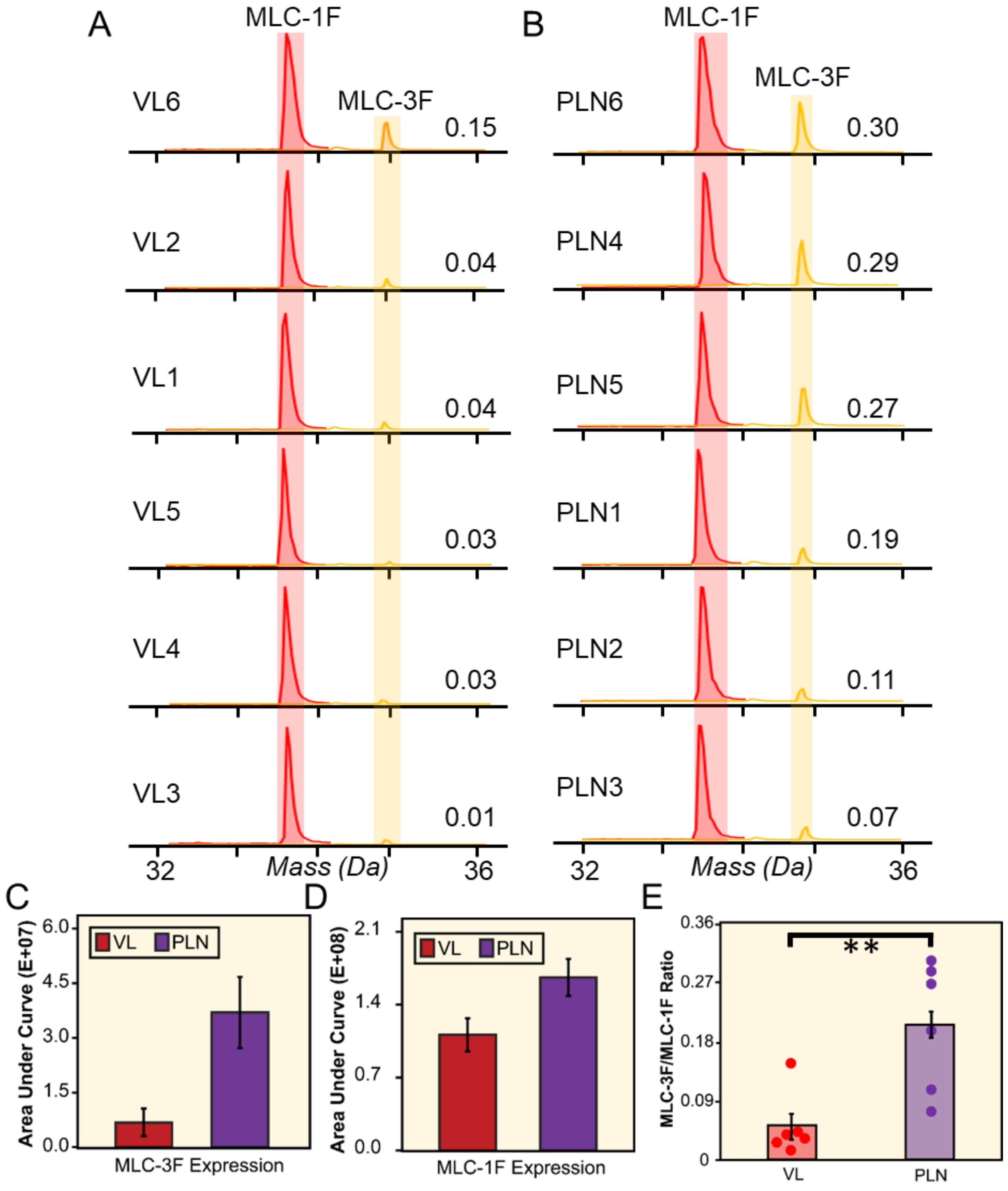


**Figure S26. The ratio of MLC1 to MLC3 isoform expression varies across VL and PLN SMFs (fast-twitch). A-B)** EICs (top five most abundant ions) of MLC-1F and MLC-3F from **A)** VL and **B)** PLN fiber samples. **C-D)** Area under the curve of the **C)** MLC-3F and **D)** MLC-1F EICs generated from for the VL and PLN fiber samples. **E)** The ratio of MLC-3F:MLC-1F reveals that there are altered levels of the ratio from fiber-to-fiber, particularly in the PLN fiber samples. ** indicates a p-value <0.01 by Student's t-test.

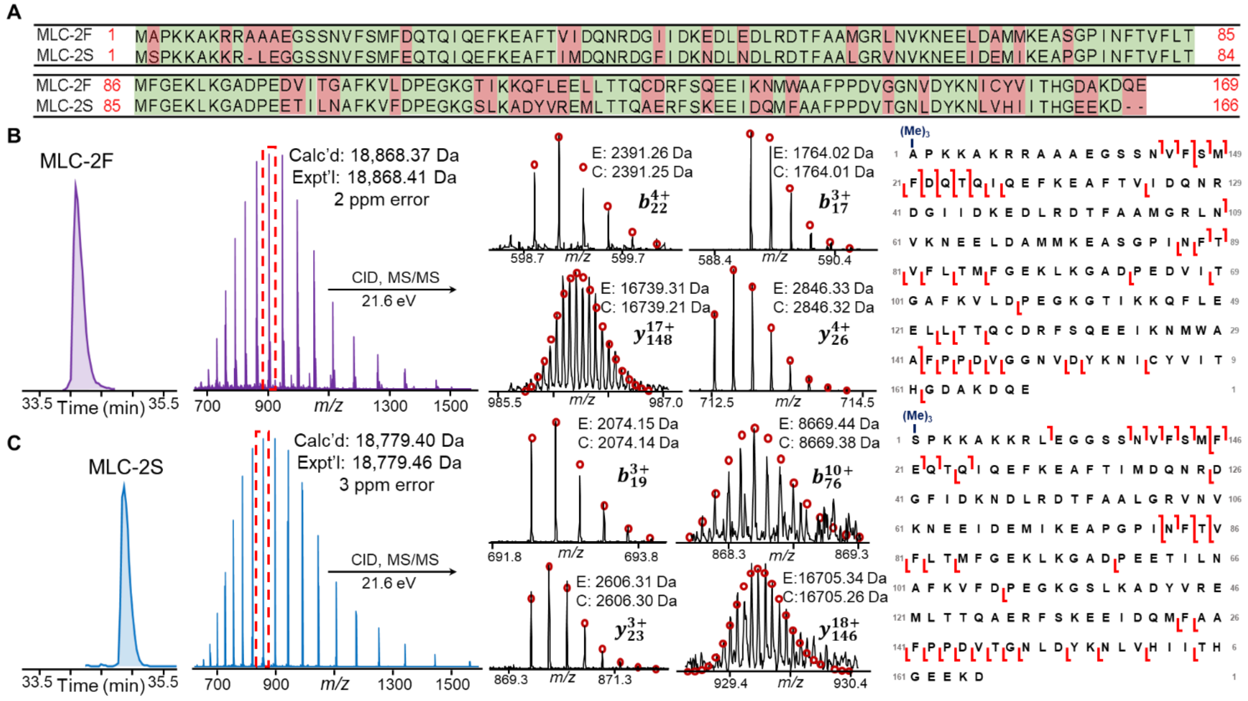


**Figure S27. Top-down MS characterization of MLC-2 isoforms from SMFs. A**) Sequence alignment of the MLC-2F and MLC-2S. Red indicates no residues shared in common between the isoforms and green indicates sequence homology between the isoforms. **B-C)** Online LC–MS/MS of **B)** MLC-2F and **C)** MLC-2S. Representative EIC for each proteoform were generated from the top five most abundant ions, which was averaged to show the MS1 spectra. Monoisotopic masses reported for the calculated (C) and experimental (E) masses of MLC-2F and MLC-2S. The UniProt accession number, P04466, was used for the MLC-2F calculated mass. The UniProt accession number, P08733, was used for the MLC-2S calculated mass. The calculated and experimental masses are within a 25 ppm mass error. Precursor ions were selected for online collisionally activated dissociation resulting in sequence informative b and y ions. The sequence table with N-terminal modification shows the characterization of MLC-2 isoforms.

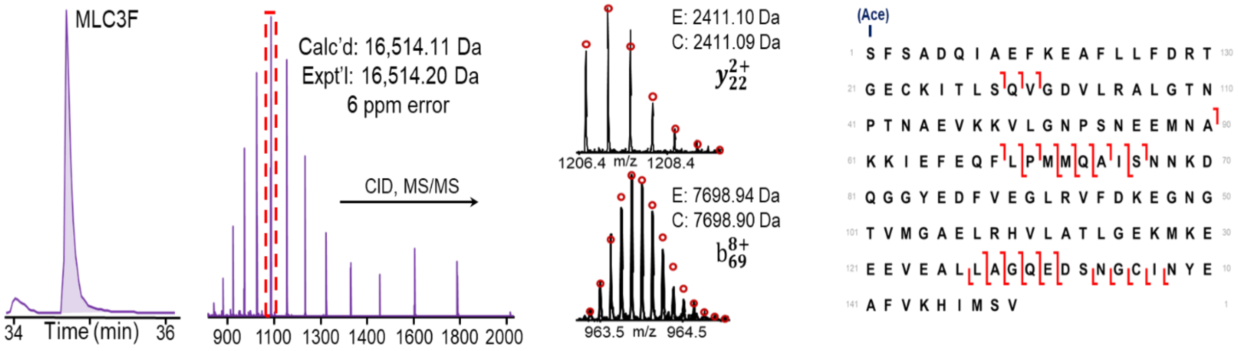


**Figure S28. Top-down MS characterization of MLC-3F proteoforms from SMFs.** Online LC–MS/MS of MLC-3F. Representative EIC was generated from the top five most abundant ions, which was averaged to show the MS1 spectra. Monoisotopic masses reported for the calculated (C) and experimental (E) masses of MLC-3F. The UniProt accession number, P02600-2, was used for the MLC-3F calculated mass. The calculated and experimental masses are within a 25 ppm mass error. Precursor ions were selected for online collisionally activated dissociation resulting in sequence informative b and y ions. The sequence table with N-terminal modification shows the characterization of MLC-3F isoforms.

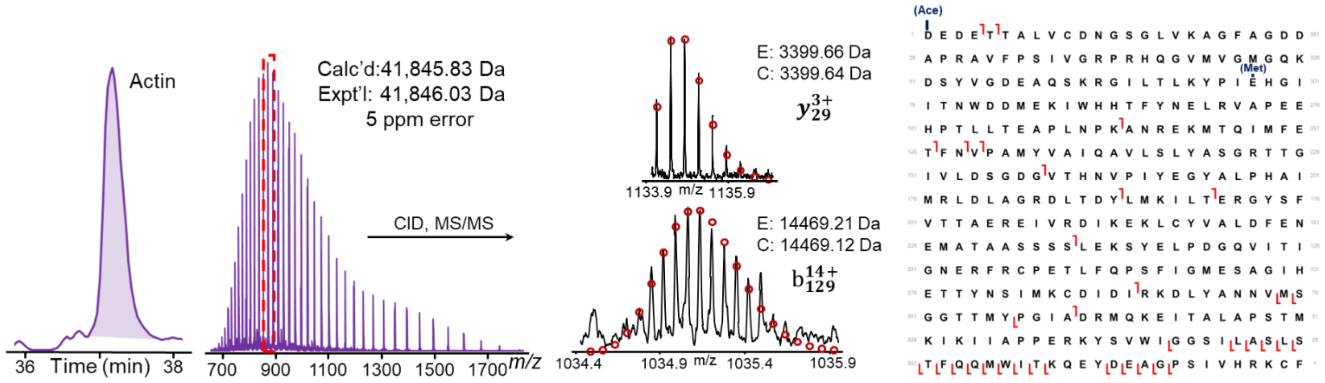


**Figure S29. Top-down MS characterization of α-sActin proteoforms from SMFs.** Online LC–MS/MS of α-sActin. Representative EIC was generated from the top five most abundant ions, which was averaged to show the MS1 spectra. Monoisotopic masses reported for the calculated (C) and experimental (E) masses of α-sActin. The UniProt accession number, P68136, was used for the α-sActin calculated mass. The calculated and experimental masses are within a 25 ppm mass error. Precursor ions were selected for online collisionally activated dissociation resulting in sequence informative b and y ions. The sequence table with N-terminal modification and methylation shows the characterization of α-sActin isoforms.

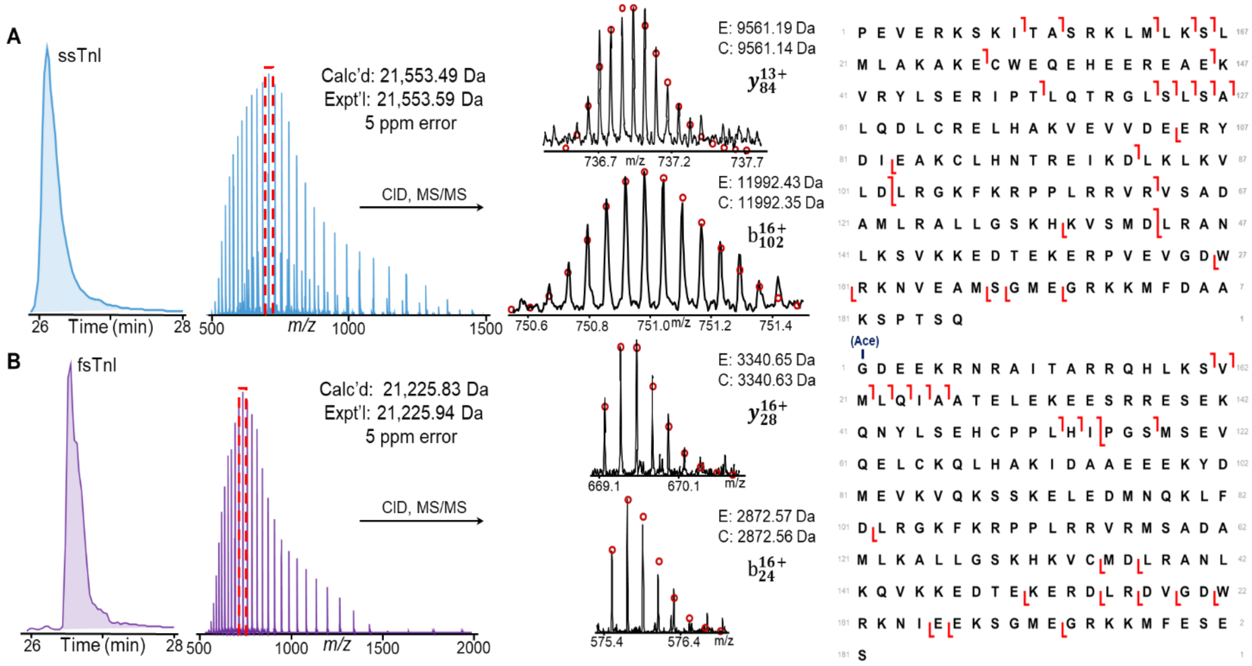


**Figure S30. Top-down MS characterization of TnI isoforms from SMFs. A-B)** Online LC–MS/MS of **A)** ssTnI (SOL) and **B)** fsTnI (PLN). Representative EIC for each isoform were generated from the top five most abundant ions, which was averaged to show the MS1 spectra. Monoisotopic masses reported for the calculated (C) and experimental (E) masses of ssTnI and fsTnI. The UniProt accession number, Q9WUZ5, was used for the ssTnI calculated mass.The UniProt accession number, P27768, was used for the fsTnI calculated mass. The calculated and experimental masses are within a 25 ppm mass error. Precursor ions were selected for online collisionally activated dissociation resulting in sequence informative b and y ions. The sequence table with N-terminal modification shows the characterization of TnI isoforms.

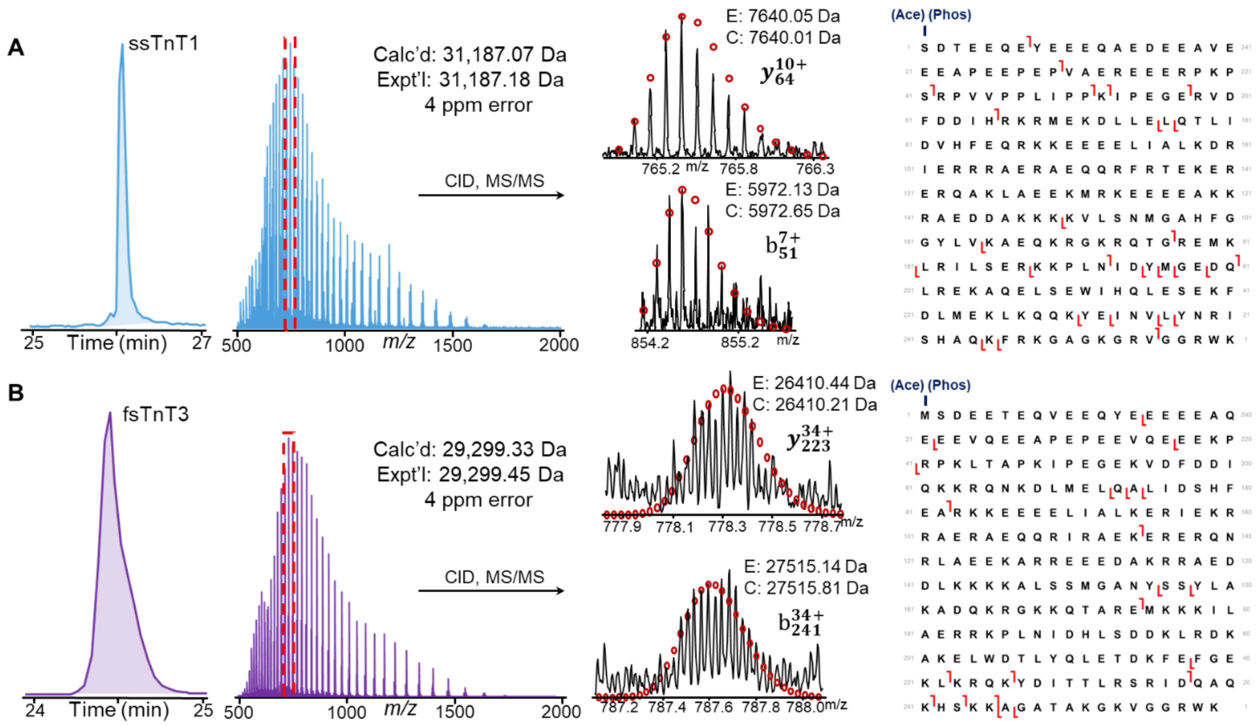


**Figure S31. Top-down MS characterization of TnT isoforms from SMFs. A-B)** Online LC–MS/MS of **A)** ssTnT1 (SOL) and **B)** fsTnI (PLN). Representative EIC for each isoform were generated from the top five most abundant ions, which was averaged to show the MS1 spectra. Monoisotopic masses reported for the calculated (C) and experimental (E) masses of ssTnT1 and fsTnT3. The UniProt accession number, P7TNB2, was used for the ssTnT1 calculated mass. The UniProt accession number, P09739, was used for the fsTnT3 calculated mass. The calculated and experimental masses are within a 25 ppm mass error. Precursor ions were selected for online collisionally activated dissociation resulting in sequence informative b and y ions. The sequence table with N-terminal modification shows the characterization of TnT isoforms.

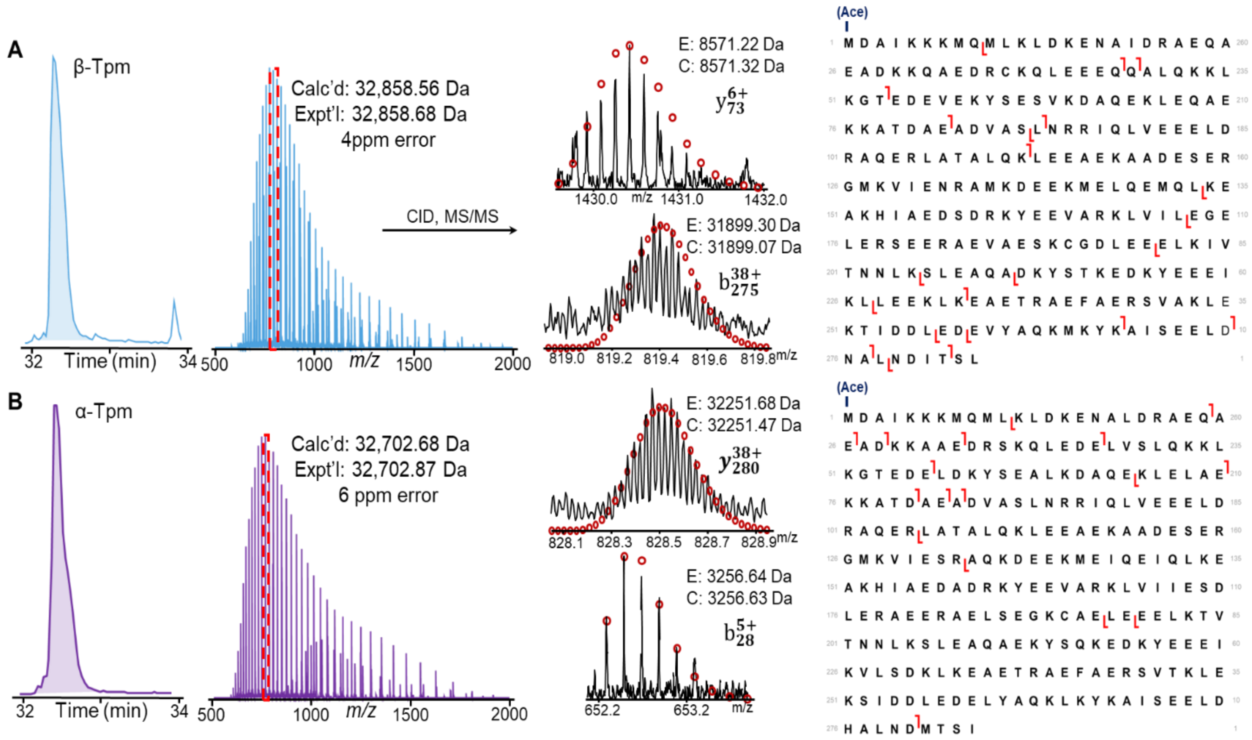


**Figure S32. Top-down MS characterization of Tpm isoforms from SMFs. A-B)** Online LC–MS/MS of **A)** β-Tpm and **B)** α-Tpm. Representative EIC for each isoform were generated from the top five most abundant ions, which was averaged to show the MS1 spectra. Monoisotopic masses reported for the calculated (C) and experimental (E) masses of β-Tpm and α-Tpm. The UniProt accession number, P04692, was used for the α-Tpmcalculated mass. The UniProt accession number, P58775, was used for the β-Tpm calculated mass. The calculated and experimental masses are within a 25 ppm mass error. Precursor ions were selected for online collisionally activated dissociation resulting in sequence informative b and y ions. The sequence table with N-terminal modification shows the characterization of Tpm isoforms.

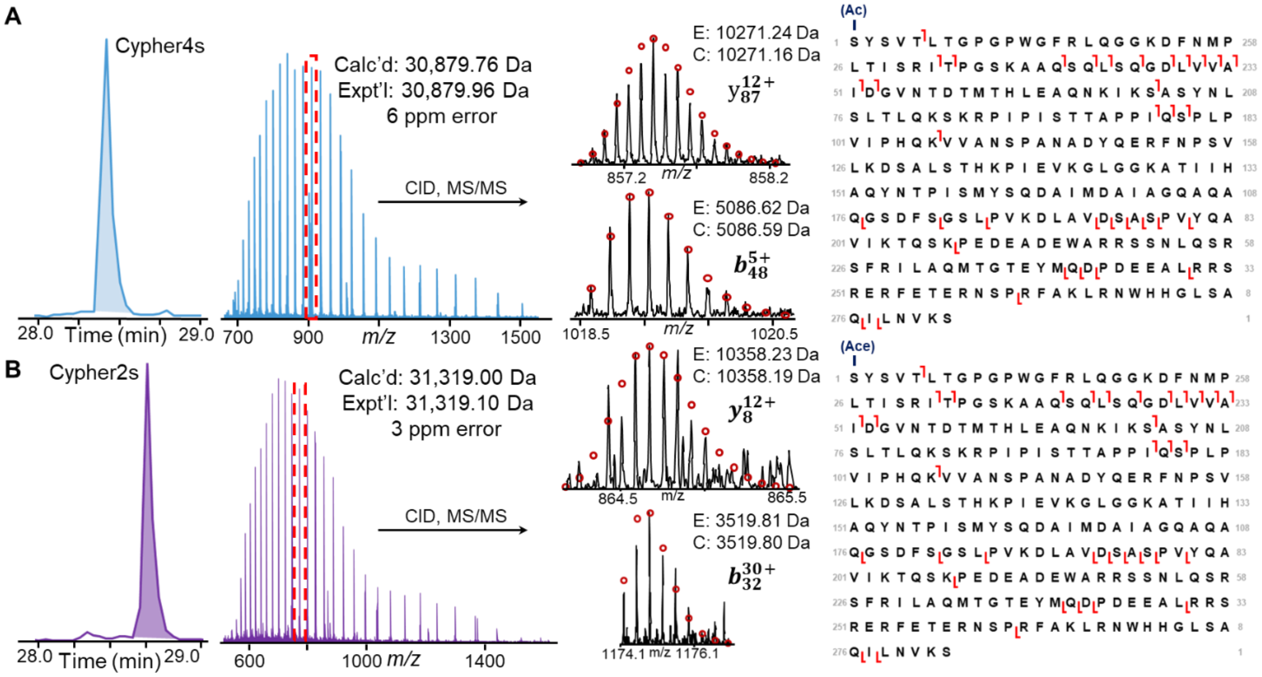


**Figure S33. Top-down MS characterization of Cypher isoforms from SMFs. A-B)** Online LC–MS/MS of **A)** Cypher4s and **B)** Cypher2s. Representative EIC for each isoform were generated from the top five most abundant ions, which was averaged to show the MS1 spectra. Monoisotopic masses reported for the calculated (C) and experimental (E) masses of Cypher4s and Cypher2s. The UniProt accession number, Q5XIG1, was used for the Cypher4s calculated mass. The UniProt accession number, Q9JKS4, was used for the Cypher2s calculated mass. The calculated and experimental masses are within a 25 ppm mass error. Precursor ions were selected for online collisionally activated dissociation resulting in sequence informative b and y ions. The sequence table with N-terminal modification shows the characterization of Cypher isoforms.

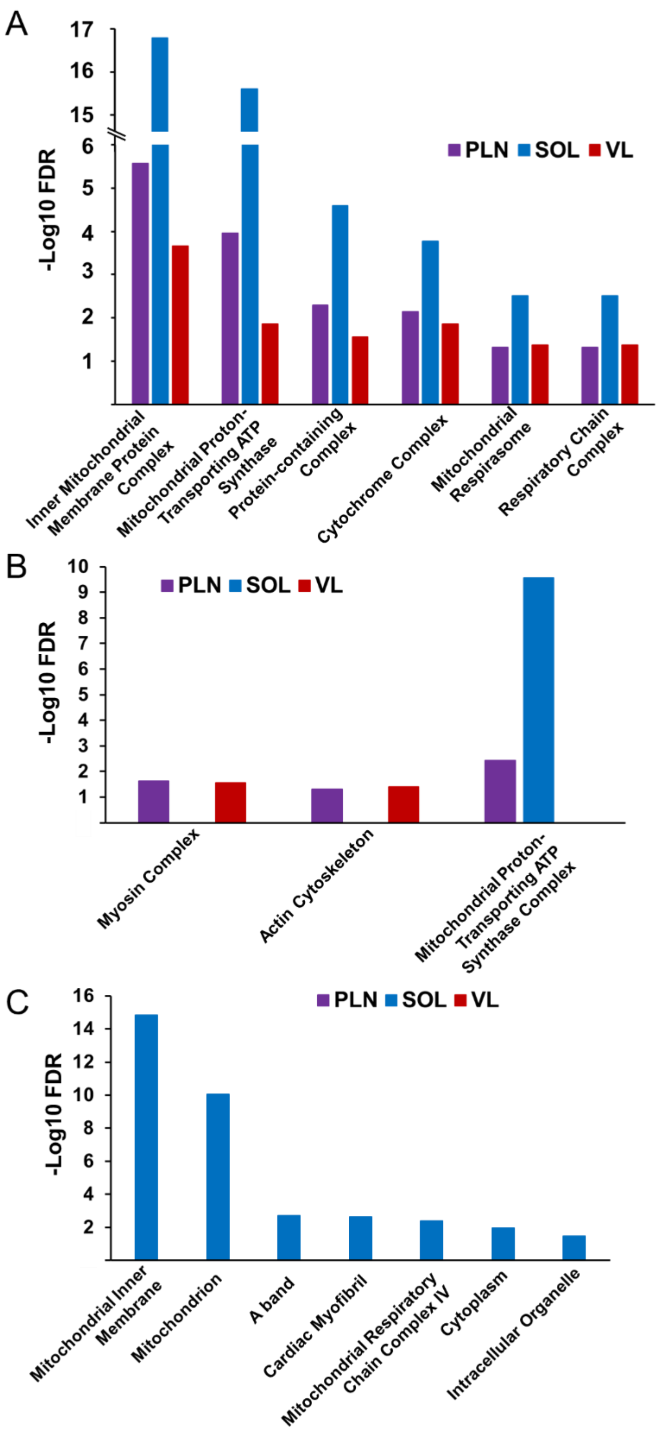


**Figure S34. Gene Ontology (GO) summary from VL, PLN, and SOL fiber samples using TopPIC search results.** A) Plot of identified cellular component GO terms which occur across all 3 fiber types; B) Plot of identified cellular component GO terms which occur across 2 fiber types; C) Plot of identified cellular component GO terms which occur in only 1 fiber type.
